# Mechanisms of antibiotic action shape the fitness landscapes of resistance mutations

**DOI:** 10.1101/2020.06.01.127571

**Authors:** Colin Hemez, Fabrizio Clarelli, Adam C. Palmer, Leonid Chindelevitch, Theodore Cohen, Pia Abel zur Wiesch

## Abstract

Antibiotic-resistant pathogens are a major public health threat. A deeper understanding of how an antibiotic’s mechanism of action influences the emergence of resistance would aid in the design of new drugs and help to preserve the effectiveness of existing ones. To this end, we developed a model that links bacterial population dynamics with antibiotic-target binding kinetics. Our approach allows us to derive mechanistic insights on drug activity from population-scale experimental data and to quantify the interplay between drug mechanism and resistance selection. We find that whether a drug acts as a bacteriostatic or bactericidal agent has little influence on resistance selection. We also show that heterogeneous drug-target binding within a population enables resistant bacteria to evolve fitness-improving secondary mutations even when drug doses remain above the resistant strain’s minimum inhibitory concentration. Our work suggests that antibiotic doses beyond this “secondary mutation selection window” could safeguard against the emergence of high-fitness resistant strains during treatment.

## 1. Introduction

The emergence and spread of antibiotic-resistant bacterial pathogens is an urgent global problem that threatens to undermine one of the most essential components of modern medicine [1]. Antibiotic resistance is also expensive, adding an average of US $1400 to the costs of treatment for each of the 2.8 million patients who become infected with a drug-resistant bacterium in the United States annually [2–4]. The scarcity of promising new antimicrobial drugs with novel mechanisms of action further exacerbates the challenges associated with managing the spread of drug resistance [5, 6]. Given the increasing incidence of resistant bacterial infections and the lack of new drugs on the horizon, clinicians, researchers, and global leaders must act to preserve the effectiveness of the world’s existing antibiotic drug arsenal [1].

Antibiotic treatment induces a strong selective pressure on bacterial populations to evolve resistance [7, 8]. Resistance mutations raise the minimum inhibitory concentration (MIC) of an antibiotic, the amount of drug needed to suppress the growth of a bacterial culture [9]. However, alleles that confer drug resistance also frequently carry fitness costs [10–12], predominantly because antibiotics target vital cellular functions (such as DNA replication and protein synthesis). Resistance mechanisms reduce the ability of a drug to disrupt its target, but do so at the expense of optimal physiological function [13].

With few exceptions [14], resistance-causing alleles induce physiological impairments in both drug-free and drug-containing environments, though resistant strains may only suffer a strict competitive disadvantage (i.e. a slower growth rate) against sensitive strains in drug-free conditions. A range of antibiotic concentrations therefore exists within which drug-resistant strains have a selective advantage over their drug-susceptible counterparts. Drugs dosed within this “resistance selection window” (also called the “mutant selection window”) favor the proliferation of drug-resistant subpopulations [15–17]. Recent advances in antimicrobial pharmacodynamics have leveraged resistance selection windows to design dosing strategies that minimize the selection of resistant pathogens without sacrificing treatment efficacy [17–19].

The existence of resistance mutations that confer physiological impairments in both drug-free and drug-containing environments implies that resistant strains face selective pressures to evolve secondary mutations that alleviate these impairments, and that these selective pressures exist even under continuous drug exposure [20, 21]. Secondary mutations can increase bacterial fitness (through faster growth rates) in the absence of drugs, or they can confer elevated levels of drug tolerance to preexisting resistant subpopulations (through attenuated drug-target interactions, faster growth rates in the presence of drugs, or both). In the case of increased bacterial fitness, secondary mutations enable drug-resistant mutants to compete against drug-susceptible strains in resource-limited, antibiotic-free environments [10, 22, 23], and are implicated in the spread of drug resistance across a wide range of timescales and clinical settings [24]. In the case of increased drug tolerance, secondary mutations can be the underlying cause of treatment failure [25, 26]. Elucidating the dynamics of secondary mutation emergence during treatment is thus crucial for managing the spread of resistance.

Since resistance mutations are frequently associated with fitness costs [11, 12] both *in vivo* [27] and *in vitro* [28], studies on the resistance selection window and on secondary adaptation have yielded valuable insights into the emergence of drug-resistant bacteria during treatment. However, the design of optimal resistance-mitigating drug dosing strategies remains challenging for two reasons. One obstacle is that bacteria may acquire resistance through a multitude of mechanisms that reduce antibiotic efficacy [29]. These molecular mechanisms may themselves influence the fitness landscape of resistance mutations (that is, the relationship between the fitness cost of resistance and the selective advantage conferred by the resistance mutation in drug-containing environments)[30]. A second challenge is that an antibiotic’s mechanism of action may affect the strength of selection for resistant strains over drug-susceptible strains during treatment. One important feature of an antibiotic’s cellular-level mechanism of action is whether the drug controls bacterial populations by increasing the rate of bacterial killing (i.e. bactericidal action) or by decreasing the rate of bacterial replication (i.e. bacteriostatic action). Clinicians and researchers alike have argued that these modes of antimicrobial action influence the dynamics of resistance selection [31, 32].

The design of resistance-mitigating antibiotic usage therefore depends on an understanding of how a drug’s mechanism of action, a pathogen’s mechanism of resistance, and the fitness landscape of resistance affect selection pressures during treatment. Tractable and quantitative strategies for systematically exploring all of these factors have so far been lacking. To address this gap, we developed a dynamical model that simulates the growth and death of bacterial populations under antibiotic exposure using molecular-scale descriptions of drug-target binding kinetics and cellular-scale descriptions of a drug’s mechanism of action. In our model, higher numbers of inactivated drug-target complexes within a cell lead to increases in antibiotic effect (either bacteriostatic, bactericidal, or a combination of the two). The relationship between drug-target inactivation and antibiotic effect can take the shape of a linear (i.e. gradual) or stepwise (i.e. sudden) function, as well as other intermediate forms (**Supplementary Figure S1**). The model enables us to estimate critical pharmacodynamic parameters from experimental datasets as effectively as with classical approaches [33], to simulate the fitness landscapes of resistance mutations against drugs with diverse mechanisms of action, and to quantify the probability of secondary mutation emergence within resistant subpopulations of bacteria during treatment.

The mathematical model described here is a linear case of nonlinear formulations we have reported previously to study the influence of drug-target binding kinetics on optimal antibiotic dosing [34]. Linearization results in a >10^2^-fold computational speed-up that enables us to robustly fit experimental kill-curve data and to simulate antibiotic dose-response relationships at high resolution. Our linear formulation also allows us to calculate an antibiotic’s MIC directly from experimentally measurable molecular parameters. We leverage the mathematical tractability and computational efficiency of the linear model to investigate the selective pressures that antibiotics with diverse mechanisms of action place on growing bacterial populations, a task that would be impractical with previous approaches.

We find that bacteria with resistance mechanisms that confer even modest reductions in drug-target binding affinity can incur strikingly high (80-99%) fitness costs while still maintaining higher drug tolerances than their susceptible counterparts, regardless of the antibiotic’s mechanism of action. We also find that drugs with stepwise effects on bacterial growth and death have narrower resistance selection windows than do drugs with linear effects. However, our model suggests that whether a drug acts primarily through bactericidal or bacteriostatic action has comparatively little influence on the strength of resistance selection during treatment. We further demonstrate that, even with aggressive treatment regimens, heterogeneous drug-target occupancy within a population enables fitness-impaired resistant strains to develop secondary mutations that can lead to treatment failure. Our work cautions that fitness costs may not limit the emergence of resistant strains that evolve through reductions in drug-target binding affinity. We propose the “secondary mutant selection window” as a novel pharmacodynamic characteristic of a drug that should be assessed alongside other classic parameters such as the MIC and the resistance selection window when designing robust resistance-mitigating antibiotic dosing strategies.

## 2. Results

### 2.1. A model that links bacterial population dynamics with molecular mechanisms of antibiotic action

We developed a linear dynamical model to describe the effect of a constant concentration of drug on the growth and death rates of a bacterial population (**Figure 1A**) (see **Methods**, *Model formulation and analysis* for a mathematical description of the model). We assume that each bacterial cell in the population carries an identical number *N* of intracellular proteins that the drug targets for inactivation. Drug molecules inactivate target proteins by binding to them with a rate *k_F_* and can dissociate from the target with a rate *k_R_*. The affinity *K_D_* of the drug is thus the ratio of off-rate to on-rate, *K_D_* = *k_R_*/*k_F_*. The model assumes that the growth and death rates of a bacterial cell depend on its drug-target occupancy (that is, the number of inactivated drug-target complexes it contains) [34, 35]. We denote drug-target occupancy with the index *i*, which ranges from 0 to *N*. Cells harboring successively larger numbers of inactivated drug-target complexes have successively faster death rates and/or slower growth rates, depending on the mechanism of action of the drug (see **Results,** *Classification of drug action*). We thus define the growth rate (*G*[*i*]) and death rate (*D*[*i*]) of each subpopulation as discrete monotonic functions of drug-target occupancy. In practice, *G*[*i*] and *D*[*i*] take the form of constrained logistic functions each controlled by a steepness and inflection point parameter, allowing us to define quasi-linear, quasi-stepwise, quasi-exponential, and sigmoid curves (**Supplementary Figure S1**).

**Figure 1.**
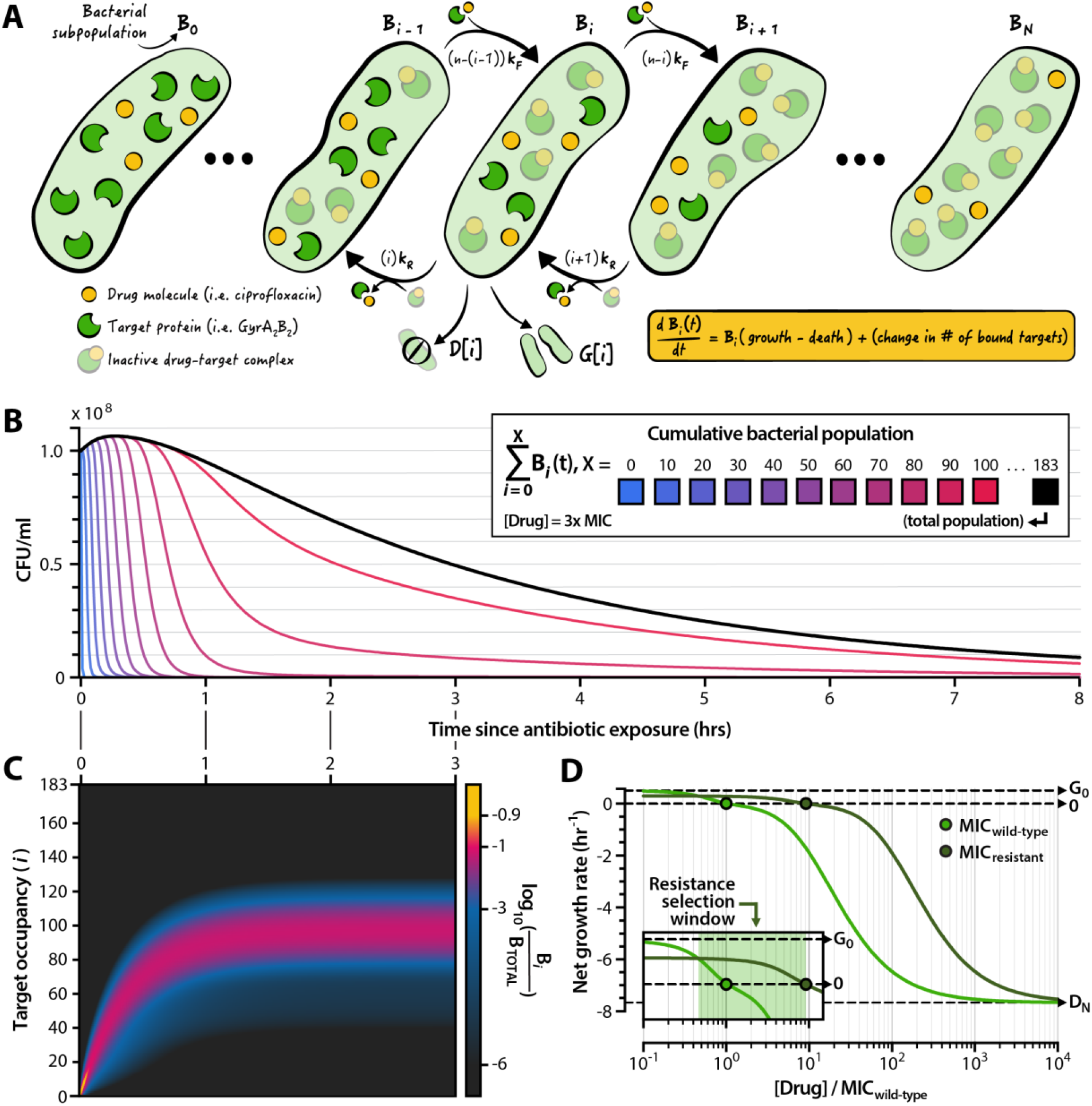
Features of a model that links bacterial population dynamics with the cellular mechanisms of antibiotic drug action. (**A**) Illustration of the model. We consider a population *B_i_* of bacterial cells harboring *i* inactive drug-target complexes. The change in the size of *B_i_* is a function of cellular growth and death rates (each of which is determined by the value of *i*, **Supplementary Figure S1**), and of the molecular kinetics of the drug binding and unbinding to its protein target. The total bacterial population is given by the sum *B_0_* + *B_1_* + … + *B_N_*_-1_ + *B_N_*, where *N* is the number of drug targets per cell. (**B**) Dynamics of a bacterial population exposed to a drug dose above the minimum inhibitory concentration (MIC). The black line represents the total bacterial population; shaded lines represent subpopulations with *x* and fewer inactivated drug-target complexes. Population dynamics as a function of drug concentration are shown in **Supplementary Figure S2**. (**C**) Proportion of the bacterial subpopulation *B_i_* as a share of total population for the first three hours of the curve shown in panel (B). (**D**) Pharmacodynamic curves derived from the model for a wild-type (light green) and drug-resistant (dark green) bacterial strain. The MIC is denoted as the drug concentration at which the net bacterial growth rate is zero. Inset: the resistance selection window (green shading) is given by the drug concentration range within which the drug-resistant strain exhibits a higher—but still positive— net growth rate compared to the wild-type strain. *G_0_* denotes the growth rate of the wild-type strain in the absence of antibiotic (i.e. the growth rate for subpopulation *B_0_*). *D_N_* denotes the maximum death rate of bacterial strains when all *N* cellular targets are inactivated (i.e. the death rate of subpopulation *B_N_*).

The model tracks the growth and death of all *N*+1 bacterial subpopulations, each denoted *B_i_*, over time (**Figure 1B**). Drug concentration determines the net growth rate of the entire bacterial population (**Supplementary Figure S2**). In the absence of drug, the population grows exponentially at a rate equal to the difference between the drug-free growth and death rates (*G_0_* and *D_o_*, respectively). When drug is present, the composition of bacterial subpopulations asymptotes towards a steady state after a transient phase during which drug molecules bind to their targets (**Figure 1C**). At steady state, the relative composition of bacterial subpopulations does not depend on the total size of the population.

We can calculate the MIC of a drug directly from model parameters (see **Methods**, *Calculation of the minimum inhibitory concentration*), and we can simulate clinically observed drug resistance mutations by modulating the parameters of the model that influence the value of the MIC. Changes in the binding kinetics of the drug (i.e. *k_F_* and *k_R_*) simulate target modification mutations that decrease the affinity of an antibiotic molecule to a cellular protein [36–38]. Changes to the value of *N* represent changes in the number of protein targets per cell, equivalent to target up- or downregulation [39–41]. We assume that fitness costs associated with resistance alleles take the form of reduced growth rates, and we simulate this cost by reducing the drug-free growth rate of resistant strains by a factor *c_R_* such that the maximum growth rate of a resistant strain (*G_0,RES_*) relative to that of a wild-type strain is *G_0,RES_* = *G_0_*(1–*c_R_*). When *c_R_* ranges from 0 (no cost) to 1 (no growth), the resistant strain exhibits a slower growth rate relative to that of the wild-type. If *c_R_* is negative, the resistant strain exhibits a faster drug-free growth rate than does the wild-type strain, as has been observed in rare cases with some fluoroquinolone-resistant *Escherichia coli* isolates [42]. The model also enables us to generate pharmacodynamic curves by calculating the net growth rates of simulated bacterial populations over a range of drug concentrations (**Figure 1D**). The resistance selection window constitutes the range of drug concentrations over which a drug-resistant mutant strain has a higher but strictly positive net growth rate relative to that of its wild-type counterpart (**Figure 1D**, inset).

### 2.2. Inferring cellular mechanisms of antibiotic action from population-scale data

To test the utility of our biochemical model for gaining cellular-scale insights into antimicrobial drug mechanisms from population-scale experiments, we calibrated our model to a family of experimental time-kill curves of the gram-negative bacterium *Escherichia coli* challenged to ciprofloxacin, a fluoroquinolone first brought to market in 1987. Ciprofloxacin has two known molecular targets in bacteria, both of which are heterotetrameric type-II topoisomerases: the DNA gyrase complex (GyrA_2_B_2_) and DNA topoisomerase IV (ParC_2_E_2_). However, ciprofloxacin preferentially binds to the GyrA_2_B_2_ complex in gram-negative bacteria [43]. We used a mass-spectrometry based estimate for the number of GyrA_2_B_2_ complexes per *E. coli* cell (*N* ∼ 183) as the number of drug targets within each bacterium [44].

We implemented an adaptive simulated annealing algorithm to calibrate the parameters of our model to an experimental dataset of ciprofloxacin time-kill curves (**Methods,** *Model calibration via simulated annealing*). We performed 249 independent parameterizations using the algorithm and selected the parameter set that yielded the lowest objective function value (**Figure 2A, Table 1, Supplementary Figure S3**). Bacterial persistence [45, 46] likely plays a role in the slower-than-expected population decline that we observe experimentally at high drug concentrations. At antibiotic doses below those that elicit persistence, the calibrated model accurately recapitulates the pharmacodynamic curve derived from experimental data (**Supplementary Figure S4**).

**Figure 2.**
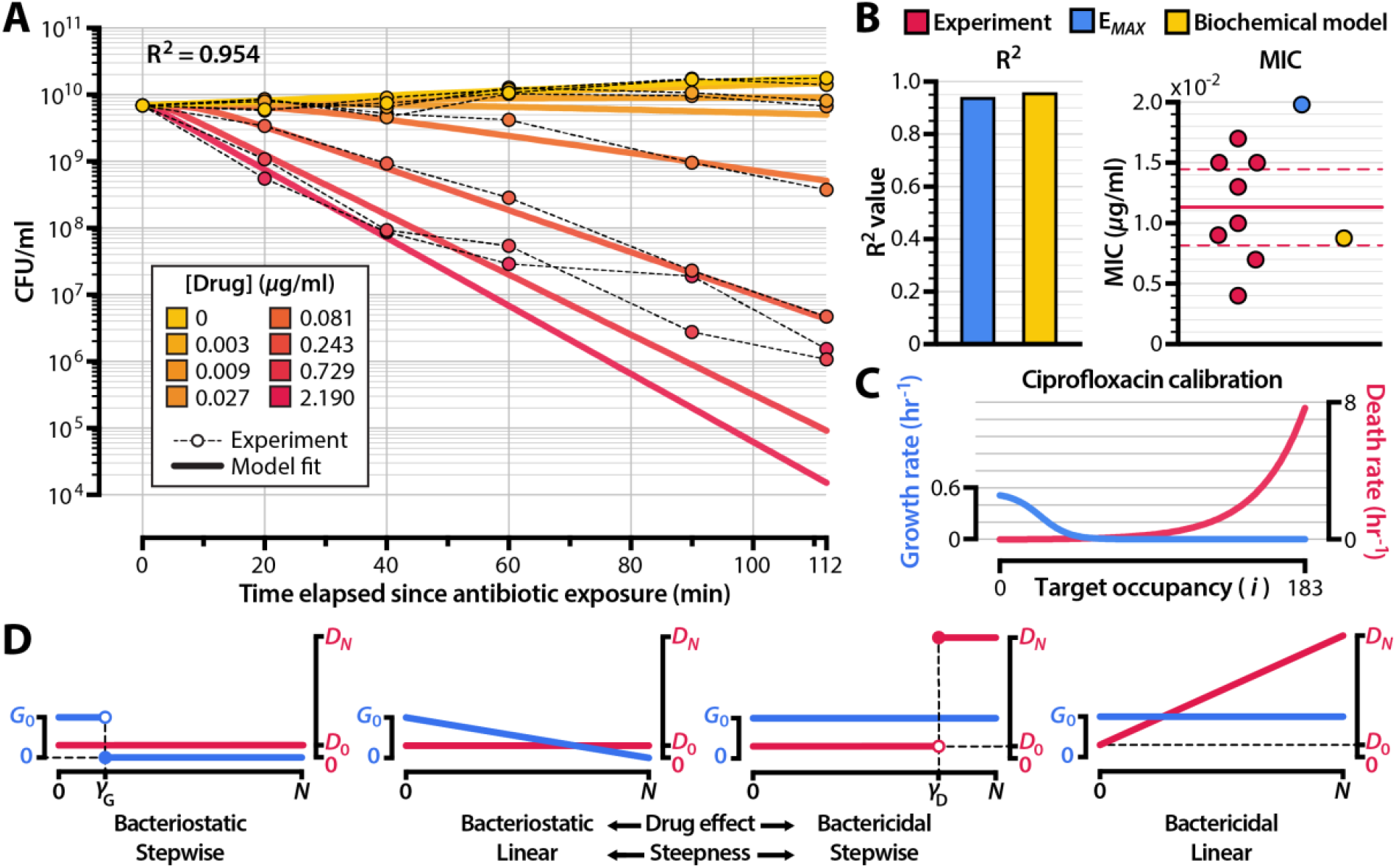
Calibrating the model to experimental data reveals underlying mechanisms of drug action. (**A**) Comparison between calibrated biochemical model (solid lines) and experimental data (shaded points). The experimental data (**Supporting Data File S1**) represent time-kill curves of *Escherichia coli* exposed to ciprofloxacin. A summary of all independent model calibrations is shown in **Supplementary Figure S3**. (**B**) Comparison of the calibrated biochemical model with the *E_MAX_* pharmacodynamic model [33]. We fit the *E_MAX_* model to the same experimental dataset shown in panel (A) and compared Pearson correlation coefficients (R^2^) and MICs. Red points in the MIC panel denote experimentally-measured ciprofloxacin MICs for *E. coli* strains isolated prior to the widespread emergence of quinolone resistance (**Supporting Data File S2**). The solid horizontal line represents the mean of experimental measurements, and the dashed lines indicate the 95% confidence interval. A comparison of the pharmacodynamic curves obtained from the models is shown in **Supplementary Figure S4**. (**C**) Cellular growth and death rates as a function of ciprofloxacin-GyrA_2_B_2_ complex number (*i*) for the model calibrated to the experimental data shown in panel (A). (**D**) Four extreme schemes of drug action resulting from two characteristics (activity and steepness) of a drug’s effect on growth and death rates as a function of drug-target occupancy. **Supplementary Figure S5** shows the simulated bacterial kill curves for these schemes at 4x MIC. Model fits for drug-free growth rate (*G_0_*) and drug-saturated death rate (*D_N_*) are shown in **Supplementary Figure S6**.

**Table 1.** Model parameters. We obtained the values of *k_F_*, *k_R_*, *α_G_*, *α_D_*, *γ_G_*, *γ_D_*, and *B_0_* by calibrating the model to experimental data (**Figure 2**). We inferred antibiotic-free growth rate and antibiotic-saturated death rate (*G_0_* and *D_N_*) by fitting an exponential curve to ciprofloxacin kill curves using 0 and 2.19 µg/ml of drug, respectively (**Supplementary Figure S6**). We use a constrained logistic function to model the growth and death rates of bacterial cells as a function of bound target number, where *α* controls the steepness of the logistic function and *γ* controls the inflection point of the logistic function (**Supplementary Figure S1**). Parameters not obtained from model calibrations to experimental data were retrieved from the literature. For the bacterial death rate in the absence of drug (*D_0_*), we used the mean of death rates reported in Wang et al., 2010.

| Model parameters |  |  |  |  |
| --- | --- | --- | --- | --- |
| Name | Description | Value | Units | Source |
| $N$ | Number of target proteins per cell (i.e. GyrA<sub>2</sub>B<sub>2</sub> copy number) | 183 | cell<sup>-1</sup> | [44] |
| $G_0$ | Bacterial growth rate in the absence of drug | 0.526 | hr<sup>-1</sup> | Model calibration |
| $D_0$ | Bacterial death rate in the absence of drug | 5.40 x 10<sup>-3</sup> | hr<sup>-1</sup> | [47] |
| $D_N$ | Bacterial death rate in saturating concentrations of drug | 7.53 | hr<sup>-1</sup> | Model calibration |
| $k_F$ | Drug-target binding rate | 5.23 x 10<sup>3</sup> | M<sup>-1</sup> sec<sup>-1</sup> | Model calibration |
| $k_R$ | Drug-target unbinding rate | 3.17 x 10<sup>-4</sup> | sec<sup>-1</sup> | Model calibration |
| $\alpha_G$ | Steepness of growth rate function $G[i]$ | 16.8 | # drug-target complexes<sup>-1</sup> | Model calibration |
| $\alpha_D$ | Steepness of death rate function $D[i]$ | 7.29 | # drug-target complexes<sup>-1</sup> | Model calibration |
| $\gamma_G$ | Inflection point of growth rate function $G[i]$ | 24.9 | # drug-target complexes | Model calibration |
| $\gamma_D$ | Inflection point of death rate function $D[i]$ | 359 | # drug-target complexes | Model calibration |
| $B_0$ | Initial size of bacterial population at the start of drug treatment | 6.88 x 10<sup>9</sup> | cell ml<sup>-1</sup> | Model calibration |
| $\mu_R$ | Mutation rate for drug resistance emergence | 2.00 x 10<sup>-7</sup> | cell<sup>-1</sup> division<sup>-1</sup> | [48, 49] |
| $\mu_C$ | Mutation rate for emergence of secondary mutations in resistant strains | 2.00 x 10<sup>-6</sup> | cell<sup>-1</sup> division<sup>-1</sup> | [48, 49] |
| $c_R$ | Cost of resistance mutation, such that the antibiotic-free growth rate of a resistant mutant is $G_0(1 - c_R)$ | 0.25 | Non-dimensional | [50] |

We compared our biochemical model’s ability to capture critical pharmacodynamic characteristics of a drug against that of an *E_MAX_* model [33]. The *E_MAX_* approach describes net bacterial growth rate directly as a function of drug concentration and does not accommodate molecular descriptions of drug-target interactions. Such models have been used extensively to estimate pharmacodynamic parameters, to design drug dosing regimens, and to predict the strength of resistance selection at nonzero drug concentrations. Our formulation delivers performance comparable to that of the *E_MAX_* model for fitting experimental time-kill curves (**Figure 2B**, left panel) and more accurately estimates MIC (which we calculated to be 8.9 x 10^-3^ *µ*g/ml for ciprofloxacin) from these data (**Figure 2B**, right panel). This demonstrates the validity of our approach for deriving pharmacodynamic insights similar to what an *E_MAX_* model provides.

Our model furthermore offers capabilities that the *E_MAX_* approach lacks, including the ability to estimate molecular kinetic parameters of drug-target binding from population-scale data. To test the robustness of these estimates, we analyzed the *K_D_* values for ciprofloxacin binding to *E. coli* GyrA_2_B_2_ generated for the 249 independent parameterizations described above. As our fitting method is stochastic, not all model calibrations reach local minima. However, the best 90% of all calibrations (that is, the 224 fits with the lowest objective function values) consistently converged upon a narrow range of affinity values (95% confidence interval: 7.2 x 10^-8^ to 1.6 x 10^-7^ M) (**Supporting Data File S3**). Our estimates lie within the range of *K_D_* values of ciprofloxacin for *E. coli* GyrA_2_B_2_ reported from experimental measurements, which span from 3.2 x 10^-8^ to 3.0 x 10^-6^ M [51–54].

### 2.3. Classification of antibiotic action

Another unique feature of our approach is the ability to describe bacterial growth and death rates as a function of drug-target occupancy. For ciprofloxacin, the calibrated model predicts three regimes of bacterial subpopulation dynamics in relation to GyrA_2_B_2_ inactivation: a growth regime in which bacterial replication dominates among subpopulations with low numbers of inactivated targets, a stalling regime for intermediate numbers of drug-target complexes in which neither growth nor death is appreciable, and a killing regime at high numbers of inactivated targets in which bacterial death increases quasi-exponentially (**Figure 2C**). The forms of G[*i*] and D[*i*] that we obtain here suggest that ciprofloxacin has a multimodal mechanism of action, a result consistent with prior experimental studies [43, 55, 56] and with more complex nonlinear modeling approaches [34]. The drug stalls cellular replication at intermediate target occupancies and induces killing only at higher doses. Like many antibiotics, ciprofloxacin thus exhibits both bactericidal and bacteriostatic effects on microbial populations [56, 57]. Our biochemical model represents this explicitly.

Most drugs nonetheless demonstrate a greater degree of bactericidal or bacteriostatic activity at clinically relevant doses [58], and we hypothesized that the ability of a drug to stall growth or to accelerate death may affect the selection for resistant strains and the emergence of secondary mutations. We also suspected that the relationship between drug-target occupancy and antibiotic effect—reflected in the steepness of the *G*[*i*] and *D*[*i*] functions—could further shape the dynamics of resistance selection.

These two characteristics (bactericidal versus bacteriostatic activity and drug effect steepness) represent two general dimensions along which a drug’s mechanism of action can affect the growth and death of bacterial populations. Four extreme cases of drug action thus exist (**Figure 2D**). In the case of a purely bacteriostatic antibiotic, death rates are a constant function of inactivated drug-target complex number (that is, D[*i*] = *D_0_* for all values of *i*). For a purely bactericidal antibiotic, the growth rate of all bacterial subpopulations remains constant (G[*i*] = *G_0_* for all values of *i*). The steepness of the drug effect is reflected in the form of the function D[*i*] for bactericidal antibiotics and G[*i*] for bacteriostatic antibiotics (**Supplementary Figure S1**). We defined linear and stepwise onset of action as our two extremes, as other monotonic forms are intermediate cases of these curves.

### 2.4. The opposing effects of increased drug resistance and decreased cellular fitness

Mutations that confer resistance against antibiotics often come at the cost of reduced growth rates compared to those of drug-susceptible strains [10, 11]. The balance of replication and death determines bacterial net growth both in the absence and in the presence of antibiotics, and very high fitness costs associated with resistance can prevent bacterial viability at any drug concentration [59]. We sought to investigate the quantitative basis for the trade-off between drug resistance and cellular growth and to investigate how the drug mechanisms defined above influence the range of permissible fitness costs that a drug-resistant mutant can incur while still maintaining a drug susceptibility that is lower than that of a wild-type strain. In the simplest case of the model, where the number of target molecules per cell is 1, the expression for the MIC captures the opposing effects of drug resistance and cellular growth (see **Methods**, *Calculation of minimum inhibitory concentration* for derivation):

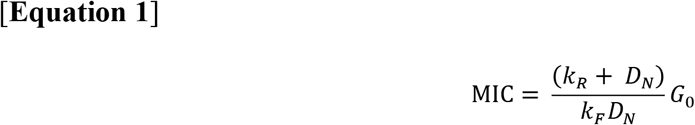

The MIC increases with reductions of the on-rate kinetics of drug-target binding (*k_F_*) and with increases in the off-rate kinetics of drug-target binding (*k_R_*), but decreases with fitness costs that manifest as reductions in the drug-free growth rate (*G_0_*). These proportionalities hold for any number *N* of drug targets.

We modeled the opposing effects of biochemical changes that reduce drug susceptibility (i.e. altered drug-target binding kinetics or target upregulation) and the fitness costs of these biochemical changes. We considered a set of five antibiotics with an identical protein target and identical molecular kinetic parameters (that is, the target number *N*, the drug-target on-rate *k_F_*, and the drug-target off-rate *k_R_* are constant for the wild-type strain) (**Supplementary File S2**, **Supplementary Figure S5**). One antibiotic in the set features growth and death dynamics derived from the model calibration to ciprofloxacin time-kill curve data (**Figure 2C**). The other four antibiotics are hypothetical and feature growth and death dynamics that represent four extremes of antibiotic action (**Figure 2D**). We simulated mutant strains of *E. coli* that acquire drug resistance phenotypes either through changes in the molecular kinetics of drug binding (*k_F_* or *k_R_*) or by increasing the copy number *N* of the drug’s cellular protein target. Each of these resistance mechanisms has been observed in clinical isolates of drug-resistant, gram-negative bacteria [11, 29, 60]. We then simulated fitness costs associated with the resistance mutation and calculated the mutant strain’s MIC relative to that of the wild-type strain.

For resistance acquired through changes in the kinetics of drug-target binding (*k_F_* and *k_R_*), we found that mutants can tolerate strikingly high (80-99%) fitness costs while still maintaining an MIC that is greater than that of the drug-susceptible wild-type (**Figure 3**, top and middle rows). This permissibility of fitness costs exists for all five of the drug mechanisms we simulated, although drugs that act with linear effects (Bacteriostatic/Linear and Bactericidal/Linear) have a narrower range of permissible fitness costs than do drugs that act with stepwise effects. For all drug mechanisms, mutant strains make larger gains in MIC by decreasing the on-rate kinetics of drug-target binding (*k_F_*) than they do by increasing the off-rate kinetics of drug-target binding (*k_R_*) by the same amount (**Supplementary Figure S7**). That is, mutations that lead to the same change in drug-target affinity (as quantified by the dissociation constant *K_D_* = *k_R_*/*k_F_*) through different changes in the on- and off-rate binding kinetics do not necessarily have the same range of permissible fitness costs. This has biological significance— limiting the opportunity for a drug to bind to its target, thereby preventing the drug from actuating its effects on cellular growth and death, should lead to lower drug susceptibilities than would accelerating the rate at which an already-formed drug-target complex disassociates. The difference in the fitness effects of mutations that modify *k_F_* and *k_R_* is especially pronounced for bactericidal drugs that elicit linear increases in cellular death (Bactericidal/Linear).

**Figure 3.**
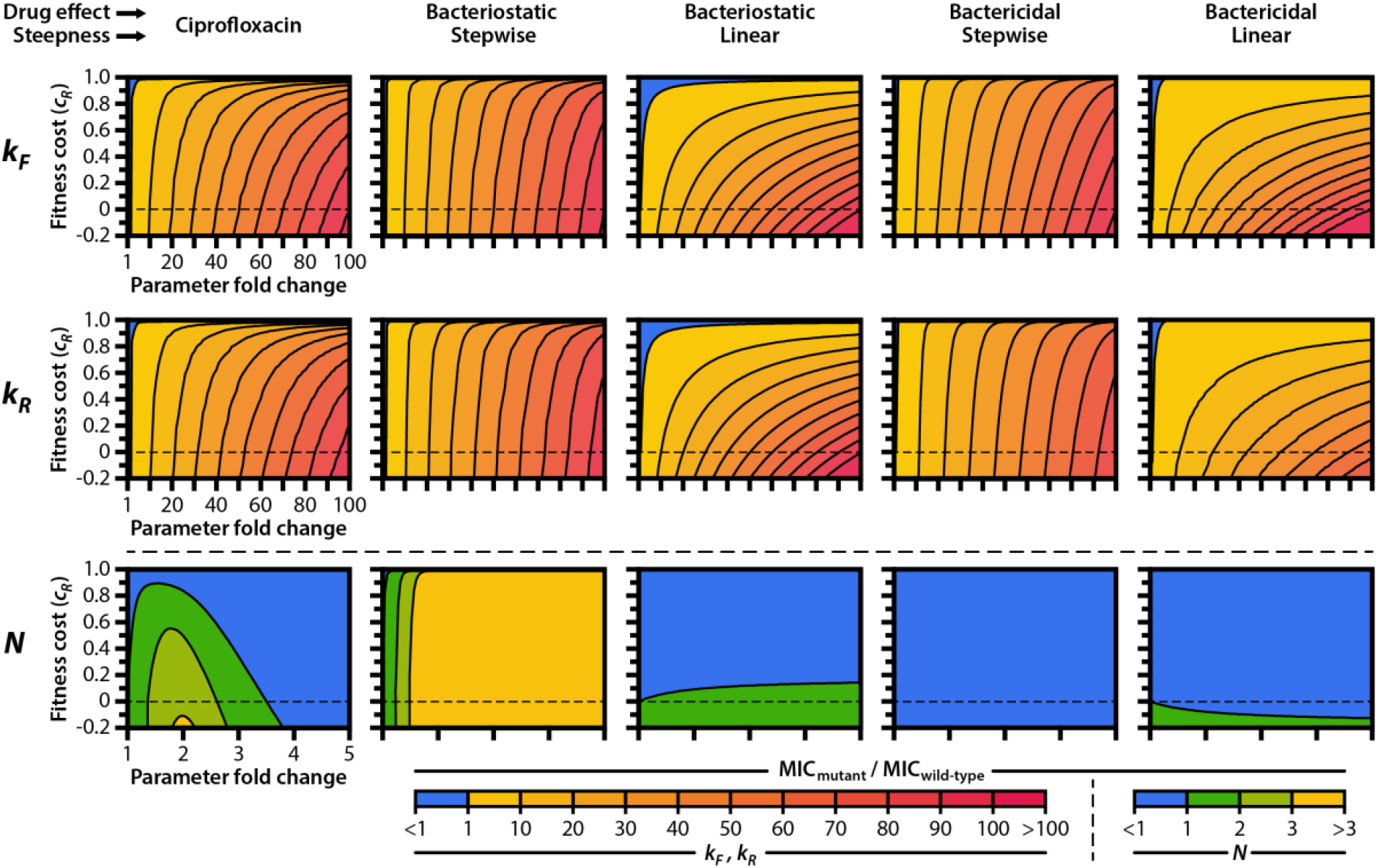
Drug mechanism influences the fitness landscapes of resistance mutations. We calculated the MIC, expressed as a fold-change relative to the MIC of the wild-type, for mutant strains carrying (top row) drug targets with reduced binding kinetics (*k_F_*), (middle row) drug targets with accelerated unbinding kinetics (*k_R_*), or (bottom row) increased numbers of drug target molecules (*N*). Mutant strains also carry fitness costs, expressed as a fractional reduction in drug-free growth rate relative to wild-type. Cost-free MIC as a function of *k_F_* and *k_R_* for all mechanisms of action are shown in **Supplementary Figure S7**. When modulating the number of drug target molecules *N* (bottom row), we assumed that cells require a fixed number of active protein targets to grow at a normal rate and that cellular killing is induced when a fixed number of inactive drug-target complexes form within a cell. Thus, the inflection point for the growth rate function (*γ_G_*) changes concomitantly with *N* such that *N*-*γ_G_* remains constant, while the inflection point for the death rate function (*γ_D_*) remains constant (see **Supplementary Figure S1** for illustrations of the effects of *γ_G_* and *γ_D_* on bacterial growth and death rates).

Ciprofloxacin exhibits a bactericidal effect by permitting GyrA_2_B_2_-mediated cleavage of DNA but preventing DNA re-ligation, resulting in widespread and eventually insurmountable chromosome fragmentation [43, 61]. When simulating the overexpression of target proteins in resistant cells (**Figure 3**, bottom row) we therefore assumed that bacterial killing is induced when a fixed number of inactivated drug-target molecules form within a cell (that is, we assume a toxicity threshold whereby *γ_D_* remains constant with changing *N*). Conversely, we assumed that a resistant cell requires a fixed number of active, non-complexed target proteins in order to maintain its maximum growth rate (that is, a survival threshold). *γ_G_* thus changes in step with *N* such that *N*-*γ_G_* remains constant. We made these same assumptions for the four hypothetical antibiotics.

We found that target overexpression has a diversity of effects on resistance that depend on the mechanism of action of the drug. For ciprofloxacin and its multimodal effects on growth and death, small increases in target number can lead to modest increases in MIC, even when the resistant cell faces large fitness costs as a result of GyrA_2_B_2_ overexpression. However, larger increases in target number lead to reductions in MIC. This result is consistent with experimental studies on target amplification, in which the overexpression of *gyrAB* in *E. coli* resulted in increased susceptibility to ciprofloxacin [40]. Target overexpression leads to substantial gains in resistance against bacteriostatic drugs that exhibit stepwise effects, even at very high fitness costs. The effect of target overexpression on drug resistance is negligible for bactericidal drugs and for bacteriostatic drugs with a linear effect on growth stalling.

### 2.5. Drug mechanism shapes the resistance selection window

To understand how a drug’s mechanism of action affects the propensity to select for resistance during treatment, we simulated the pharmacodynamics of wild-type and drug-resistant strains challenged to each of the five drugs in the set outlined above. MICs for clinical isolates of ciprofloxacin-resistant *E. coli* strains with single point mutations in GyrA, which may reduce the affinity of ciprofloxacin to GyrA_2_B_2_, range from 10 to 16 times greater than the MIC of a drug-susceptible wild-type [36, 60, 62, 63]. Data on the fitness costs associated with mutant GyrA-mediated ciprofloxacin resistance in *E. coli* are sparse, but studies of rifampicin-resistant clinical isolates of *Mycobacterium tuberculosis* with point mutations in the *rpoB* gene have suggested that a 20-30% reduction in growth rate is approximately the maximum fitness cost that drug-resistant mutants can incur before facing extinction in competitive drug-free environments [50]. To model drug-resistant strains, we therefore scaled *k_F_* and *k_R_* such that the MIC of the resistant strain is 12 times that of its drug-susceptible counterpart given a 25% fitness cost (*c_R_* = 0.25) (**Figure 4A**).

**Figure 4.**
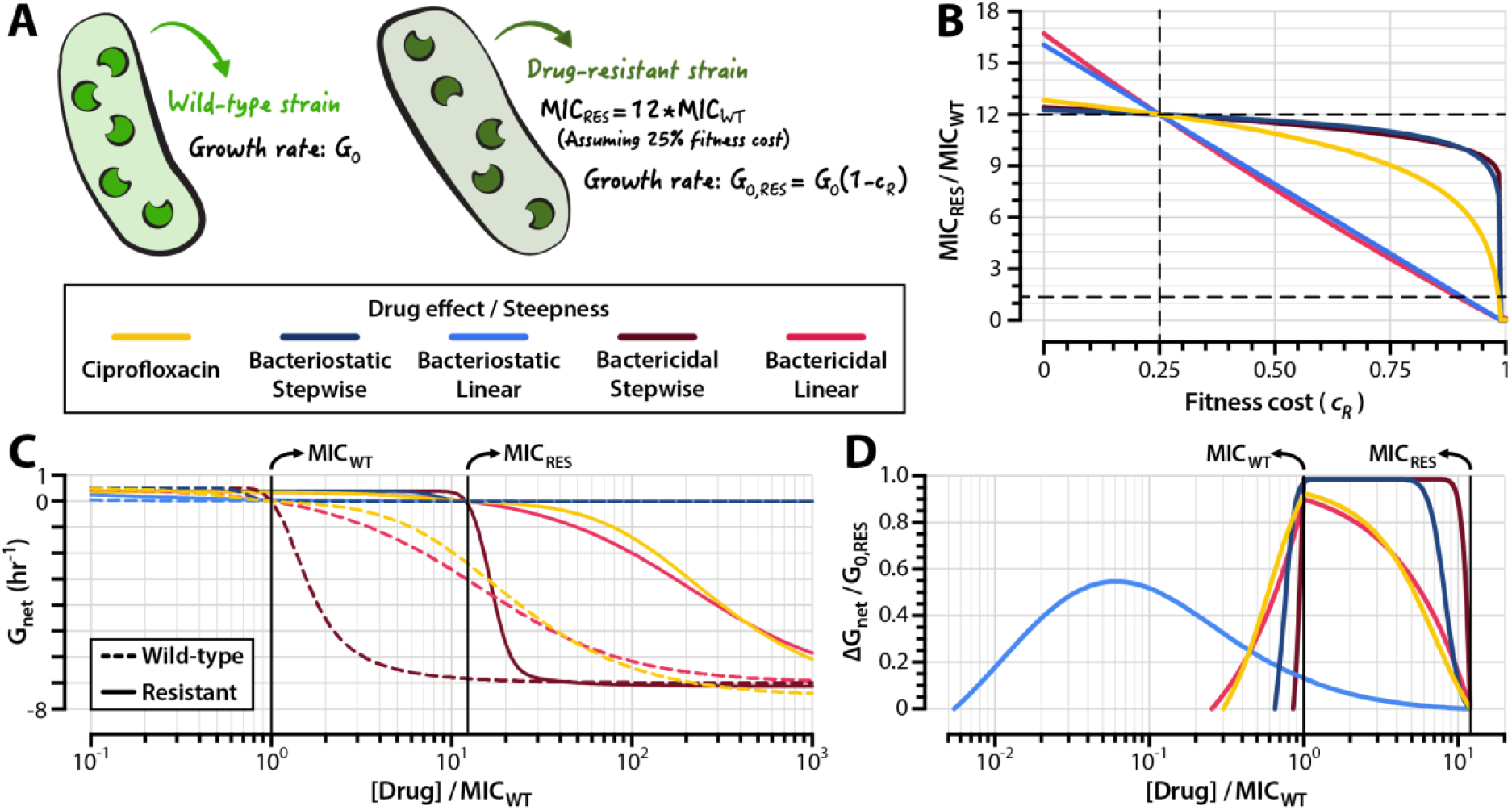
The propensity to select for resistant mutants depends on drug mechanism. (**A**) We modeled wild-type strains using the parameters obtained from the calibration detailed in Figure 2. (**B**) Relationship between MICs of resistant strains (expressed as multiples of MIC_WT_) and fitness cost of resistance. Horizontal dashed lines indicate the MICs of the wild-type and resistant strains described in panel (A); the vertical dashed line indicates the fitness cost at which all resistant strains have the same fold-increase in MIC relative to that of wild-type (*c_R_* = 0.25). (**C**) Pharmacodynamic curves for the wild-type and resistant strains described in panel (A). (**D**) Resistance selection windows for drug-resistant strains. The fitness advantage of resistant strains over wild-type strains is shown within the drug concentration range in which the resistant strain has a positive net growth rate that is larger than that of the wild-type. The fitness advantage is expressed as a proportion of the resistant strain’s growth rate in the absence of drug (*G_0,RES_*). **Supplementary Figure S8** illustrates the relationship between the size of the resistance selection window and the steepness of a drug’s pharmacodynamic curve.

A nearly linear relationship exists between drug resistance and fitness cost for strains resistant to drugs with a linear effect on growth or death (**Figure 4B**, Bacteriostatic/Linear and Bactericidal/Linear). By contrast, drugs with stepwise effects on growth and killing (Bacteriostatic/Stepwise and Bactericidal/Stepwise) exhibit only modest reductions in MIC until they incur very high (>90%) fitness costs. We determined resistance selection windows for strains resistant to the five drugs in our set by simulating pharmacodynamic curves for wild-type and resistant strains (**Figure 4C**). To quantify the magnitudes of selection for resistant strains, we calculated the difference in net growth rates between wild-type and susceptible strains over the concentration range that defines the resistance selection window for each drug (**Figure 4D**). For linear-effect bacteriostatic drugs (Bacteriostatic/Linear), we found that the resistance selection window begins at drug concentrations as low as 200x below the MIC of the susceptible strain. Drugs with stepwise effects on growth or killing (Bacteriostatic/Stepwise and Bactericidal/Stepwise) have narrower resistance selection windows than their counterparts with more linear activity profiles.

Consistent with prior studies on the pharmacodynamic profiles of antimicrobial agents [17, 19, 64], we find that the size of the resistance selection window is associated with the steepness of a drug’s pharmacodynamic curve. Given a cellular effect (i.e. bacteriostatic or bactericidal), drugs with steeper pharmacodynamic curves tend to have narrower selection windows (**Supplementary Figure S8**). However, we also find that strains resistant to drugs with narrower resistance selection windows have higher net growth rates within the resistance selection regime than do strains resistant to drugs with wider resistance selection windows (**Figure 4D**). This finding has clear clinical significance: drugs with steeper pharmacodynamic profiles feature relatively small concentration ranges that select for resistance, but the negative consequences of dosing within the resistance selection window are higher for these drugs.

### 2.6. The secondary mutant selection window is narrower for antibiotics with stepwise effects on growth and death

The genotypic space for mutations that confer resistance to antibiotics by modifying the binding kinetics of a drug to its target, such as those described in **Figure 4**, is typically highly constrained [22, 65]. Accordingly, a return to a drug-susceptible state requires reversion of the specific genetic changes that conferred resistance in a bacterial population. In contrast to resistance reversion, secondary mutation accumulation can involve a wider range of genetic changes throughout the cell’s metabolic network. Therefore, the probability that a bacterial population evolves secondary mutations that compensate for the fitness costs of a resistance mutation is often higher than the probability that a bacterial population will revert to susceptibility in drug-free environments [20, 66]. During treatment, resistant bacterial populations may also accumulate secondary mutations that further raise MIC. In order to understand how drug mechanism influences such secondary adaptation, we simulated the emergence of secondary mutants from drug-resistant subpopulations of a bacterial population faced with antibiotic challenge (**Supplementary Figure S9**; **Methods**, *Simulating the emergence of secondary mutations*).

The probability of secondary mutation emergence is substantially higher for drugs with linear effects on cellular growth and death than it is for drugs with stepwise effects (**Figure 5A**). This holds true for both bactericidal and bacteriostatic agents. Counterintuitively, then, the suppression of secondary mutation emergence is not necessarily guaranteed by rapid killing as suggested by earlier studies [67]. Likewise, rapid attenuation of cell division does not halt the emergence of secondary mutations. We studied the basis for this result by investigating the steady-state target occupancy distributions of cells under antibiotic exposure. By accounting for the kinetics of drug-target binding, our biochemical model shows that target occupancy among cells follows a distribution and is not a single value even in otherwise clonal bacterial subpopulations (**Figure 5B**). This results in heterogeneous replication rates within the drug-resistant subpopulation (**Supplementary Figure S10**) that allow some bacteria to mutate. Classical population-dynamic models of antibiotic action [33, 67], which assume that a drug affects the net growth rate of all cells equally, overlook this phenomenon.

**Figure 5.**
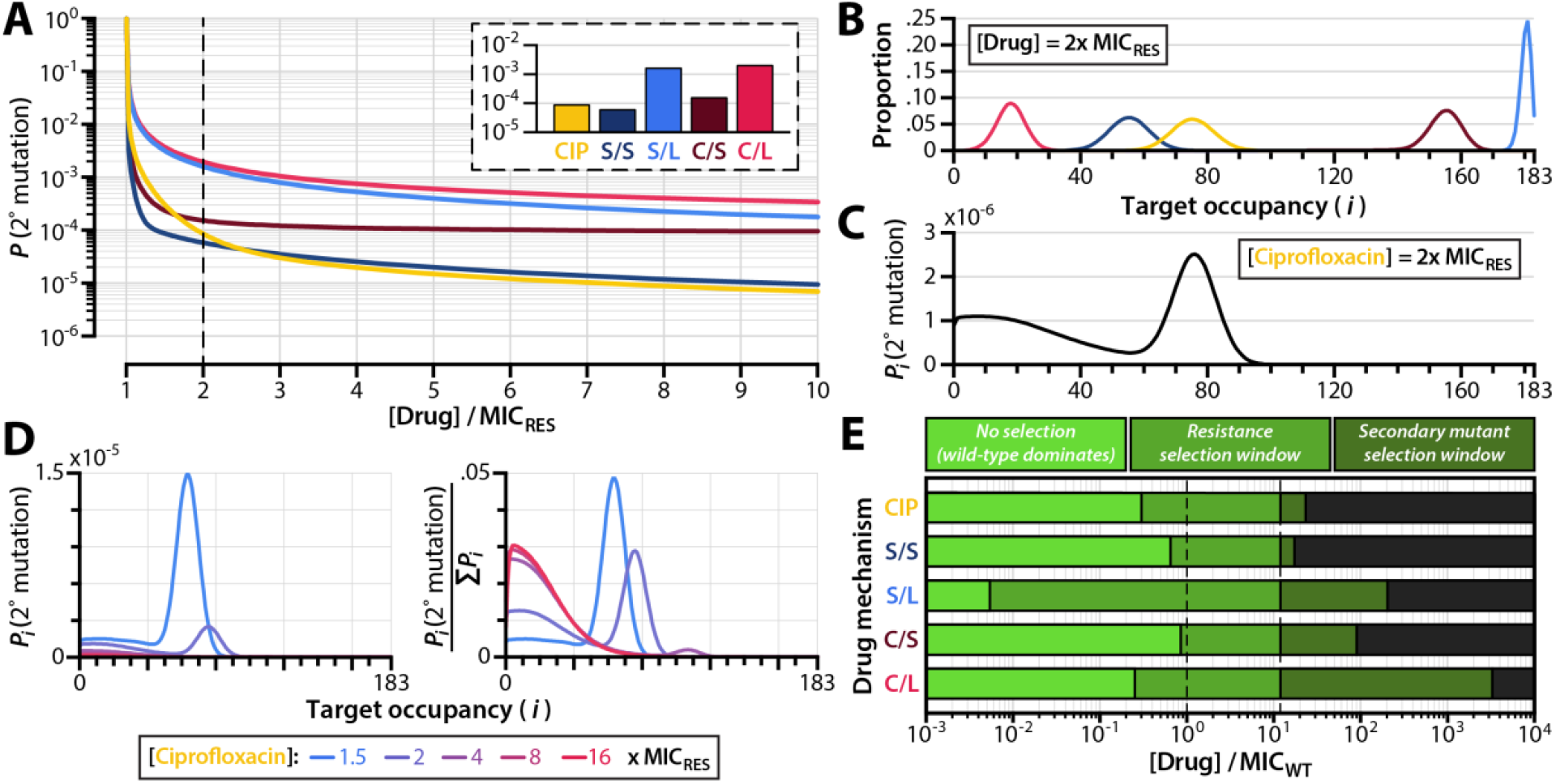
Emergence of secondary mutations among resistant subpopulations of infecting bacteria. (**A**) Probability of a drug-resistant strain with secondary mutations emerging from an infecting bacterial population before the infection is cleared (i.e. before the total bacterial population decreases to less than 1, **Supplementary Figure S9**). The initial population size for this simulation is 10^9^ cells. Inset shows probabilities of secondary mutation emergence before infection clearance when the drug concentration used is 2x MIC_RES_. (**B**) Frequency distributions of inactive drug-target complexes for drug-resistant subpopulations undergoing steady-state exponential decline at 2x MIC_RES_. Growth and death rate distributions for these populations are shown in **Supplementary Figure S10**. (**C**) Probability of secondary mutant emergence from bacterial subpopulations with *i* inactivated drug-target complexes, shown for ciprofloxacin dosed at 2x MIC_RES_. (**D**) Probability of secondary mutant emergence from bacterial subpopulations as a function of drug dose, shown for ciprofloxacin dosed at 2x MIC_RES_. Probabilities are shown as absolute values (left panel) and as values normalized to the total probability of compensation for the entire bacterial population over the course of treatment (right panel). (**E**) Resistance and secondary mutant selection windows for different drug action mechanisms. The resistance selection window (middle green) is defined as the drug concentration range over which a drug-resistant strain has a growth advantage over wild-type. The secondary mutant selection window (dark green) is defined as the drug concentration range over which the probability of a resistant strain with secondary mutations emerging before infection clearance exceeds 10^-4^ (see **Supplementary Figure S11** and **Methods**, *Simulating the emergence of secondary mutations*). Dashed lines indicate the MICs of the wild-type and resistant strains. CIP: ciprofloxacin; S/S: bacteriostatic/stepwise effect; S/L: bacteriostatic/linear effect; C/S: bactericidal/stepwise effect; C/L: bactericidal/linear effect; MIC_WT_: MIC of the wild-type strain; MIC_RES_: MIC of the resistant strain.

For ciprofloxacin doses only slightly above the MIC of the resistant strain ([Drug] = 2x MIC_RES_), we found that secondary mutations are most likely to emerge once the bacterial population has reached a steady-state target occupancy distribution (**Figure 5C**). A considerable probability of secondary mutation emergence nonetheless exists among bacterial subpopulations with low numbers of inactivated drug-target complexes. These low-occupancy subpopulations have faster growth rates and thus higher mutation rates. They are also present in very large numbers during the initial stages of treatment, when drug molecules are binding to their cellular targets and before the overall population begins to decline (**Figure 1C**). We found that drug concentration influences the likelihood of a secondary mutant arising from a steady-state or a low-occupancy subpopulation (**Figure 5D**). While the overall probability of secondary mutation emergence decreases with higher drug dose (**Figure 5D**, left panel), the relative probability that a secondary mutation arises from a low-occupancy population is greater for higher drug doses (**Figure 5D**, right panel). This implies that secondary mutations are more likely to emerge very early during treatment when high drug doses are used.

Prior studies have estimated that the probability of the existence of a fitness cost-free bacterial pathogen prior to treatment ranges from 5 x 10^-5^ to 3 x 10^-4^ per infection [68]. We sought to determine the range of drug concentrations over which the likelihood of secondary mutation emergence during treatment is at least as high as the likelihood for preexisting secondary resistance. We therefore determined the drug concentration at which the probability of secondary mutation emergence before population extinction equals 10^-4^ (that is, each treatment course has a 1 in 10,000 chance of giving rise to a resistant strain bearing secondary mutations). We used this value as an upper boundary for the “secondary mutant selection window,” the range of drug concentrations over which the probability of the emergence of a drug-resistant bacterial strain with secondary mutations is substantial (**Supplementary Figure S11**). The secondary mutant selection window extends the range of drug concentrations defined by the resistance selection window over which drug-resistant strains may be selected (**Figure 5E**).

As with the resistance selection window, we found that the size of the secondary mutant selection window varies dramatically depending on a drug’s mechanism of action. Drugs with linear effects on cellular growth and death have larger secondary mutant selection windows than do drugs with stepwise effects on cellular growth and death. This is because for drugs with stepwise effects, it is possible to shift the entire distribution of target occupancy to a range where bacterial replication is virtually eliminated (or where bacterial death far outweighs replication) across the entire population. With linear action, replication can still occur even at high target occupancy, enabling the emergence of mutants. Drugs that fully suppress cellular replication above MIC (i.e. Bacteriostatic/Stepwise) have small secondary mutant selection windows, as the probability that additional mutations emerge over the course of treatment is equal to the probability that a resistant strain with secondary mutations emerges during the transient phase of drug-target binding immediately after treatment onset, which lasts on the order of a few hours (**Figure 1C**).

## 3. Discussion

The increasing prevalence of first line- and multi-drug resistant bacteria [1, 2] signals the need for new antibiotics and robust drug dosing strategies that minimize the emergence and spread of resistance [4]. Despite this need, little is known about the role that a drug’s mechanism of action plays on the evolution of antibiotic resistance. We studied the relationship between drug mechanism and drug resistance using a mathematical model that connects bacterial population dynamics with molecular-scale descriptions of drug-target binding kinetics (**Figure 1A**). Our biochemical model allows us to describe bacterial replication and death as functions of drug-target occupancy, enables us to estimate molecular kinetic parameters from population-scale data, and delivers performance on par with that of classical pharmacodynamic models (**Figure 2B**).

We calibrate the model to an experimental dataset of ciprofloxacin time-kill curves (**Figure 2A, Table 1**), and we show that drug-resistant strains can incur strikingly high fitness costs associated with mutations that reduce drug-target binding kinetics (**Figure 3**). We find that the relationship between drug-target inactivation and antibiotic effect (i.e. bacterial killing, growth stalling, or both) exerts a strong influence on the strength of selection for resistant strains during treatment, regardless of whether the drug is bactericidal or bacteriostatic (**Figure 4D**). We also show that the molecular kinetics of drug-target binding within cells results in heterogeneous replication rates among members of an otherwise homogenous population (**Figure 5B**). This enables some drug-resistant strains to develop secondary mutations that can further reduce drug susceptibility, increase resilience in drug-free environments, and ultimately lead to treatment failure.

The clinical consequence of the frequently-observed trade-off between bacterial fitness and drug resistance [10] is the existence of a resistance selection window—a range of drug concentrations that selects for the propagation of drug-resistant strains over their drug-susceptible counterparts [5, 15]. It is important to note that numerous factors not captured by the resistance selection window can contribute to resistance selection in clinical settings, most notably ecological interactions between drug-susceptible strains, drug-resistant strains, and host physiology [69]. Our approach nonetheless enables us to isolate the roles that a drug’s mechanism of action play in driving the emergence of resistance.

We show that the resistance selection window is narrower for drugs that exert their effects on growth or death in a stepwise (i.e. sudden) manner, resulting in a steeper pharmacodynamic curve (**Figure 4C-4D, Supplementary Figure S8**). This result is consistent with other studies on the pharmacodynamics of antimicrobial agents, which have found that the size of the resistance selection window is associated with the steepness of the pharmacodynamic curve [17, 19, 64]. The characteristics of antimicrobial agents that enable steeper pharmacodynamic curves nonetheless remain poorly described. Models that capture the effects of antibiotic drugs on multiple scales, such as that described in this study and elsewhere [34, 35], could serve as helpful tools for studying the interplay between a drug’s molecular mechanism and its effect on bacterial population dynamics, enabling the design of new antimicrobial agents with narrow resistance selection windows.

Mutations that alleviate the fitness costs associated with drug resistance and/or that further raise a strain’s MIC play a major role in driving the spread of antimicrobial resistance across bacterial populations and clinical settings [24]. Our study sheds quantitative light on the mechanistic factors that govern the emergence of these secondary mutations during treatment. We propose the use of the secondary mutant selection window (**Supplementary Figure S11**) as a tool for illustrating the likelihood of further mutation acquisition at nonzero drug concentrations. As with the size of the resistance selection window, the size of the secondary mutant selection window varies greatly depending on the mechanism of action of the antibiotic (**Figure 5E**). We stress that the secondary mutant selection window does not necessarily indicate a region on the pharmacodynamic profile of a drug over which the selection of a resistant strain with secondary mutations is favored. The strength of selection depends on the physiological effect of the secondary mutation itself—that is, whether the mutation accelerates growth rate, slows drug-target binding, or exerts a multitude of other possible effects. Indeed, secondary mutations that act strictly by restoring growth rates to wild-type levels lead only to modest (usually sublinear) increases in MIC (**Figure 4B**), implying that strains with cost-free resistance phenotypes would still have MICs well below the upper boundary for the secondary mutant selection windows shown in **Figure 5E**. Rather, the secondary mutant selection window defines the drug concentration range within which the accumulation of secondary mutations is substantial and therefore clinically significant.

Suppressing secondary mutation is crucial for reducing the survival of drug-resistant mutants in antibiotic-free environments, where drug-resistant strains enter into direct competition with other microbial organisms for limited resources [10, 23]. We demonstrate that dosing drugs at or slightly above the MIC of a resistant strain may not be sufficient for preventing the spread of resistance, and that—for drugs with linear effects on bacterial growth and death as a function of drug-target occupancy—there exist appreciable risks of selecting for secondary mutations even at doses substantially above the MIC of the resistant strain. Reassessing the range of drug concentrations that selects for resistant mutants as a composite of the resistance selection window and the secondary mutant selection window (**Figure 5E, Supplementary Figure S11**) could facilitate the design of drug dosing strategies that holistically mitigate the emergence and spread of resistance.

Our study shows that both bactericidal and bacteriostatic drugs are capable of exhibiting narrow resistance selection windows and low probabilities of secondary mutation emergence in bacterial populations subjected to antibiotic treatment. This finding challenges the long-accepted notion that bactericidal agents are superior to bacteriostatic agents in suppressing the emergence of resistance during treatment [31], and signals the need to look beyond a drug’s ability to kill or stall bacterial replication to assess the risks of resistance emergence. The relationship between drug-target inactivation and overall antibiotic effect has a much stronger influence on the strength of resistance selection than does the drug’s bacteriostatic or bactericidal activity (**Figure 4D**). The processes that may dictate such a relationship for any given antibiotic nonetheless remain enigmatic. This underscores the need for deeper experimental and theoretical research on the molecular processes that govern the pharmacodynamics of antibiotic drugs.

We note that the model reported here makes a number of simplifying assumptions that limit its scope and generalizability. One key assumption made is that growth and death rates are monotonically decreasing and increasing functions, respectively, of drug-target occupancy. Non-monotonic dose-response curves have been described for numerous drugs since the early years of the antibiotic era [70], and these imply the existence of non-monotonic drug-target occupancy schemes or of drug-induced cellular responses (such as reduced outer membrane permeability) that lower drug-target occupancy at high antibiotic concentrations. Our model also has limitations on the scope of resistance mechanisms that it can recapitulate—a consequence of the trade-off between mathematical tractability and generalizability. While some classes of antibiotics (particularly fluoroquinolones and rifamycins) frequently elicit resistance through altered drug-target affinity, other classes elicit resistance through additional mechanisms (including drug efflux, enzymatic degradation, and off-target binding) not captured in the linear model presented here. Other models have been devised that link these additional mechanisms of resistance (such as efflux pump activity, membrane permeability, and cellular metabolic states) with critical pharmacologic parameters (i.e. MIC) [30, 71], but do not accommodate explicit descriptions of an antibiotic’s mechanism of action. Other models have provided valuable insights into the genotypic determinants of antimicrobial resistance fitness landscapes [72]. Adapting existing models to study the relationship between antibiotic mechanism, fitness cost, and other mechanisms of resistance constitutes a promising direction for future research.

### 3.1. Conclusions

The proper use of antibiotics in clinical and non-clinical settings constitutes a core action for addressing the worldwide threat of antibiotic resistance [4]. The quantitative approach we present in this study may prove useful for identifying strategies that manage the emergence of resistance to existing and future antimicrobial agents. We argue that dosing regimens should account for a drug’s resistance and secondary mutant selection windows if they are to minimize the selection of resistance phenotypes during treatment. Our findings suggest that even drugs with seemingly straightforward pharmacodynamic classifications (i.e. bacteriostatic versus bactericidal action) can set bacterial populations on complex and sometimes counterintuitive evolutionary trajectories with respect to resistance selection. In the clinic, there exists little evidence that bactericidal antibiotics lead to more favorable outcomes than do bacteriostatic antibiotics, especially for combatting uncomplicated infections [57, 73]. Yet it is precisely in the treatment of uncomplicated, drug-susceptible infections that the greatest gains are to be made in mitigating the emergence of resistance. Mechanistic models such as that presented in this study can help to uncover clinically useful drug characteristics that classical models may overlook. We envision a coupling of our quantitative approach with high-throughput experimental platforms [74, 75] to aid in the development of new drugs with optimal pharmacodynamic profiles and to accelerate the discovery of drug- and pathogen-specific dosing regimens that minimize resistance emergence.

## 4. Methods

### 4.1. Bacterial time-kill curve experiment

We conducted time-kill curve experiments using *Escherichia coli* strain BW25113 (Coli Genetic Stock Center #7636) [76]. We diluted liquid overnight cultures of BW25113 1:1000 into pre-warmed lysogeny broth (LB) and grew cells to an optical density at 600nm (OD_600_) of 0.50. We then prepared a 1:3 dilution series of ciprofloxacin (highest concentration: 2.19 *µ*g/ml) and added the antibiotics to bacterial cultures. We quantified bacterial population sizes at regular (20-30 min) time intervals by plating a 1:10 dilution series of liquid culture onto LB agar plates and counting colony forming units. We performed colony counting blind to plate condition, and we did not exclude any plates from the analysis. To keep shot noise below 15% during colony counting, we quantified plates with 50 or greater colony forming units.

To further assess the biological reproducibility of our experiment, we repeated cytotoxicity assays on different days, once with a fixed timepoint measurement at 90 minutes post-drug exposure, and another with a timecourse (i.e. that presented in **Figure 2A** and **Supporting Data File S1**). When compared at matching timepoints of drug exposure (90 minutes), dose-response data from these biological replicates collected on different days were highly reproducible, with Pearson correlation of 0.987, *p* < 10^-5^. Each time the experiment was performed, counts of colony forming units before drug treatment were conducted in technical triplicate.

The time-kill curve obtained at the highest ciprofloxacin concentration (2.19 µg/ml, ∼250x MIC) was used to determine the maximum death rate (*D_N_*) of bacterial cells, and a growth curve obtained using the same protocol with the omission of ciprofloxacin was used to determine the maximum growth rate (*G_0_*) of cells in an antibiotic free environment (**Supplementary Figure S6**).

### 4.2. Model formulation and analysis

Our biochemical model constitutes a system of linear ordinary differential equations that describe how successive numbers of inactivated drug-target complexes affect bacterial replication and death. We consider a population of initial size *B_0_* of phenotypically homogenous bacteria exposed to a constant concentration *C_0_* of drug. When no drug is present, bacterial cells replicate at a rate *G_0_* and die at a rate *D_0_*. All cells have an identical number *N* of proteins that drug molecules target for inactivation. We assume first-order kinetics for drug-target binding: drug molecules bind to cellular protein targets within cells, thereby inactivating the protein, at a rate *k_F_*. Inactivated drug-protein targets dissociate at a rate *k_R_*. The first-order affinity of the drug to its protein target (*K_D_*) is therefore the ratio of the molecular dissociation rate to the molecular on-rate (*K_D_* = *k_R_*/*k_F_*).

We stratify the entire bacterial population into *N*+1 subpopulations according to the number *i* of inactivated drug-target complexes within each cell (i.e. the drug-target occupancy), and we assume that growth and death rates of each bacterial cell depend on the drug-target occupancy. That is, bacterial subpopulations with a higher drug-target occupancy have slower growth rates and/or higher death rates than do bacterial subpopulations with a lower drug-target occupancy. Growth rate is therefore a monotonically decreasing discrete function of *i* (*G*[*i*]), and death rate is a monotonically increasing discrete function (*D*[*i*]). We use generalized logistic equations (**Supplementary Figure S1**) to describe overall growth and death rates as a function of drug-target occupancy, allowing these functions to take the form of a line, a sigmoidal curve, an exponential curve, or a step function. We assume that when a drug inactivates all *N* protein targets in a cell, growth rate falls to zero (for bacteriostatic drugs), death rate attains a maximal value *D_N_* (for bactericidal drugs), or growth and death rates are both affected (for drugs with mixed bactericidal and bacteriostatic action). In all of these cases, the maximal rate of killing or growth attenuation can occur before all *N* target proteins are inactivated if, for instance, *G*[*i*] and/or *D*[*i*] are step functions with inflection points between 0 and *N*. During replication, a bacterial cell partitions its inactivated drug-target complexes to two daughter cells according to a binomial distribution.

The change over time in the number of bacterial cells with exactly *i* inactivated drug-target complexes (*B_i_*) thus depends on the growth rate *G_i_*, the death rate *D_i_*, and the binding kinetics of the drug to its protein target:

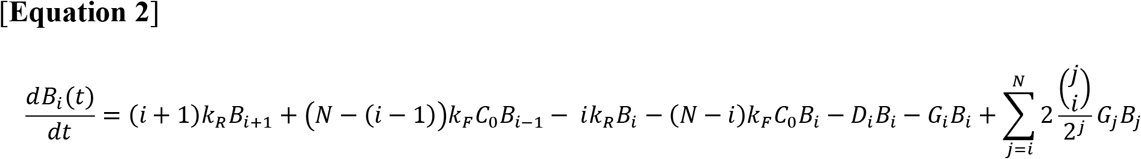

The first four terms on the right side of **Equation 2** describe changes in *B_i_* due to drug-target binding and unbinding. The fifth term describes bacterial death, the sixth term describes bacterial growth, and the seventh term describes the partitioning of drug-target complexes upon replication according to a binomial distribution. **Equation 2** is a linear form of a model we have described previously that treats drug-target complex number as a continuous variable rather than as a natural number [34]. Linearization allows us to define *B*(*t*) as a vector whose elements comprise the set of all bacterial subpopulations (*B_0_*, *B_1_*, …, *B_i_*, …, *B_N-1_*, *B_N_*) at a given time *t*. We can then describe the temporal dynamics of the entire bacterial population as a system of linear differential equations:

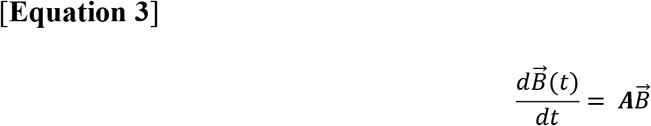

In the equation above, ***A*** denotes the matrix of coefficients describing the system of equations for the vector *B*(*t*). The values for the coefficients in ***A*** depend on the concentration *C_0_* of drug, on the drug’s binding kinetics, and on the growth and death rate functions *G*[*i*] and *D*[*i*].

**Equation 3** represents an initial value problem. This system of linear differential equations with a constant coefficient matrix has a unique solution given by

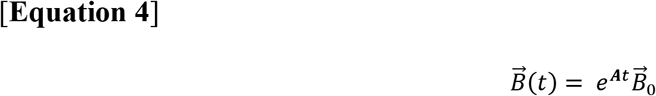

where the vector *B̄*_0_ denotes the initial composition of bacterial subpopulations at *t* = 0. The solution can also be written as a linear superposition of a product of complex exponentials (with arguments determined by eigenvalues) and polynomials (whose degree is determined by the geometric multiplicity of these eigenvalues and whose coefficients are uniquely determined by the initial conditions). In practice, *B*(*t*) describes a family of exponential growth and decay curves that represent the replication and death of all *N*+1 bacterial subpopulations over time (**Figure 1B**). We solve for *B*(*t*) numerically by calculating the matrix exponential of ***A*** using a scaling and squaring algorithm implemented in MATLAB (MathWorks, Newton, MA) [77].

### 4.3. Calculation of minimum inhibitory concentration

We define the MIC as the concentration *C_0_* of an antibiotic such that any concentration of drug at or above *C_0_* is guaranteed to cause the eventual extinction of the bacterial population. This occurs precisely when one eigenvalue of matrix ***A*** (from **Equation 3**) is zero and all other eigenvalues have a negative real component. We thus express the MIC as

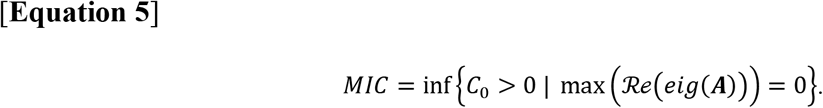

With this formulation, finding the MIC amounts to finding the value of *C_0_* such that the greatest real component of the eigenvalues of ***A*** is zero. Deriving the expression for the MIC in the simplest case of the model, when *N* = 1, serves to illustrate this approach. For the purposes of this derivation, we consider a drug that elicits both a bactericidal and a bacteriostatic effect, so *G*[*i* = 1] = 0 and *D*[*i* = 1] = *D_N_*. However, the approach for finding the MIC is identical for any mechanism of drug action. The matrix ***A*** describing all bacterial subpopulations (*B_i=0_* and *B_i=1_*) in this simple case is

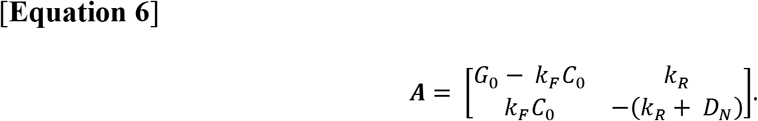

We wish to find the concentration *C_MIC_* of antibiotic that yields negative real components of all but one eigenvalues *λ* of matrix ***A***. For the 2-by-2 matrix given by **Equation 6**, the characteristic polynomial is given by *λ*^2^ -tr(***A***)*λ* + det(***A***), and the Routh-Hurwitz stability criterion needed to satisfy the negative value constraints on *λ* is tr(***A***) ≤ 0 and det(***A***) ≥ 0. For the matrix described in **Equation 6**, these expressions correspond to

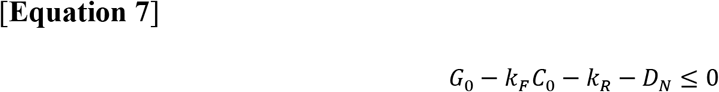

and

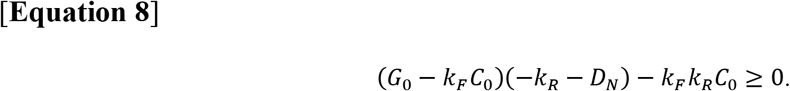

Solving for the concentration *C_o_* in both of these cases yields

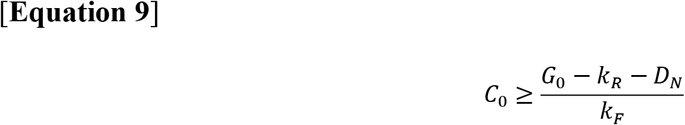

in the case of **Equation 7** and

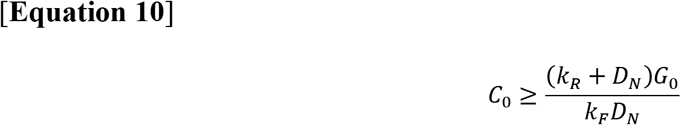

in the case of **Equation 8**. We expect the value of *k_R_* to be greater than that of *G_0_* (that is, we expect the rate of drug-target unbinding to be greater than the rate of bacterial replication). We also expect the value of the death rate at saturating drug concentrations (*D_N_*) to be nonzero and positive. Therefore, **Equation 9** is guaranteed to be satisfied if **Equation 10** is also satisfied. We thus find the expression for the MIC to be

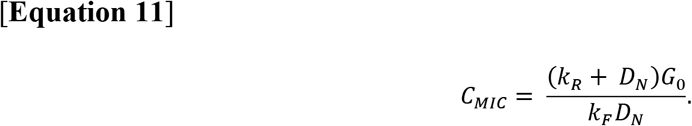

From this expression, we can infer the following proportionalities for the value of the MIC relative to the values of other model parameters:

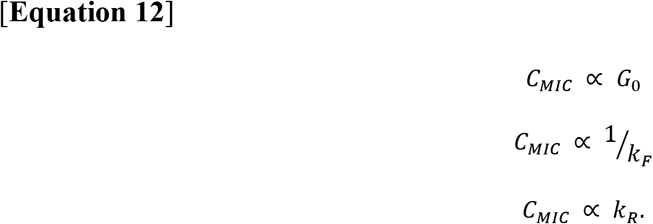

Polynomial expressions for the MIC, as shown in **Equation 11**, become exceedingly complex beyond *N* = 3. However, we conjecture (although we have not been able to prove) that the structure of the linear system shown in **Equation 3** guarantees the existence of the MIC for any *N*. For larger values of *N*, we leverage numerical schemes to calculate the eigenvalues of matrix ***A***. We use MATLAB’s *eig()* function, which calculates eigenvalues using the QZ algorithm [78].

### 4.4. Model calibration via simulated annealing

Numerical values for the model parameters *N*, *D_0_*, *µ_R_*, and *µ_C_* were obtained from the literature (**Table 1**). The values for *G_0_* and *D_N_* were obtained by fitting experimental kill curves at drug concentrations of zero and 2.19 *µ*g/ml, respectively, to exponential functions (**Supplementary Figure S6**). We leveraged an adaptive simulated annealing algorithm coupled with local gradient descent to obtain the remaining parameters (*k_F_*, *k_R_*, *α_G_*, *α_D_*, *γ_G_*, and *γ_D_*). Detailed descriptions of the adaptive simulated annealing algorithm are available elsewhere [79, 80]; in brief, simulated annealing is a global optimization algorithm capable of escaping local minima. It is therefore well suited to applications involving the optimization of many parameters. Adaptive simulated annealing is a variant on the classical simulated annealing algorithm that probes global parameter space with greater efficiency by accounting for each parameter’s magnitude when formulating a new parameter set at every iteration of the algorithm. We used adaptive simulated annealing to minimize the difference between experimental time-kill curves and model simulations of bacterial populations challenged to the same antibiotic doses. The difference between experimental observation and simulation is expressed through the objective function, whose value *ψ* the algorithm seeks to minimize:

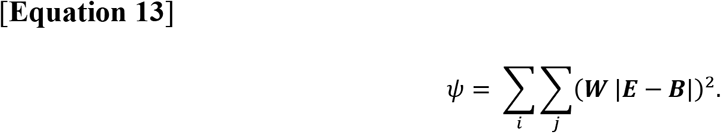

***E*** denotes an *m*-by-*n* matrix of experimentally-measured population sizes at *m* drug concentrations and *n* timepoints, ***B*** denotes simulated population sizes at the same drug concentrations and timepoints, and ***W*** denotes an *m*-by-*n* weighting matrix (for our application, simply a matrix of ones). ***B*** is a function of the parameters being optimized (that is, ***B*** = *f*(*k_F_*, *k_R_*, *α_G_*, *α_D_*, *γ_G_*, *γ_D_*)).

Coupling the adaptive simulated annealing optimization with a local gradient descent assures that our calibration procedure always converges on a local minimum. We used an exponential cooling schedule for the simulated annealing algorithm, which allows the optimization to run ergodically [79]. That is, repeating the optimization many times from random initial starting conditions in parallel yields roughly the same results as running the optimization once for a very long time. This allowed us to parallelize the optimization procedure by running the algorithm repeatedly across several cores of a computer and to characterize the distributions of parameter values obtained from these calibrations (**Supplementary Figure S3**). After performing 249 independent model calibrations, we selected the parameter set with the lowest objective function value to use in subsequent simulations. The parameter values for this set are shown in **Table 1**. Parameter sets for all model optimizations performed are available in **Supporting Data File S3**.

### 4.5. Simulating the emergence of secondary mutations

We assumed that drug-resistant bacterial strains with secondary mutations that compensate for fitness costs and/or that further increase MIC emerge from preexisting drug-resistant subpopulations present in the initial population at the start of treatment (**Supplementary Figure S9**). The size of the drug-resistant subpopulation in the absence of antibiotic (*B_0,R_*) is given by the mutation-selection balance, which expresses the equilibrium at which the rate of emergence of drug resistance alleles by spontaneous mutation equals the rate of elimination of those alleles due to competitive fitness costs [81]:

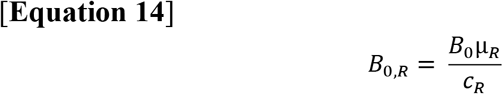

Here, *µ_R_* denotes the mutation rate for drug resistance emergence per unit time.

In order to quantify the probability of secondary mutation emergence from this drug-resistant subpopulation, we adapted a formulation that Lipsitch and Levin developed to study the evolution of drug-resistant bacterial strains during antibiotic treatment [67]. We assumed that secondary mutations emerge exclusively due to errors in DNA replication during bacterial growth. The expected number of resistant cells with secondary mutations that emerge from a bacterial population with *i* inactivated drug-target complexes (*E*(*M_RC,i_*)) is proportional to the total number of replications that the subpopulation undergoes before extinction and the rate of secondary mutation emergence:

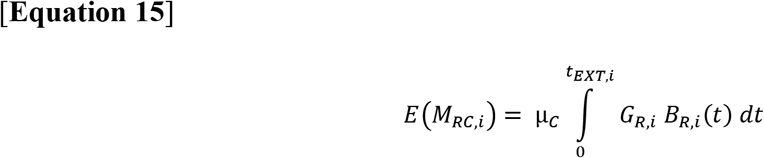

In this equation, *µ_C_* denotes the secondary mutation rate, *G_R,i_* represents the growth rate of a resistant strain with exactly *i* inactivated drug-target complexes, *B_R,i_*(*t*) describes the population dynamics of the *i*th drug-resistant bacterial subpopulation, and *t_EXT,i_* describes the amount of time elapsed from treatment onset until the bacterial subpopulation is eliminated (*B_R,i_* = 1 when *t* = *t_EXT_*). The total number E(*M_RC_*) of resistant mutants with secondary mutations that we expect to observe over the course of treatment is thus the sum of **Equation 15** over all values of *i*, and the probability *P_RC_* that a compensated resistant mutant will emerge over the course of treatment follows from the Poisson assumption that secondary mutations arise stochastically and independently of other mutations:

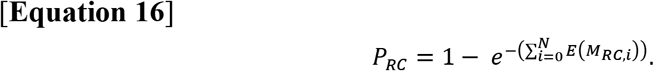

The summation term in **Equation 16** describes the total number of resistant strains with secondary mutations expected to emerge before extinction. This equation thus quantifies the Poisson probability that at least one resistant strain with a secondary mutation will emerge over the course of treatment.

### 4.6. Code and data

We wrote all code in MATLAB. All of the code used to implement and solve our model, to analyze experimental data, and to generate simulation data shown in all figures is available as a software package in **Supplementary File S1**. Experimental data represented in **Figures 2A** & **2B** and in **Supplementary Figure S4** are available within **Figure 2 – Source Data Files 1, 2** & **4**, respectively, and the parameter values for all iterations of model optimization are available in **Supporting Data File S3**.

## Acknowledgements

We extend sincere thanks to Sören Abel, Benjamin Akhuetie-Oni, and Laura Quinto for helpful feedback on the manuscript.

## Author contributions

Conceptualization: P.AzW.

Data curation: C.H.

Formal analysis: L.C., C.H., F.C., P.AzW.

Funding acquisition: P.AzW., T.C.

Investigation: C.H., F.C., A.P.

Methodology: L.C., P.AzW., C.H., A.P.

Project administration: P.AzW., T.C.

Resources: A.P., T.C., P.AzW.

Software: C.H.

Validation: C.H., L.C.

Visualization: C.H.

Writing – Original draft preparation: C.H.

Writing – Review & editing: C.H., P.AzW., T.C., F.C., L.C., A.P.

---

**Supplementary Figure S1.**
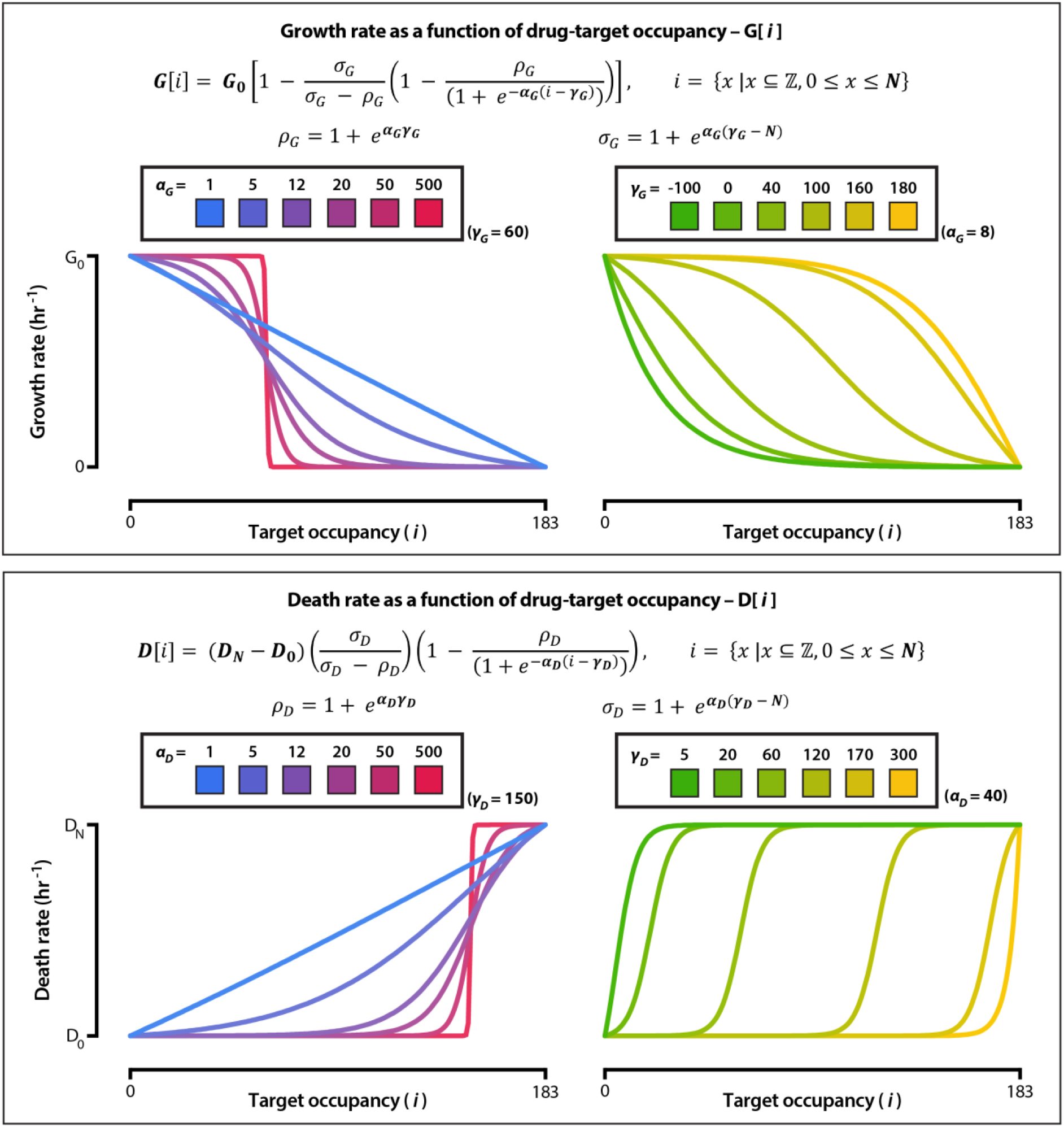
Bacterial growth and death rates as a function of drug-target occupancy. We define the functions G[*i*] and D[*i*] as constrained logistic curves such that G[*i* = 0] = *G_o_*, G[*i* = *N*] = 0, D[*i* = 0] = *D_0_*, and D[*i* = *N*] = *D_N_*. The parameters *α_G_* and *α_D_* define the steepness of the logistic curves for the growth and death rate function, respectively. *α_G_* and *α_D_* are unitless and range from 1 to 500; 1 yields a quasi-linear function, while 500 yields a quasi-step function. The parameters *γ_G_* and *γ_D_* define the inflection point of the logistic curves for the growth and death rate function, respectively. *γ_G_* ranges from –*N* to *N* and *γ_D_* ranges from 0 to 2*N*; the curve is quasi-sigmoidal if *γ_G_* and *γ_D_* are in between 0 and N and is quasi-exponential if *γ_G_* and *γ_D_* are outside of these bounds.

**Supplementary Figure S2.**
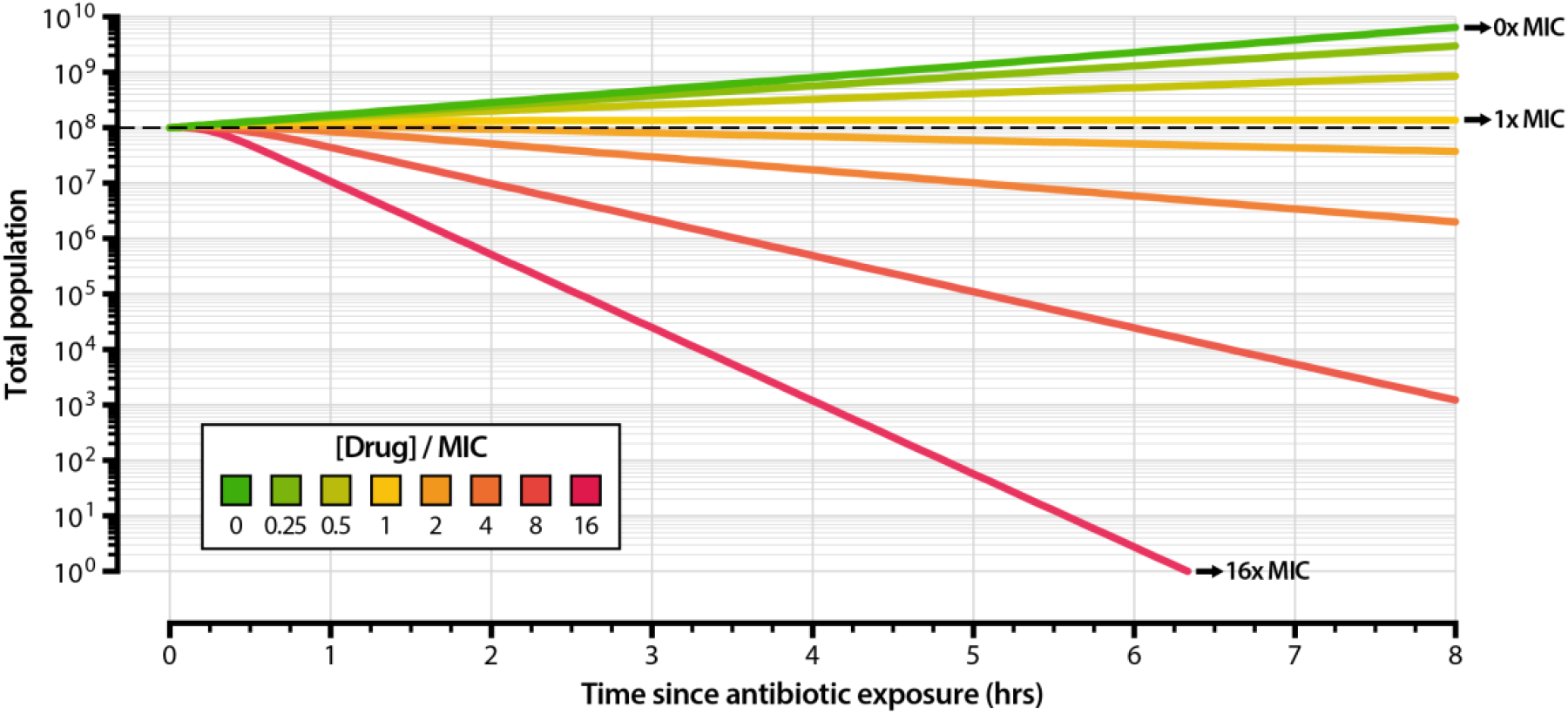
Simulated time-kill curves of *Escherichia coli* exposed to a range of drug concentrations. We used the parameter set outlined in **Table 1** to model the growth and death of bacterial populations subjected to drug concentrations up to 16x minimum inhibitory concentration (MIC). Drug concentrations are expressed as factors of the MIC. The net growth rate of the entire bacterial population over the time course of the simulation decreases with increasing drug concentration.

**Supplementary Figure S3.**
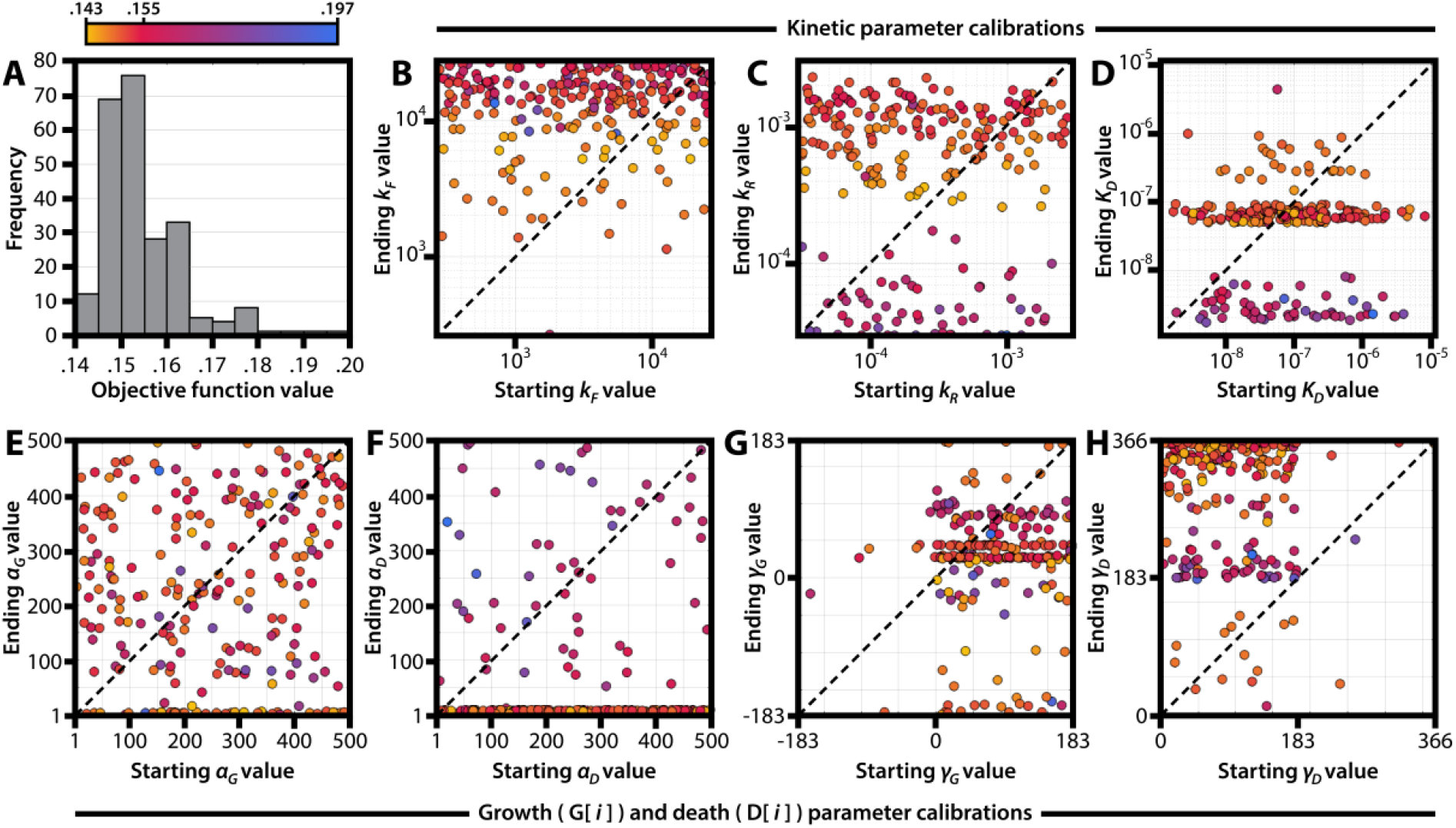
Results from 249 independent model calibrations to experimental data. We used adaptive simulated annealing coupled with gradient descent (see **Methods**, *Model calibration via simulated annealing*) to fit the model to experimental kill curve data of *E. coli* exposed to ciprofloxacin (**Supporting Data File S1**). Shown in this figure are the results for 249 independent model fits (**Supporting Data File S3**), each beginning with randomly-chosen values for the parameters describing drug-target binding rate *k_F_*, drug-target unbinding rate *k_R_*, steepness of the growth rate function *α_G_*, steepness of the death rate function *α_D_*, inflection point of the growth rate function *γ_G_*, and inflection point of the death rate function *γ_D_*. (**A**) Frequency distribution of objective function values obtained from independent model calibrations. The objective function value describes the goodness of the fit between experimental data and simulation; smaller values indicate higher goodness of fit. (**B-H**) Optimization plots showing randomly chosen initial parameter values (x-axis) and calibrated parameter values (y-axis) for all independent model calibrations. The optimized parameters are *k_F_* (**B**), *k_R_* (**C**), *K_D_* (the ratio of *k_R_* to *k_F_*) (**D**), *α_G_* (**E**), *α_D_* (**F**), *γ_G_* (**G**), and *γ_D_* (**H**). The final objective function value of each model fit is colored according to the color bar above panel (**A**).

**Supplementary Figure S4.**
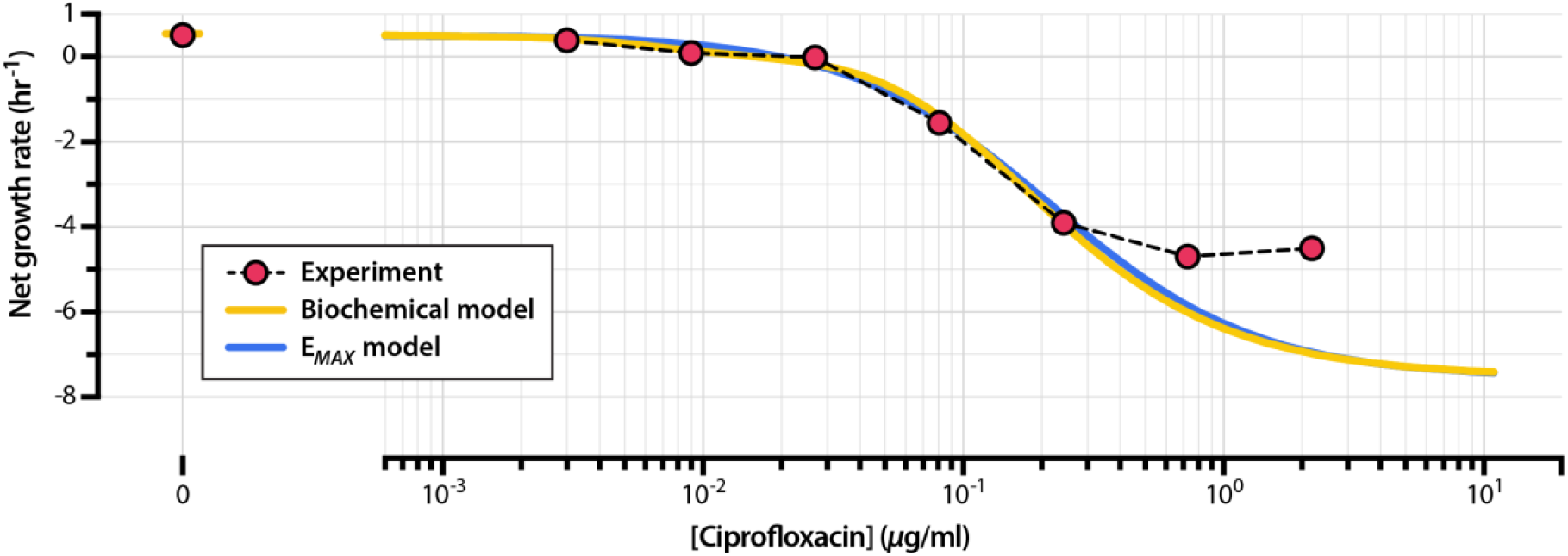
Pharmacodynamic curves generated from experimental data and from the calibrated model. The experimental pharmacodynamic curve was generated by calculating the net growth rates of *E. coli* exposed to a set of ciprofloxacin drug concentrations (**Supporting Data File S1**). The time-kill curves of this same experimental dataset are shown in Figure 2A; see **Supporting Data File S4** for experimental data on net growth rate as a function of drug concentration. The model-calibrated pharmacodynamic curve was generated by simulating bacterial time-kill curves over the same range of drug concentrations used in the experiment and calculating associated net growth rates.

**Supplementary Figure S5.**
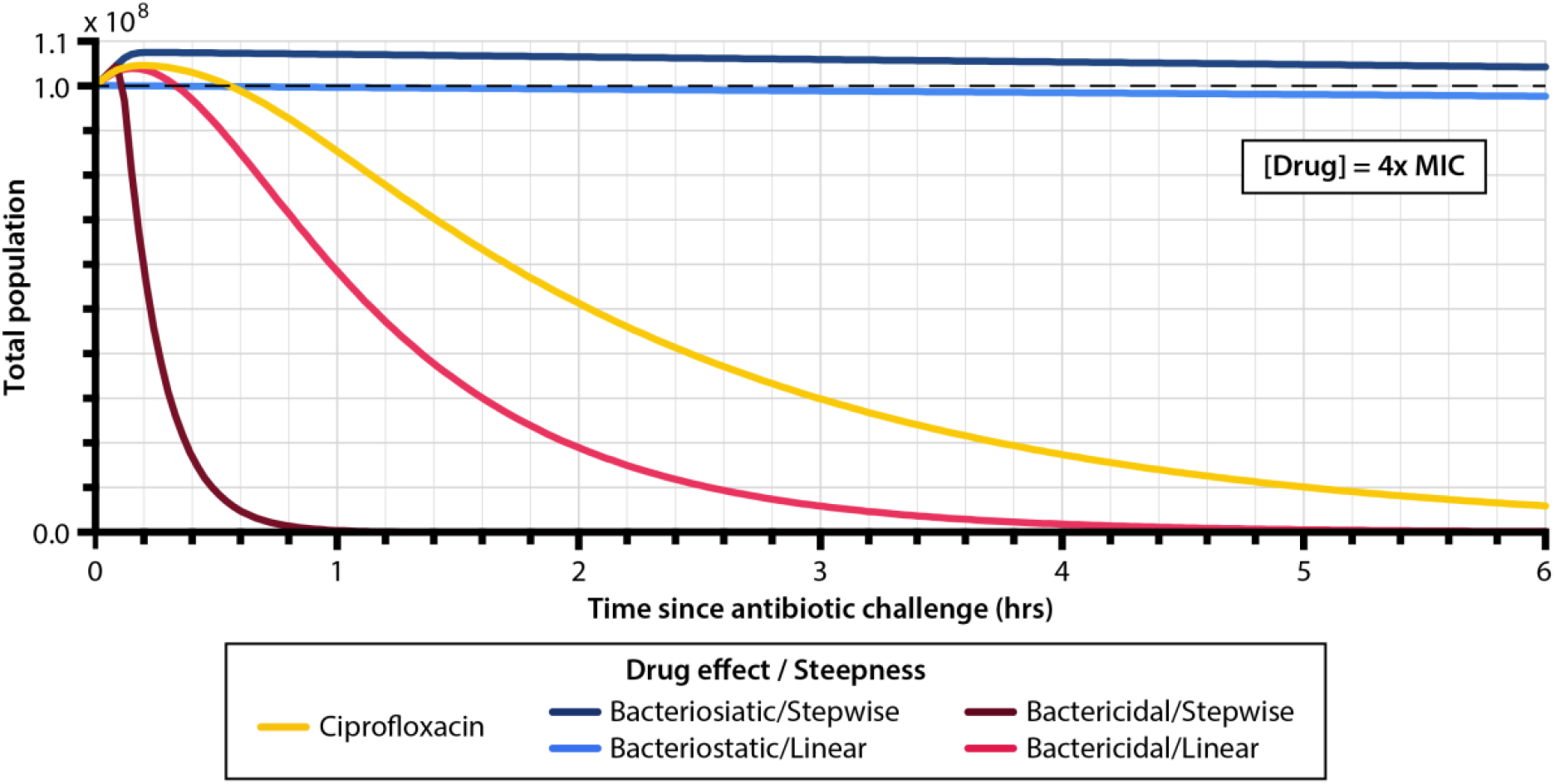
Simulated population curves for ciprofloxacin and for four extreme modes of antibiotic drug mechanism. We simulated a bacterial population of 10^8^ cells exposed to antibiotic drug at 4x MIC. The ciprofloxacin curve corresponds to the drug mechanism obtained from the model calibration to experimental data and detailed in Figure 2C, and the remaining curves correspond to the extreme schemes of drug mechanism shown in Figure 2D.

**Supplementary Figure S6.**
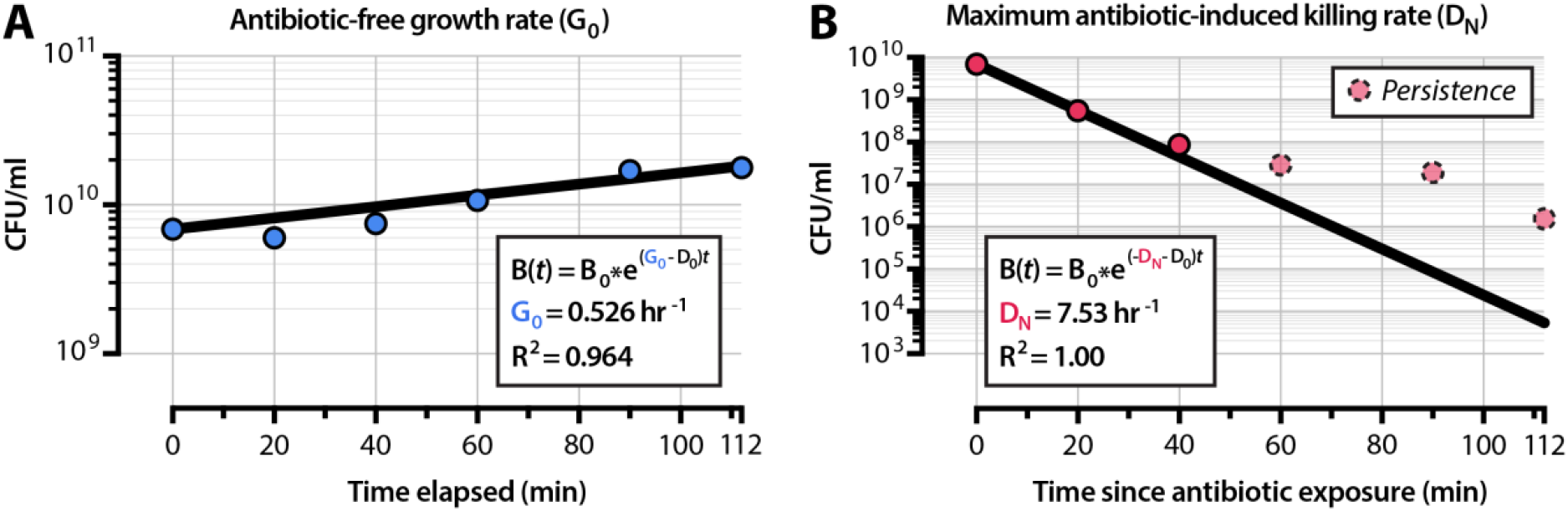
Obtaining *G_0_* and *D_N_* from experimental data. (**A**) To obtain the value of *G_0_* (growth rate in the absence of antibiotic) used in simulations, we fit an exponential growth curve to experimental data for *E. coli* cells grown in the absence of antibiotic. (**B**) To determine the value of *D_N_* (maximum death rate in saturating conditions of antibiotic), we fit an exponential decay curve to experimental data for *E. coli* cells exposed to 2.19 µg/ml of ciprofloxacin (∼200 x MIC). The population size deviates from exponential decay at later timepoints (dashed and shaded) likely because of the emergence of persistent subpopulations of bacteria [45]. The R^2^ values shown are the linear correlation coefficients for the model fit, and are not the correlation coefficients for the log-transform of the data.

**Supplementary Figure S7.**
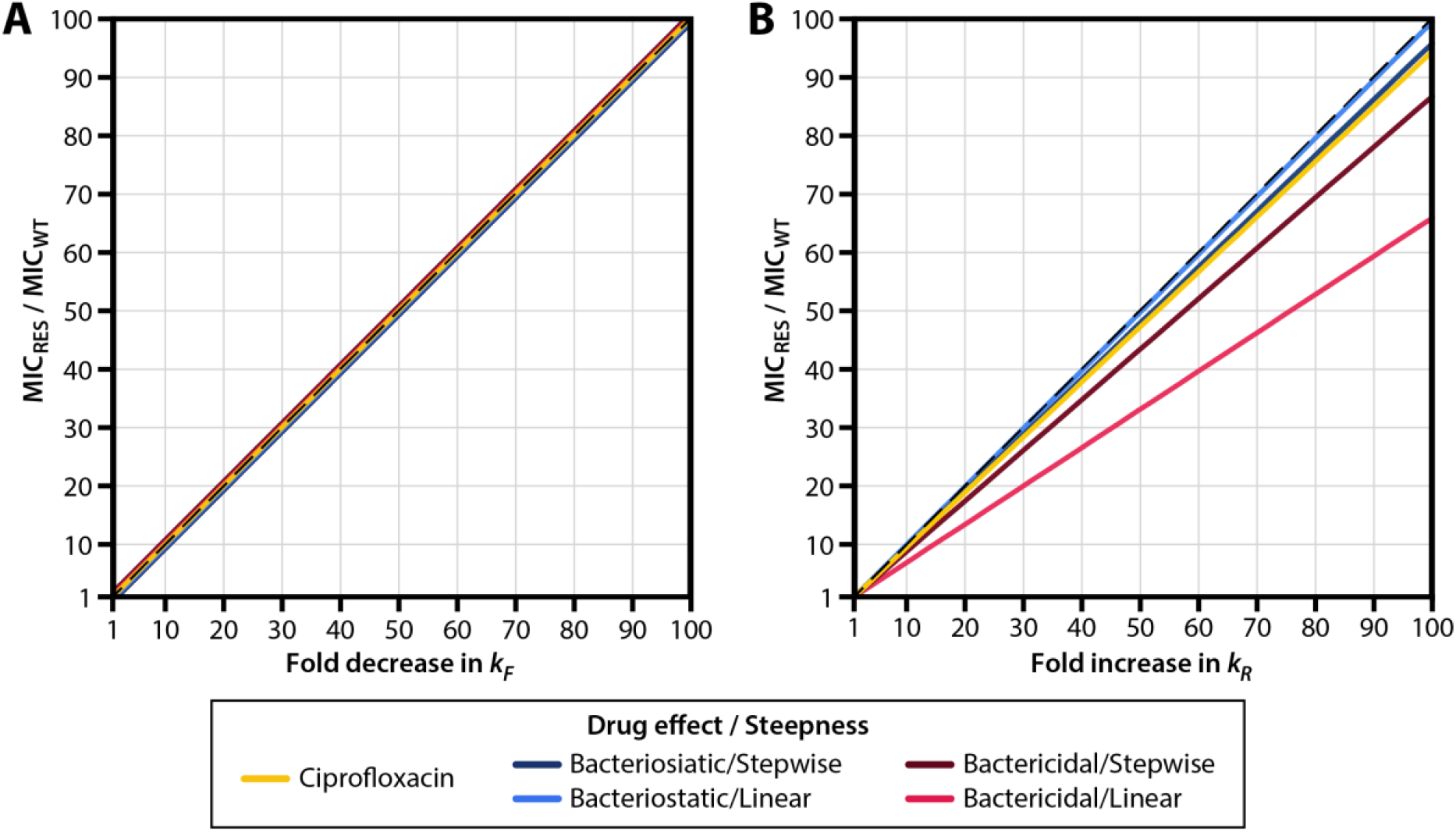
MIC as a function of drug-target binding and unbinding kinetics. The MIC of a mutant (normalized to the MIC of the wild-type) is plotted against the fold-change in (**A**) drug-target binding (*k_F_*) or (**B**) drug-target complex disassociation (*k_R_*). For this simulation, mutants have no fitness costs associated with changes in *k_F_* and *k_R_* (*c_R_* = 0). For drug-target binding (*k_F_*), fold increase in MIC is directly proportional to fold decrease in *k_F_* for all drug mechanisms. In both panels, the dashed line indicates the line of direct proportionality. MIC_WT_: MIC of the wild-type strain; MIC_RES_: MIC of the resistant strain.

**Supplementary Figure S8.**
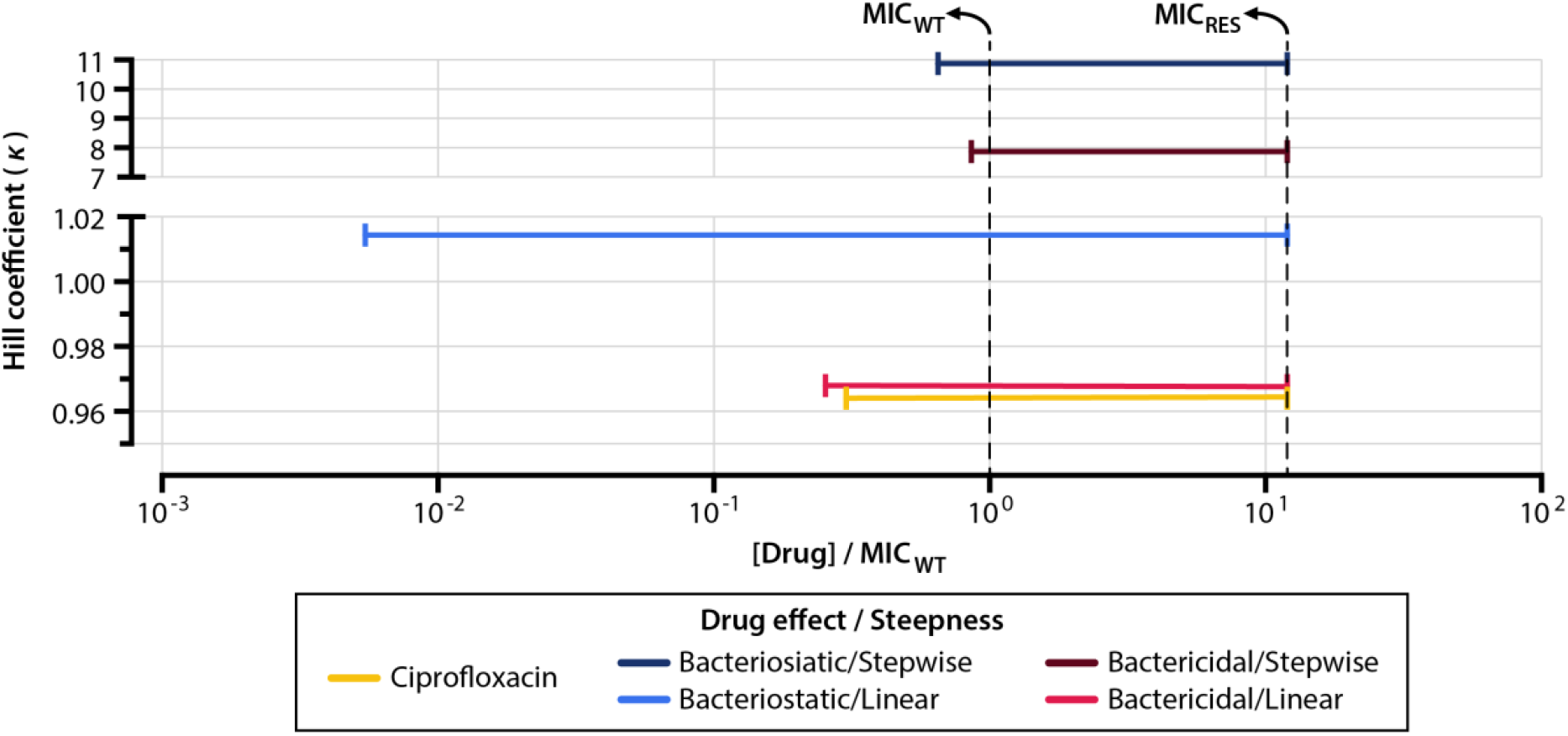
Drugs with steeper pharmacodynamic curves have narrower resistance selection windows given a cellular effect (bacteriostatic/bactericidal). To quantify the steepness of pharmacodynamic curves, we fit the curves for drug-resistant strains shown in Figure 4C to the pharmacodynamic function formulated by Regoes et al. [33]. The equation describes the net growth rate *G_net_* of a bacterial population as a function of drug concentration *C_0_* and other parameters (MIC, *G_0_*, *D_N_*) derived from the model:

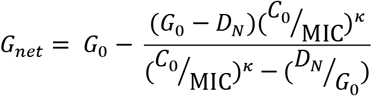

In this equation, *κ* describes the Hill coefficient, which serves as a measure of the steepness of the pharmacodynamic curve. Larger values of *κ* indicate steeper curves. For each of the drug mechanisms described in this study (**Supplementary File S2**), we generated pharmacodynamic curves for drug-resistant mutants (Figure 4C, solid lines), determined the value of *κ* that best fits the curve, and plotted *κ* against the range of drug concentrations that represents the resistance selection window (Figure 4D). MIC_WT_: MIC of the wild-type strain; MIC_RES_: MIC of the resistant strain.

**Supplementary Figure S9.**
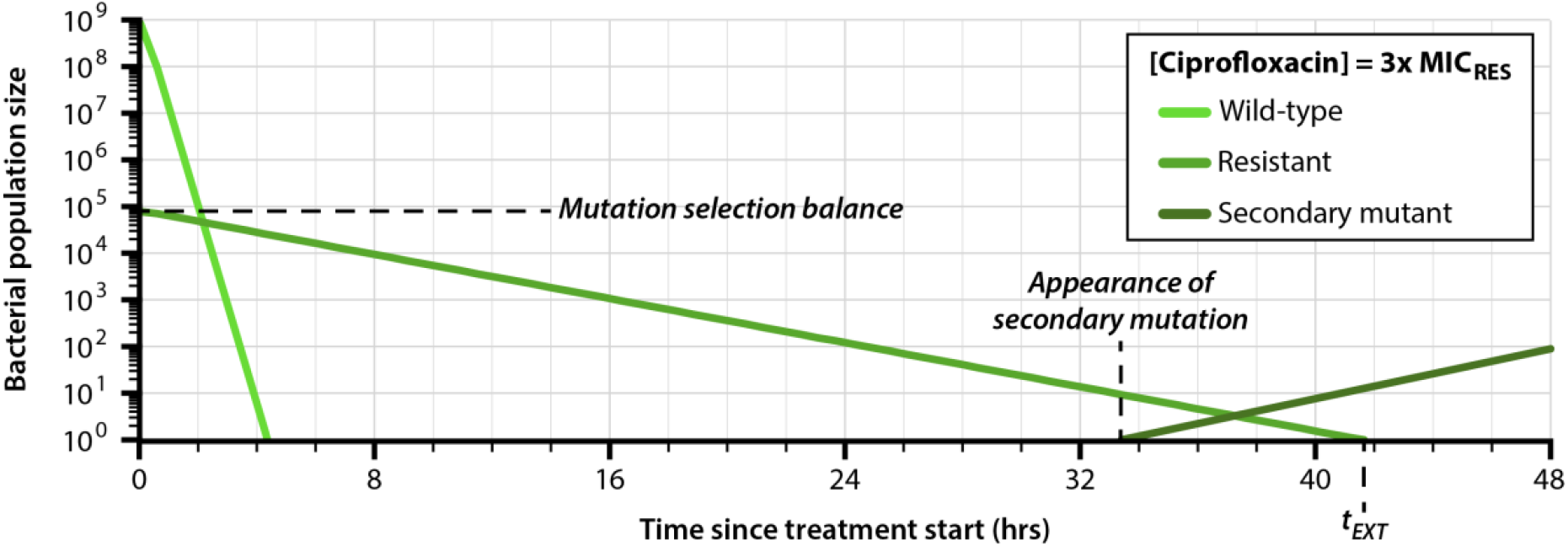
Emergence of secondary mutations within subpopulations of drug-resistant bacteria during antibiotic treatment. When simulating the emergence of secondary mutations, we assume that a drug-resistant subpopulation (middle green) of bacteria is present at the start of treatment; the size of this subpopulation is given by the mutation selection balance of the allele that confers the drug-resistance phenotype [81]. We calculate the probability that a drug-resistant strain with secondary mutations (dark green) emerges from this subpopulation before the elimination of the drug-resistant strain (at time *t_EXT_*).

**Supplementary Figure S10.**
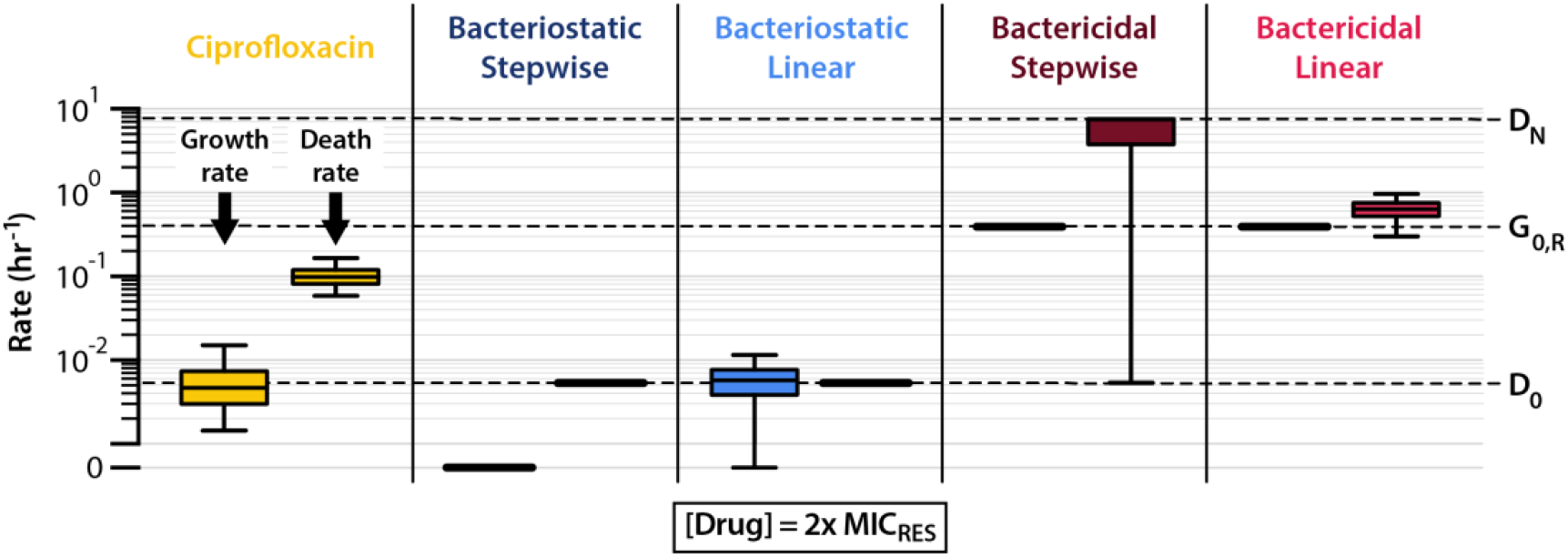
Distributions of growth and death rates for drug-resistant bacterial subpopulations undergoing steady-state exponential decline at 2x MIC_RES_. Boxes denote the central 50% of the growth and death rate distributions, and whiskers denote the central 95% of the growth and death rate distributions.

**Supplementary Figure S11.**
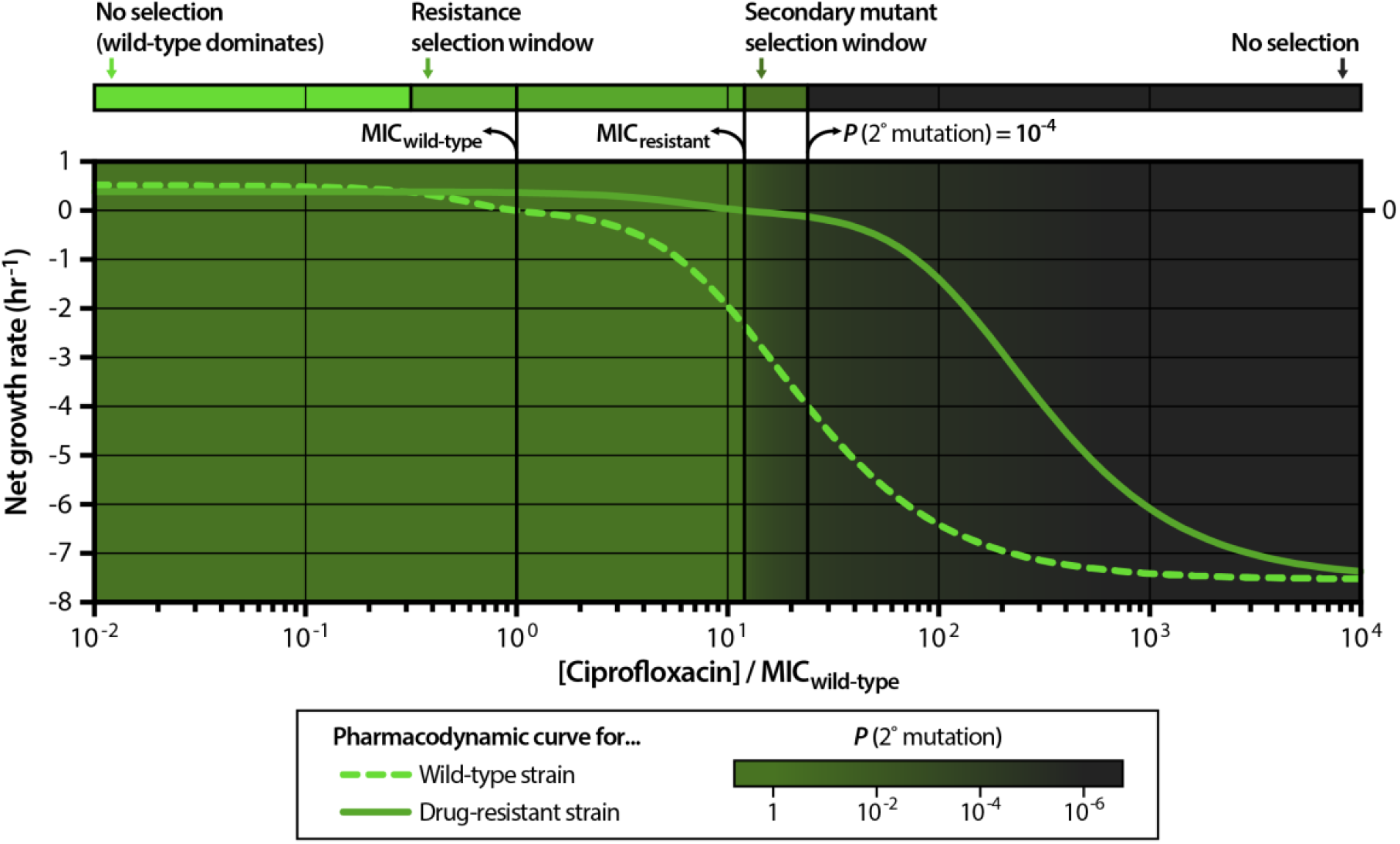
The secondary mutant selection window. The secondary mutant selection window comprises the drug concentration range over which the net growth of the drug-resistant strain is negative but the probability of secondary resistance emergence before the end of treatment exceeds a defined threshold (in our simulations, 10^-4^, or a 1 in 10,000 chance). Four regimes of selection exist: the null selection window in which the wild-type strain dominates, the resistance selection window, the secondary mutant selection window, and the complete killing window. We simplify these four regimes by disregarding the relative strengths of selection for each strain in each regime and we instead illustrate the boundaries of each region along a drug concentration axis (top bar); these simplified selection regimes are shown for all five drug mechanisms studied in Figure 5E.

## Additional files included in submission

**Supplementary File S1 – MATLAB code package containing the code written for this study.** This file contains scripts that we used to implement our model, to analyze data, and to generate simulation data for all main text and supplementary figures. Documentation detailing how to use the software is included in each script of the code package.

**Supplementary File S2 – Parameters for a set of five drugs with different mechanisms of action.** The parameters *α_G_* and *α_D_* describe the steepness of the growth and death rate functions, respectively, around the inflection point. The parameters *γ_G_* and *γ_D_* describe the inflection points of the growth and death rate functions (see Supplementary Figure S1). Bacteriostatic potency refers to the magnitude of growth rate decline at saturating concentrations of drug; a value of 1 indicates that that growth rate declines to zero in saturating concentrations of drug (*G*[*i* = *N*] = 0), and a value of 0 indicates that growth rate is unaffected by drug concentration (*G*[*i*] = *G_0_* for all *i*). Bactericidal potency refers to the magnitude of death rate increase at saturating conditions of drug; a value of 1 indicates that death rate increases to maximum in saturating concentrations of drug (*D*[*i* = *N*] = *D_N_* > *D_0_*), and a value of 0 indicates that death rate is unaffected by drug concentration (*D*[*i*] = *D_0_* for all *i*). All other parameters (including drug-target binding rate *k_F_*, drug-target unbinding rate *k_R_*, and target number *N*) are identical for all drugs in the set.

**Supporting Data File S1 – Experimental data for the ciprofloxacin time-kill curve experiment represented in Figure 2A and Supplementary Figure S6.**

**Supporting Data File S2 – Experimentally-measured minimum inhibitory concentrations (MICs) for ciprofloxacin against *Escherichia coli* represented in Figure 2B.** We collated this list of experimentally-measured MICs from the literature; study sources are given in the file.

**Supporting Data File S3 – Model calibrations obtained via simulated annealing.** Starting and ending values for all model parameters are given for each iteration of the model fitting procedure described in **Methods,** *Model calibration via simulated annealing*.

**Supporting Data File S4 – Experimental pharmacodynamic curve data represented in Supplementary Figure S4.** We generated these data by calculating the net growth rates of bacterial populations at each drug concentration in the experiment detailed in **Supporting Data File S1**.

