## Supplementary data file 1 for "Mechanisms of antibiotic action shape the fitness landscapes of resistance mutations": ABkinetic9P.m

```
% Antibiotic Simulator, version 9 FOR PARAMETERIZATION
% Finite time advancement of model by
% Pia Abel zur Wiesch
% Leonid Chindelevitch
% Colin Hemez

% Implemented by
% Colin Hemez
% 2019

% INPUTS
% kinParams: Kinetic parameters
%            Vector of form [N G0 D0 Dn kF kR aG aD bG bD cG cD]
%            N  = number of target complexes per cell [1/cell]
%            G0 = growth rate in the absence of antibiotic [1/sec]
%            D0 = death rate in the absence of antibiotic [1/sec]
%            Dn = death rate in saturating antibiotic concentration [1/sec]
%            kF = drug-target binding rate constant [1/M/sec]
%            kR = drug-target unbinding rate constant [1/sec]
%            aG = Sharpness of growth rate function G[i]
%            aD = Sharpness of death rate function D[i]
%            bG = Bacteriostatic potency of drug at high concentrations
%            bD = Bactericidal potency of drug at high concentrations
%            cG = Inflection point for growth rate function G[i]
%            cD = Inflection point for death rate function D[i]
% simParams: Simulation parameters
%            Vector of the form [nC Vtot MM]
%            nC   = initial number of cells in medium [dimensionless]
%            Vtot = total volume of the simulation [l]
%            MM   = molar mass of drug being simulated [g/mol]
% Binit:     Initial distribution of cells in each bound target category,
%            as a column vector of dimensions [N x 1]. Distributions
%            should be expressed as proportions of the total, such that
%            sum(Binit) = 1
% C0:        Concentration of drug [g/l]
% T:         Vector of timepoints to simulate [sec]

% OPTIONAL ADDITIONAL INPUTS:
%  'mechanism' specifies predefined pair of growth and death rate functions
%   Acceptable inputs are
%    'general': Ciprofloxacin parameter fir to PDE model
%    'cipro': Ciprofloxacin parameter fit to ODE model
%    'nstat': Stabilization/bacteriostatic
%    'ncide': Stabilization/bactericidal
%    'cstat': Competition/bacteriostatic
%    'ccide': Competition/bactericidal
%    'curve': Use kinParams for growth and death rate function parameters
%    If a different input is given, 'curve' is used
% 'solution' specifies the ODE solver method
%   Acceptable inputs are
%    'exact': solve ODEs using MATLAB's matrix exponentiation algorithm
%    'numerical': solve ODEs by taking small time steps from t = 0

% Additional functions needed in directory
% SYSTEMMATRICES9P()
% BFUN()
% RATEFUN()
% GROWTHLIMIT()
% DEATHLIMIT()

function [Bi,runtime,parameters] = ABkinetic9P(kinParams,...
                                               simParams,...
                                               Binit,...
                                               C0,...
                                               T,...
                                               varargin)

%%%%%%%%%%%%%%%%%%%%%%%%%%%%%%%%
%%%% SET UP THE STIMULATION %%%%
%%%%%%%%%%%%%%%%%%%%%%%%%%%%%%%%

% Check inputs
if length(kinParams) ~= 12
    error('kinParams vector is incorrect length')
end

if length(simParams) ~= 3
    error('simParams vector is incorrect length')
end

% Determine matrix construction method and solution method
defaultsit = 'curve';
defaultsol = 'exact';

if nargin == 5
    situation = defaultsit;
    sol    = defaultsol;

elseif nargin == 6
    if strcmp(varargin{1},'curve')
        situation = 'curve';
        sol    = defaultsol;
    elseif strcmp(varargin{1},'general')
        situation = 'general';
        sol    = defaultsol;
    elseif strcmp(varargin{1},'nstat')
        situation = 'nstat';
        sol    = defaultsol;
    elseif strcmp(varargin{1},'cstat')
        situation = 'cstat';
        sol    = defaultsol;
    elseif strcmp(varargin{1},'ncide')
        situation = 'ncide';
        sol    = defaultsol;
    elseif strcmp(varargin{1},'ccide')
        situation = 'ccide';
        sol    = defaultsol;
    elseif strcmp(varargin{1},'cipro')
        situation = 'cipro';
        sol    = defaultsol;
    elseif strcmp(varargin{1},'exact')
        sol    = 'exact';
        situation = defaultsit;
    elseif strcmp(varargin{1},'numerical')
        sol    = 'numerical';
        situation = defaultsit;
    else
        situation = defaultsit;
        sol    = defaultsol;
    end
    
elseif nargin == 7
    if strcmp(varargin{1},'exact') ||...
       strcmp(varargin{1},'numerical')
        sol = varargin{1};
        if strcmp(varargin{2},'general') ||...
           strcmp(varargin{2},'nstat') ||...
           strcmp(varargin{2},'cstat') ||...
           strcmp(varargin{2},'ncide') ||...
           strcmp(varargin{2},'ccide') ||...
           strcmp(varargin{2},'cipro')
            situation = varargin{2};
        else
            situation = defaultsit;
        end
    elseif strcmp(varargin{1},'general') ||...
           strcmp(varargin{1},'nstat') ||...
           strcmp(varargin{1},'cstat') ||...
           strcmp(varargin{1},'ncide') ||...
           strcmp(varargin{1},'ccide') ||...
           strcmp(varargin{1},'cipro')
        situation = varargin{1};
        if strcmp(varargin{2},'exact') ||...
           strcmp(varargin{2},'numerical')
            sol = varargin{2};
        else
            sol = defaultsol;
        end
    else
        situation = defaultsit;
        sol    = defaultsol;
    end
end

% Check and modify inputs as needed
if sum(Binit) ~= 1
    Binit = Binit ./ sum(Binit);
    disp('ABKINETIC9P says')
    warning('sum(Binit) ~= 1; scaling vector to satisfy condition')
elseif Binit == 1
    Binit = zeros(kinParams(1) + 1, 1);
    Binit(1) = 1;
end

% Unpack needed parameters
N  = kinParams(1);
G0 = kinParams(2);
D0 = kinParams(3);
Dn = kinParams(4);
kF = kinParams(5);
kR = kinParams(6);
aG = kinParams(7);
aD = kinParams(8);
bG = kinParams(9);
bD = kinParams(10);
cG = kinParams(11);
cD = kinParams(12);
states = N + 1;

nC   = simParams(1);
Vtot = simParams(2);
MM   = simParams(3);
Na = 6.022e23; % Avogadro constant

% Normalize the units
Cmol = C0 / MM;
Cn = Cmol * Vtot * Na; % Number of drug molecules in the simulation
kFn = kF / (Na * Vtot); % Rescale kF to number of molecules
B0 = Binit .* nC;

% Define parameters for growth and death rate functions
if strcmp(situation, 'curve')
    GDfunction = [aG aD bG bD cG cD];
elseif strcmp(situation, 'general')
    GDfunction = [1 500 0 1 cG cD];
elseif strcmp(situation,'cipro')
    GDfunction = [1 500 0 1 cG cD];
elseif strcmp(situation, 'nstat')
    GDfunction = [500 1 0 0 cG cD];
elseif strcmp(situation, 'ncide')
    GDfunction = [1 500 1 1 cG cD];
elseif strcmp(situation, 'cstat')
    GDfunction = [1 1 0 0 cG cD];
elseif strcmp(situation, 'ccide')
    GDfunction = [1 1 1 1 cG cD];
else
    warning('Invalid input for MECHANISM; using values from KINPARAMS')
    GDfunction = [aG aD bG bD cG cD];
end

% Repackage parameter vector using the growth and death rate functions
matrixParams = kinParams(1:6);
matrixParams = [matrixParams GDfunction];

%%%%%%%%%%%%%%%%%%%%%%%%%%
%%%% SOLVE THE SYSTEM %%%%
%%%%%%%%%%%%%%%%%%%%%%%%%%

% Time the run
tic

if strcmp(sol, 'exact')

    %%%%%%%%%%%%%%%%%%%%%%%%%%%%%%%%%%%%%
    % SOLUTION BY MATRIX EXPONENTIATION %
    %%%%%%%%%%%%%%%%%%%%%%%%%%%%%%%%%%%%%

    % Set aside memory
    Bi = zeros(states, length(T));

    % Build system matrix
    % A0 = Concentration-scaled portion of system matrix
    % A1 = Constant portion of system matrix
    [A0, A1] = systemMatrices9P(matrixParams, 0, 0);
    A = A1 + (A0 .* Cn .* kFn);

    % Generate solution in time
    % Solutions are of the form Bi(t) = exp(At)*B0
    for i = 1:length(T)
        Ti = T(i);
        power = A .* Ti;
        E = expm(power);
        Btime = E * B0;
        Bi(:,i) = Btime;
    end

    % Return parameters in a complete vector
    parameters = [N G0 D0 Dn kF kR aG aD bG bD cG cD nC Vtot MM C0];
    
elseif strcmp(sol, 'numerical')
    
    %%%%%%%%%%%%%%%%%%%%%%
    % NUMERICAL SOLUTION %
    %%%%%%%%%%%%%%%%%%%%%%
    
    % NEED TO IMPLEMENT DIFFERENT DRUG ACTION SITUATIONS
    
    % Unpack parameters, correct the units
    ni    = kinParams(6);
    Vtot  = simParams(2);
    kf0   = kinParams(2);
    kf    = kf0/(Na*Vtot);
    kr    = kinParams(3);
    dmax  = kinParams(4);
    ic    = kinParams(5);
    r0    = kinParams(1);
    
    if Binit == 1
        Bin = Binit;
    elseif length(Binit) ~= 1
        Bin = 1;
    elseif Binit ~= 1
        Bin = 1;
    end

    Bin   = Bin * simParams(1);
    muf   = simParams(3);
    dt    = tstep;

    % Generate time vector
    tm    = min(trange):tstep:max(trange);
    tlen  = length(tm);

    % The variable x means the real number of occupied places
    XL    = ni;
    xmin  = 0;  
    xmax  = XL;
    xelem = ni+1;
    x     = linspace(xmin, xmax, xelem)';
    xlen  = length(x);

    % Molecules of drug
    A     = C0/muf*Na*Vtot;

    % Death and growth functions
    r             = r0*(1-x(:)/ni);
    dd             = zeros(xlen,1);
    dd((ic+1):end) = dmax;
    % Add 1 to iC because INDEX for iC is one more than the defined iC

    % Allocate memory
    B     = zeros(ni+1,tlen);
    Ball  = zeros(1,tlen);
    Pb    = zeros(xlen,xlen); % first index > second! q=h:xlen

    % Calculate binomial coefficients using bfun
    for hi = 1:xlen
        for hj = hi:xlen
            % Before i number of targets and then j from i to ni
            Pb(hi,hj) = bfun(x(hj),x(hi));
        end
    end

    % Iterate through time
    B(1,1) = Bin;
    errorcounter = 0;

    for n = 1:tlen-1

        % Check stability
        mm = max([dt*kf*A*ni, dt*r0, dt*dmax, dt*kr*ni]);
        if mm > 0.5
            warning('Stability error. Results may be inaccurate.')
            errorcounter = errorcounter + 1;
        else
        end

        B(1,n+1) = B(1,n) -...
                   dt*kf*A*ni*B(1,n) +...
                   dt*kr*x(2)*B(2,n) -...
                   dt*r(1)*B(1,n) -...
                   dt*dd(1)*B(1,n) +...
                   dt*(Pb(1,:)*(r(:).*B(:,n)));

        B(2:xlen-1,n+1) = B(2:xlen-1,n) +...
                          dt*kf*A*(ni-x(1:xlen-2)).*B(1:xlen-2,n) -...
                          dt*kf*A*(ni-x(2:xlen-1)).*B(2:xlen-1,n) -...
                          dt*kr*x(2:xlen-1).*B(2:xlen-1,n) +...
                          dt*kr*x(3:xlen).*B(3:xlen,n) -...
                          dt*dd(2:xlen-1).*B(2:xlen-1,n) -...
                          dt*r(2:xlen-1).*B(2:xlen-1,n) +...
                          dt*(Pb(2:xlen-1,:)*(r(:).*B(:,n)));

        B(xlen,n+1) = B(xlen,n) -...
                      dt*dd(xlen)*B(xlen,n) -...
                      dt*kr*x(xlen)*B(xlen,n) -...
                      dt*r(xlen)*B(xlen,n) +...
                      dt*kf*(ni-x(xlen-1))*B(xlen-1,n) +...
                      dt*(Pb(xlen,:)*(r(:).*B(:,n)));

        Ball(n+1) = sum(B(:,n+1));

    end

    Bi = B;
    % Return parameters in a complete vector
    parameters = [N G0 D0 Dn kF kR aG aD bG bD cG Cd nC Vtot MM C0];
    
end

runtime = toc;

end
```
