## Supplementary data file 1 for "Mechanisms of antibiotic action shape the fitness landscapes of resistance mutations": ABkineticExt9P.m

```
% Modified ABkinetic9P for finding time to extinction

% Colin Hemez
% 2019

function B = ABkineticExt9P(kinParams,...
                            simParams,...
                            Binit,...
                            C0,...
                            T,...
                            ext,...
                            varargin)

Bi = ABkinetic9P(kinParams,simParams,Binit,C0,T,varargin);
Bsum = sum(Bi);

B = Bsum - ext;
                        
end
```
