## Supplementary data file 1 for "Mechanisms of antibiotic action shape the fitness landscapes of resistance mutations": ABkineticMinimize9P.m

```
% Optimization of parameters for linear ODE model

% Model by
% Pia Abel zur Wiesch
% Leonid Chindelevitch
% Colin Hemez

% Implemented by
% Colin Hemez
% 2019

% INPUTS
% problem: Struct with the following fields
%          experiment: NxM matrix of experimental data
%          timevals: Mx1 vector of experimental timepoints [sec]
%          concvals: Nx1 vector of experimental concentrations [g/l]
%          kinParams: 12-value parameter vector
%          simParams: 3-value simulation vector
%          paramLB: lower bounds for all parameters
%          paramUB: upper bounds for all parameters
%          strategy: specifies which kinetic parameters to optimize
%          scoringMatrix: NxM matrix for weighting minimization function
%          hybrid: specifies whether to use hybrid optimization method

% OUTPUTS
% objectiveProblem: Struct that is passed into the solver
% solution: Struct containing the results of the optimization problem

% Additional functions needed in directory
% ABKINETICOBJECTIVE9P()
% PLOTKINETIC9P()
% ABKINETIC9P()
% SYSTEMMATRICES9P()
% BFUN()
% RATEFUN()
% GROWTHLIMIT()
% DEATHLIMIT()

function [solution, objectiveProblem] = ABkineticMinimize9P(problem)

% Unpack problem struct
expdata  = problem.experiment;
time     = problem.timevals;
drug     = problem.concvals;
K        = problem.kinParams;
S        = problem.simParams;
LB       = problem.paramLB;
UB       = problem.paramUB;
strategy = problem.strategy;
score    = problem.scoringMatrix;
H        = problem.hybrid;
SA       = problem.annealingParams;

%%%%%%%%%%%%%%%%%%%%%%%%%%%%%%%%%%%%%%%
% DETERMINE THE OPTIMIZATION STRATEGY %
%%%%%%%%%%%%%%%%%%%%%%%%%%%%%%%%%%%%%%%

if strcmp(strategy, '8param1')
    % 8-parameter minimization: kF, kR, aG, aD, bG, bD, cG, cD
    optIndex = 5:12;
elseif strcmp(strategy, '2param1')
    % 2-parameter minimization: cG, cD
    optIndex = 11:12;
elseif strcmp(strategy, '7param1')
    % 7-parameter minimization: kF, kR, aG, aD, bG, cG, cD
    optIndex = [5 6 7 8 9 11 12];
elseif strcmp(strategy, '6param1')
    % 6-parameter minimization: kF, kR, aG, aD, cG, cD
    optIndex = [5 6 7 8 11 12];
elseif strcmp(strategy, '4param1')
    % 4-parameter minimization: aG, aD, cG, cD
    optIndex = [7 8 11 12];
elseif strcmp(strategy, '2param2')
    % 4-parameter minimization: kF, kR
    optIndex = [5 6];
end

optK = K(optIndex); % Define knietic parameters to optimize
conIndex = setdiff(1:12, optIndex);
conK = K(conIndex); % Define kinetic parameters that will remain constant

% Extract upper and lower bounds for parameters to optimize
objLB = LB(optIndex);
objUB = UB(optIndex);

%%%%%%%%%%%%%%%%%%%%%%%%%%%%%%%
% SET UP MINIMIZATION PROBLEM %
%%%%%%%%%%%%%%%%%%%%%%%%%%%%%%%

% Package additional information into a struct for ABkineticObjective9P
objectiveProblem.experiment     = expdata;
objectiveProblem.timevals       = time;
objectiveProblem.concvals       = drug;
objectiveProblem.simParams      = S;
objectiveProblem.scoringMatrix  = score;
objectiveProblem.optimIndex     = optIndex;
objectiveProblem.constantIndex  = conIndex;
objectiveProblem.optimParams    = optK;
objectiveProblem.constantParams = conK;

% Declare anonymous function
OBJ = @(optK)ABkineticObjective9P(optK,conK,objectiveProblem);

% Set options for the simulated annealing algorithm
iterations = SA(1);
itemp = SA(2);
reanneal = SA(3);

if H == 1
    % Set options for the local search algorithm
    hybridoptions = optimoptions(@fmincon,...
                                 'Display','iter',...
                                 'Algorithm','interior-point',...
                                 'PlotFcn',...
                                 {@optimplotx,@optimplotfval},...
                                 'TypicalX',optK,...
                                 'MaxIterations',SA(4));
    options = optimoptions(@simulannealbnd,...
                           'PlotFcn',...
                           {@saplottemperature,@saplotx,...
                            @saplotf,@saplotbestf},...
                           'MaxIterations',iterations,...
                           'HybridFcn',{@fmincon,hybridoptions},...
                           'InitialTemperature',itemp,...
                           'ReannealInterval',reanneal,...
                           'AnnealingFcn',@scaleFreeAnnealFcn9P,...
                           'TemperatureFcn',@temperatureexp);
else
    % No local search; set options for SA only
    options = optimoptions(@simulannealbnd,...
                           'PlotFcn',...
                           {@saplottemperature,@saplotx,...
                            @saplotf,@saplotbestf},...
                           'MaxIterations',iterations,...
                           'InitialTemperature',itemp,...
                           'ReannealInterval',reanneal,...
                           'AnnealingFcn',@scaleFreeAnnealFcn9P,...
                           'TemperatureFcn',@temperatureexp);
end

%%%%%%%%%%%%
% MINIMIZE %
%%%%%%%%%%%%

% Optimize via simulated annealing
[S,fval,terminal,process] = simulannealbnd(OBJ,optK,objLB,objUB,options);

% Repackage optimized kinetic parameters
Kfit = zeros(1, length(K));
Kfit(optIndex) = S;
Kfit(conIndex) = conK;

% Generate a plot of the model fit against experimental data
[H,R2] = plotKinetic9P(expdata,drug,time,Kfit,objectiveProblem.simParams);

% Output results
solution.paramopt   = S;
solution.paramset   = Kfit;
solution.fvalue     = fval;
solution.exit       = terminal;
solution.output     = process;
solution.plothandle = H;
solution.pearson    = R2;
end
```
