## Supplementary data file 1 for "Mechanisms of antibiotic action shape the fitness landscapes of resistance mutations": ABkineticObjective9P.m

```
% ABkinetic objective function
% For fitting antibiotic ODE model to experimental data

% Model by
% Pia Abel zur Wiesch
% Leonid Chindelevitch
% Colin Hemez

% Implemented by
% Colin Hemez
% 2019

% Objective function minimizes the sum of the squares of the normalized
% differences between experimental and simulated data

% Additional functions needed in directory
% ABKINETIC9P()
% SYSTEMMATRICES9P()
% BFUN()
% RATEFUN()
% GROWTHLIMIT()
% DEATHLIMIT()

function [diff, diffMat] = ABkineticObjective9P(optK,conK,problem)

% Unpack information from the problem struct
optI  = problem.optimIndex;
conI  = problem.constantIndex;
E     = problem.experiment;
time  = problem.timevals;
drug  = problem.concvals;
S     = problem.simParams;
score = problem.scoringMatrix;

% Construct kinetic parameter array
Kcon = zeros(1, (length(optI)+length(conI)));
Kopt = Kcon;
Kcon(optI) = optK;
Kopt(conI) = conK;
K = Kcon + Kopt;

% Prepare for simulations
simB = zeros(length(drug), length(time));
B0 = 1;

% Run simulations using given concentration and time values
for i = 1:length(drug)
    C0 = drug(i);
    Bi = ABkinetic9P(K, S, B0, C0, time, 'curve', 'exact');
    B = sum(Bi);
    simB(i,:) = B;
end

% Calculate the difference between experiment and simulation
simDiff = simB - E;

% Scale the differences relative to experimental values
scaledSimDiff = simDiff ./ E;

% Score the difference according to scoring matrix
scaledSimDiff = scaledSimDiff .* score;

% Calculate the square of the scaled differences, sum, and normalize
diffMat = scaledSimDiff .^ 2;
dim = size(diffMat);
diff = sum(sum(diffMat))/(dim(1) * dim(2));
end
```
