## Supplementary data file 1 for "Mechanisms of antibiotic action shape the fitness landscapes of resistance mutations": bfun.m

```
% Binomial related function
% q and h are the number of targets, not the indices
% 2* q!/(h!*(q-h)!) /2^q

function bf = bfun(q,h)

bf = 2*exp(gammaln(q+1)-gammaln(h+1)-gammaln(q-h+1))/2^q; % choosek(j,i) => choosek(q,h)

end
```
