## Supplementary data file 1 for "Mechanisms of antibiotic action shape the fitness landscapes of resistance mutations": bindingConstant1.m

```
% Investigate the effect of initial population size on compensation prob

% Colin Hemez
% 2019

clear;
close all;

X = getMutParams9P('ecoli','cipP1','infect9','curve');

% Initial population sizes
Mi = [X.Kres(5)*5 X.Kres(5)*3 X.Kres(5) X.Kres(5)/3 X.Kres(5)/5];

% Resolution for simulations
res = 120;

% Define parameter to vary for probability calculations
Ptarget = 'C0';
Pvals = linspace(X.MICres/2, X.MICres*10, res);

% Calculate probabilities
probvals = zeros(res, length(Mi));

for i = 1:length(Mi)
    % Redefine binding constant
    X.Kres(5) = Mi(i);
    X.MSB = mutSelBal9P(X.Kres, X.S, X.M, X.R);
    
    Pi = probabilityCurve9P(X.Kres,...
                            X.S,...
                            X.MSB,...
                            X.M(2),...
                            X.MICres,...
                            Ptarget,...
                            Pvals,...
                            'curve', 'exact');
                        
    probvals(:,i) = Pi;
end

% Plot
figure
plot(Pvals./(X.MICres), probvals, 'linewidth', 3);
grid on
xlabel('[DRUG] / MIC_{RESISTANT}')
ylabel('Probability of compensation emergence')
legend(num2str(Mi(1)),...
       num2str(Mi(2)),...
       num2str(Mi(3)),...
       num2str(Mi(4)),...
       num2str(Mi(5)),...
       'location','northeast')
title('Probabilities depend strongly on mutation rate')
```
