## Supplementary data file 1 for "Mechanisms of antibiotic action shape the fitness landscapes of resistance mutations": cellSubPop.m

```
% Cell subpopulation distribution

% Colin Hemez
% 2019

function P = cellSubPop(K, S, Binit, C0, T, plotter, varargin)

% Run the ABkinetic simulation
if isempty(varargin)
    Bi = ABkinetic9P(K, S, Binit, C0, T);
elseif length(varargin) == 1
    Bi = ABkinetic9P(K, S, Binit, C0, T, varargin{1});
else
    Bi = ABkinetic9P(K, S, Binit, C0, T, varargin{1}, varargin{2});
end

Bt = Bi(:,end);
Btot = sum(Bt);
Bpop = Bt ./ Btot;

if plotter
    figure
    plot(0:1:K(1), Bpop, 'ok', 'markerfacecolor', [0 0 0], 'markersize', 8)
    grid on
    xlim([0 K(1)])
end

P = Bpop;

end
```
