## Supplementary data file 1 for "Mechanisms of antibiotic action shape the fitness landscapes of resistance mutations": compensationProb9P.m

```
% Calculate drug concentration for a given probability of compensation

% Colin Hemez
% 2019

function C = compensationProb9P(kinParams,...
                                simParams,...
                                mutParams,...
                                MSB,...
                                prob,...
                                tolerance,...
                                varargin)

% Calculate the MIC and mutation selection balance of the strain
MICres = findMIC9P(kinParams, 1e-12, 'curve');
MICres = MICres * simParams(3);

mu = mutParams(2);
ABres = 100;

% Find an upper bound for the drug concentration
UBfactor = 10;
currC = MICres * UBfactor;
[~,~,currP] = expectedMutants9P(kinParams,...
                                simParams,...
                                MSB,...
                                mu,...
                                currC,...
                                ABres,...
                                varargin);
while currP > prob
    currC = currC * UBfactor;
    [~,~,currP] = expectedMutants9P(kinParams,...
                                    simParams,...
                                    MSB,...
                                    mu,...
                                    currC,...
                                    ABres,...
                                    varargin);
end

% Define bounds
UB = currC;
LB = UB / UBfactor;
bounds = logspace(log10(LB), log10(UB), 3);
currC = bounds(2);
[~,~,currP] = expectedMutants9P(kinParams,...
                                simParams,...
                                MSB,...
                                mu,...
                                currC,...
                                ABres,...
                                varargin);

while abs(currP - prob) >= tolerance
    
    if currP > prob % Current concentration is too low
        LB = currC;
    else % Current concentration is too high
        UB = currC;
    end
    
    bounds = logspace(log10(LB), log10(UB), 3);
    currC = bounds(2);
    
    % Recalculate probability
    [~,~,currP] = expectedMutants9P(kinParams,...
                                    simParams,...
                                    MSB,...
                                    mu,...
                                    currC,...
                                    ABres,...
                                    varargin);
    
end

C = currC;

end
```
