## Supplementary data file 1 for "Mechanisms of antibiotic action shape the fitness landscapes of resistance mutations": drugMechanism1.m

```
% Investigate the effect of initial population size on compensation prob

% Colin Hemez
% 2019

clear;
close all;

mech = {'curve','nstat','ncide','cstat','ccide'};
X = getMutParams9P('ecoli','cipP1','infect9',mech{1});
MICfactor = 2;

% Look at steady-state distributions of bound targets
SS = zeros((X.Kwt(1) + 1), length(mech));
disPop = 10000;
Gdist = zeros(disPop, length(mech));
Ddist = Gdist;
Gmeans = zeros(length(mech), 1);
Dmeans = Gmeans;
Gstd = Gmeans;
Dstd = Gmeans;

for i = 1:length(mech)
    
    % Retrieve parameters
    X = getMutParams9P('ecoli','cipP1','infect9',mech{i});
    C = X.MICres * MICfactor;
    
    % Find the steady-state distribution
    SSi = SSDistribution(X.Kres, X.S, 1, C, 0, 'curve', 'exact');
    
    % Store distributions in memory
    SS(:,i) = SSi;
    
    % Retrieve growth and death curves for the mechanism
    [~,~,G,D] = systemMatrices9P(X.Kres, 0, 0);
    
    % Determine steady-state growth and death rate distributions
    Gvalsi = zeros(disPop, 1);
    Dvalsi = Gvalsi;
    fillInd = 1;
    for j = 1:(X.Kres(1) + 1)
        Gj = G(j);
        Dj = D(j);
        Nj = round(SSi(j) * disPop);
        Gvalsi(fillInd:(fillInd+Nj)) = Gj;
        Dvalsi(fillInd:(fillInd+Nj)) = Dj;
        fillInd = fillInd + Nj;
    end
    
    Gdist(:,i) = Gvalsi(1:disPop);
    Ddist(:,i) = Dvalsi(1:disPop);
    
    Gmeani = mean(Gvalsi(1:disPop));
    Dmeani = mean(Dvalsi(1:disPop));
    Gstdi  = std(Gvalsi(1:disPop));
    Dstdi  = std(Dvalsi(1:disPop));
    
    Gmeans(i) = Gmeani;
    Dmeans(i) = Dmeani;
    Gstd(i)   = Gstdi;
    Dstd(i)   = Dstdi;
    
end

% Plot steady-state distributions of bound targets
figure
plot(0:X.Kwt(1), SS, 'linewidth', 3)
grid on
xlabel('Number of bound targets')
ylabel('Proportion of steady-state cell population')
legend(upper(mech{1}),...
       upper(mech{2}),...
       upper(mech{3}),...
       upper(mech{4}),...
       upper(mech{5}),...
       'location','northwest')
title(['Drug-target distributions of steady-state populations at ',...
      num2str(MICfactor),...
      'x MIC'])
xlim([0 X.Kwt(1)])

% % Plot steady-state distributions of growth and death rates
% figure
% nedges = linspace(0, X.Kres(2), 50);
% subplot(2,1,1)
% histogram(Gdist(:,1), nedges)
% hold on
% histogram(Gdist(:,2), nedges)
% histogram(Gdist(:,4), nedges)
% legend(upper(mech{1}),...
%        upper(mech{2}),...
%        upper(mech{4}),...
%        'location','northeast')
% 
% nedges = linspace(0, X.Kres(4), 50);
% subplot(2,1,2)
% histogram(Ddist(:,1), nedges)
% hold on
% histogram(Ddist(:,3), nedges)
% histogram(Ddist(:,5), nedges)
% legend(upper(mech{1}),...
%        upper(mech{3}),...
%        upper(mech{5}),...
%        'location','northwest')

% Plot mean normalized growth and death rates for all mechanisms
figure
subplot(2,1,1)
bar((Gmeans./X.Kres(2)))
subplot(2,1,2)
bar((Dmeans./X.Kres(3)))

%% Resolution for probability simulations
res = 10;

% Define target parameter
Ptarget = 'C0';

% Set aside memory
probvals = zeros(res, length(mech));

for i = 1:length(mech)
    
    % Retrieve parameters
    X = getMutParams9P('ecoli','cipP1','infect9',mech{i});
    
    % Define parameter values
    Pvals = linspace(X.MICres*1, X.MICres*70, res);
    
    % Calculate probabilities
    Pi = probabilityCurve9P(X.Kres,...
                            X.S,...
                            X.MSB,...
                            X.M(2),...
                            X.MICres,...
                            Ptarget,...
                            Pvals,...
                            mech{i}, 'exact');
    
    probvals(:,i) = Pi;
end

% Plot
figure
plot(Pvals./(X.MICres), probvals, 'linewidth', 3);
grid on
xlabel('[DRUG] / MIC_{RESISTANT}')
ylabel('Probability of compensation emergence')
legend(upper(mech{1}),...
       upper(mech{2}),...
       upper(mech{3}),...
       upper(mech{4}),...
       upper(mech{5}),...
       'location','northeast')
title('Probabilities depend on drug action mechanism')

% Export probability values and drug concentrations
csvwrite('plotMechanismDrugConc.csv', (Pvals./(X.MICres))')
csvwrite('plotMechanismProbabilities.csv', probvals)
```
