## Supplementary data file 1 for "Mechanisms of antibiotic action shape the fitness landscapes of resistance mutations": estimateGrowthDeath9.m

```
% Estimate growth and drug-induced death rates from experimental data
% Data from Adam Palmer

% Assumptions for data fitting:
% For time series {B(t) | (C=0)}, all cells have 0 bound targets
% For time series (B(t) | (C=Cmax)}, all cells have 100% bound targets
% As a result of assumptions, population dynamics follow exponential curve

clear; close all;

% Load data
load('Cipro_best_fit_Adam.mat')
noDrug = load('cipro_noDrug_Adam.csv');
noDrug = noDrug .* 10000;

Czero = noDrug;
Cmax = Cexp7; % Population curve at maximum antibiotic concentration
B0 = Cexp7(1); % Initial population size

% Declare initial guesses for growth and death rates
Rguess = 2e-4;  % [1/sec]
Dguess = -1e-3; % [1/sec]

% Natural death rate
Dn = -1.5e-6;

% Create anonymous function
fun = @(R,texp)expFun(R,Dn,B0,texp);

% Solve
[Dfit, R2D, residualD] = lsqcurvefit(fun, Dguess, texp, Cmax);
[Rfit, R2R, residualR] = lsqcurvefit(fun, Rguess, texp, Czero);
Dfitt = expFun(Dfit, Dn, B0, texp); % Simulate with experimental timepoints
Rfitt = expFun(Rfit, Dn, B0, texp); % Simulate with experimental timepoints

% Render simulated data
tsim = linspace(min(texp), max(texp), 100);
Dsim = expFun(Dfit, Dn, B0, tsim);
Rsim = expFun(Rfit, Dn, B0, tsim);

% Plot
figure
semilogy(tsim, Dsim, '-k', 'linewidth', 4)
hold on
semilogy(texp, Cmax, 'or', 'markerfacecolor',[1 0 0])
grid on

figure
semilogy(tsim, Rsim, '-k', 'linewidth', 4)
hold on
semilogy(texp, Czero, 'ob', 'markerfacecolor', [0 0 1])
grid on

% Export simulated data
csvwrite('DeathSimulation.csv', [(tsim./60)' Dsim'])
csvwrite('DeathExperiment.csv', [(texp./60) Cexp7 Dfitt])
csvwrite('GrowthSimulation.csv', [(tsim./60)' Rsim'])
csvwrite('GrowthExperiment.csv', [(texp./60) Czero Rfitt])
```
