## Supplementary data file 1 for "Mechanisms of antibiotic action shape the fitness landscapes of resistance mutations": expectedMutants9P.m

```
% Calculate the expected number of mutants to arise before extinction

% Colin Hemez
% 2019

function [Y,Yi,P] = expectedMutants9P(kinParams,...
                                      simParams,...
                                      N0,...
                                      mu,...
                                      C,...
                                      resolution,...
                                      varargin)

% Package initial population size into simulation parameters vector
simParams(1) = N0;

% Define other inputs for ABKinetic
Binit = 1;

% Calculate time to extinction
Text = extinctionTime9P(kinParams,simParams,Binit,C,varargin);
if isnan(Text)
    Y = NaN;
    Yi = NaN(kinParams(1), 1);
    P = 1;
    return
end

% Define the time vector for ABkinetic simulation
T = logspace(0, log10(Text), resolution);

% Run the ABkinetic simulation
Bi = ABkinetic9P(kinParams,simParams,Binit,C,T,varargin);

% Repackage parameter vector using the growth and death rate functions
matrixParams = kinParams;

% Retrieve growth rates as a function of bound target
[~, ~, G, ~] = systemMatrices9P(matrixParams, 0, 0);

% Multiply growth rates by subpopulation sizes and integrate
BiGi = G' .* Bi;
I = trapz(T',BiGi,2);

% Factor in the mutation rate
Yi = I .* mu;

% Sum to obtain the total number of expected mutants
Y = sum(Yi);

% Calculate the Poisson probability that a mutant arises before extinction
P = 1 - exp(-Y);
                           
end
```
