## Supplementary data file 1 for "Mechanisms of antibiotic action shape the fitness landscapes of resistance mutations": expFun.m

```
% Exponential growth and decay fitting function
% For fitting exponential growth curves

% Allows for the explicit declaration of a natural death or growth rate Dn

function B = expFun(R, Dn, B0, t)

B = B0 .* exp((R + Dn) .* t);

end
```
