## Supplementary data file 1 for "Mechanisms of antibiotic action shape the fitness landscapes of resistance mutations": extinctionTime9P.m

```
% Find the time to extinction (Btotal =< 1) for a bacterial population

% Colin Hemez
% 2019

% Function returns an error if the C0 < MIC

function Text = extinctionTime9P(kinParams,simParams,Binit,C0,varargin)

% Check to make sure C0 > MIC
MIC = findMIC9P(kinParams, 1e-12, 'curve');
MIC = MIC * simParams(3);
if MIC >= C0
    warning('C0 given is at or below MIC; extinction time does not exist')
    Text = NaN;
    return
end

% Define extinction criterion
ext = 1;

% Generate the anonymous function
zeroFun = @(T)ABkineticExt9P(kinParams,simParams,Binit,C0,T,ext,varargin);

% Determine the range of time values to pass into zero finder
LB = 0;
UB = 1;
fval = ABkineticExt9P(kinParams,simParams,Binit,C0,UB,ext,varargin);

while fval > 0
    LB = UB;
    UB = UB * 10;
    fval = ABkineticExt9P(kinParams,simParams,Binit,C0,UB,ext,varargin);
end

guess = [LB UB];

% Find the extinction time
options = optimset('display', 'off');
Text = fzero(zeroFun, guess, options);

end
```
