## Supplementary data file 1 for "Mechanisms of antibiotic action shape the fitness landscapes of resistance mutations": findHill9P.m

```
% Find the Hill coefficient that best fits a PD curve

% Colin Hemez
% 2019

% Inputs
% pdCurve: Net growth rate values from the experimental PD curve
% C: Concentration range for PD curve, as multiples of MICwt

function [Hfit,R2] = findHill9P(pdCurve, params, concvals)

% Define MIC
MIC = params.MICres / params.MICwt;

% Find maximum growth and death rates
[~,~,G,D] = systemMatrices9P(params.Kres, 0, 0);
Gmax = max(G);
Dmax = max(D);

% Set initial guess
H0 = 2;
LB = H0 / 1000;
UB = H0 * 1000;

% Create anonymous function
EmaxAnon = @(H,Crange)pdEmax(H,MIC,Gmax,Dmax,Crange);

% Find the Hill coefficient
options = optimoptions('lsqcurvefit',...
                       'OptimalityTolerance',1e-10,...
                       'FunctionTolerance',1e-10);
Hfit = lsqcurvefit(EmaxAnon, H0, concvals, pdCurve, LB, UB, options);

% Make a plot
pdSim = pdEmax(Hfit,MIC,Gmax,Dmax,concvals);
figure
semilogx(concvals,pdCurve,concvals,pdSim,'linewidth',3)
grid on

% Calculate the R-squared value
corrvals = corrcoef(pdCurve, pdSim);
R2 = corrvals(2,1);

end
```
