## Supplementary data file 1 for "Mechanisms of antibiotic action shape the fitness landscapes of resistance mutations": findKdFactor9P.m

```
% Find the fold change in kF and kR values for a mutant strain

% Colin Hemez
% 2019

function [Q,Kres] = findKdFactor9P(Kwt, MICfold, cost)

% Define tolerance for MIC calculations
MICtol = 1e-12;
opttol = 2 * MICtol;

% Find the MIC of the wild-type
MICwt = findMIC9P(Kwt, MICtol, 'curve');
targMIC = MICwt * MICfold;

% Make an initial guess for the Q-factor
Qguess = MICfold / (1 - cost);
Qguess = sqrt(Qguess);

% Duplicate kinetic parameters vector
Kres = Kwt;
Kres(2) = Kres(2) * (1 - cost); % Incorporate cost into mutant growth rate

% Find the bounds for the Q-factor
Kres(5) = Kwt(5) / Qguess;
Kres(6) = Kwt(6) * Qguess;
Q = Qguess;

% Find the upper bounds for Q-factor
MICres = findMIC9P(Kres, MICtol, 'curve');
if MICres <= targMIC % Q-factor needs to be larger
    while MICres <= targMIC
        Q = Q * sqrt(2);
        Kres(5) = Kwt(5) / Q;
        Kres(6) = Kwt(6) * Q;
        MICres = findMIC9P(Kres, MICtol, 'curve');
    end
else % Q-factor needs to be smaller
    while MICres > targMIC
        Q = Q / sqrt(2);
        Kres(5) = Kwt(5) / Q;
        Kres(6) = Kwt(6) * Q;
        MICres = findMIC9P(Kres, MICtol, 'curve');
    end
    Q = Q * sqrt(2);
end

ubQ = Q;
lbQ = ubQ / sqrt(2);

% Find the value of the Q-factor within MIC tolerance
Qvals = logspace(log10(lbQ), log10(ubQ), 3);
Q = Qvals(2);
Kres(5) = Kwt(5) / Q;
Kres(6) = Kwt(6) * Q;
MICres = findMIC9P(Kres, MICtol, 'curve');
while (abs(MICres - targMIC)) > opttol
    if MICres < targMIC
        lbQ = Q;
        Qvals = logspace(log10(lbQ), log10(ubQ), 3);
        Q = Qvals(2);
    else
        ubQ = Q;
        Qvals = logspace(log10(lbQ), log10(ubQ), 3);
        Q = Qvals(2);
    end
    Kres(5) = Kwt(5) / Q;
    Kres(6) = Kwt(6) * Q;
    MICres = findMIC9P(Kres, MICtol, 'curve');
end

end
```
