## Supplementary data file 1 for "Mechanisms of antibiotic action shape the fitness landscapes of resistance mutations": findMIC9P.m

```
% MIC Finder
% Based on the model by
% Pia Abel zur Wiesch
% Leonid Chindelevitch
% Colin Hemez

% Implemented by
% Colin Hemez
% 2019

% INPUTS
% kinParams: Kinetic parameters
%            Vector of form [N G0 D0 Dn kF kR aG aD bG bD cG cD]
%            N  = number of target complexes per cell [//cell]
%            G0 = growth rate in the absence of antibiotic [1/sec]
%            D0 = death rate in the absence of antibiotic [1/sec]
%            Dn = death rate in saturating antibiotic concentration [1/sec]
%            kF = drug-target binding rate constant [1/M/sec]
%            kR = drug-target unbinding rate constant [1/sec]
%            aG = Sharpness of growth rate function G[i]
%            aD = Sharpness of death rate function D[i]
%            bG = Bacteriostatic potency of drug at high concentrations
%            bD = Bactericidal potency of drug at high concentrations
%            cG = Inflection point for growth rate function G[i]
%            cD = Inflection point for death rate function D[i]
% tolerance: margin of error for finding MIC (1e-10 is usually good)
% mechanism: growth and death rate functions G[i] and D[i]
%            'general': ciprofloxacin parameter fit to PDE model
%            'cipro': ciprofloxacin parameter fit to ODE model
%            'nstat': stabilization/bacteriostatic
%            'ncide': stabilization/bactericidal
%            'cstat': competition/bacteriostatic
%            'ccide': competition/bactericidal
%            'curve': use parameters supplied in kinParams array

% Optional imput:
% dialogue: enable or suppress dialogue
%           1 = turns on function dialogue
%           0 = suppresses dialogue
%           If no input is given, dialogue is suppressed

% Additional functions needed in directory
% SYSTEMMATRICES9P()
% BFUN()
% RATEFUN()
% GROWTHLIMIT()
% DEATHLIMIT()

function MIC = findMIC9P(kinParams, tolerance, mechanism, varargin)
% Check for a correct number and type of input arguments
if nargin < 2
    error('Not enough input arguments')
elseif nargin > 4
    error('Too many input arguments')
end

% Verify class of PARAMS object
if length(kinParams) ~= 12
    error('PARAMS must be a 12-value parameter vector')
end

% Verify class of TOLERANCE object
if length(tolerance) ~= 1
    error('TOLERANCE must be a single value')
end

% Unpack needed variables
kF = kinParams(5); % Convert rate to 1/hr if needed
tol = tolerance;

% Determine matrix construction method and dialogue options
defaultdia = 0;

if nargin == 3
    situation = mechanism;
    dialog = defaultdia;
elseif nargin == 4
    if (varargin{1} == 1) || (varargin{1} == 0)
        situation = mechanism;
        dialog = varargin{1};
    else
        warning('Invalid input for DIALOGUE; suppressing all dialogue')
        situation = mechanism;
        dialog = 0;
    end
end

% Unpack needed parameters
kF = kinParams(5);
aG = kinParams(7);
aD = kinParams(8);
bG = kinParams(9);
bD = kinParams(10);
cG = kinParams(11);
cD = kinParams(12);

% Define growth and death function parameters
if strcmp(situation, 'curve') || strcmp(situation, 'cipro')
    GDfunction = [aG aD bG bD cG cD];
elseif strcmp(situation, 'general')
    GDfunction = [1 500 0 1 cG cD];
elseif strcmp(situation,'cipro')
    GDfunction = [1 500 0 1 cG cD];
elseif strcmp(situation, 'nstat')
    GDfunction = [500 1 0 0 cG cD];
elseif strcmp(situation, 'ncide')
    GDfunction = [1 500 1 1 cG cD];
elseif strcmp(situation, 'cstat')
    GDfunction = [1 1 0 0 cG cD];
elseif strcmp(situation, 'ccide')
    GDfunction = [1 1 1 1 cG cD];
else
    warning('Invalid input for MECHANISM; using values from KINPARAMS')
    GDfunction = [aG aD bG bD cG cD];
end

% Rebuild parameter vector
kinParams = [kinParams(1:6) GDfunction];

%%%%%%%%%%%%%%%%%%%%%%%%%%%%%%%%%%%%%%%%%%%%%
% BUILD MATRIX REPRESENTATION OF ODE SYSTEM %
%%%%%%%%%%%%%%%%%%%%%%%%%%%%%%%%%%%%%%%%%%%%%
[A0, A1] = systemMatrices9P(kinParams, 0, 0);

% Determine upper bound for MIC by log iteration
eigenval = 0;
currentC = 0;
while eigenval >=0
    if currentC == 0
        currentC = 1e-8;
    else
        currentC = currentC * 10;
    end
    MICmat = A1 + (kF .* currentC .* A0);
    eigenval = max(real(eig(MICmat)));
end

% Search for MIC within tolerance given a known upper bound
UB = currentC;
if UB == 1
    LB = 0;
else
    LB = UB/10;
end

if dialog
    disp(' ')
    disp(['MIC upper bound (mol/L): ', num2str(UB)])
    disp(['MIC lower bound (mol/L): ', num2str(LB)])
    disp(' ')
    disp('Searching for MIC value...')
    disp(' ')
end

iteration = 1; % Iteration counter (for dialogue if desired)

while (UB-LB) >= tol
    currentC = sqrt(UB * LB);
    MICmat = A1 + (kF .* currentC .* A0);
    eigenval = max(real(eig(MICmat)));
    if real(eigenval) >= 0
        LB = currentC;
    else
        UB = currentC;
    end
    
    if dialog
        disp(['Iteration...................', num2str(iteration)])
        disp(['Current MIC value (mol/L)...', num2str(currentC)])
        disp(['Eigenvalue..................', num2str(eigenval)])
        disp(' ')
    end
    
    iteration = iteration + 1;
end

MIC = currentC;

end
```
