## Supplementary data file 1 for "Mechanisms of antibiotic action shape the fitness landscapes of resistance mutations": findSSDecayTime.m

```
% Find the time at which population decay becomes quasi-exponential

% Colin Hemez
% 2019

% Concentration of drug given (C0) MUST be above MIC

% INPUTS
% kinParams: Kinetic parameters
%            Vector of form [N G0 D0 Dn kF kR aG aD bG bD cG cD]
%            N  = number of target complexes per cell [1/cell]
%            G0 = growth rate in the absence of antibiotic [1/sec]
%            D0 = death rate in the absence of antibiotic [1/sec]
%            Dn = death rate in saturating antibiotic concentration [1/sec]
%            kF = drug-target binding rate constant [1/M/sec]
%            kR = drug-target unbinding rate constant [1/sec]
%            aG = Sharpness of growth rate function G[i]
%            aD = Sharpness of death rate function D[i]
%            bG = Bacteriostatic potency of drug at high concentrations
%            bD = Bactericidal potency of drug at high concentrations
%            cG = Inflection point for growth rate function G[i]
%            cD = Inflection point for death rate function D[i]
% simParams: Simulation parameters
%            Vector of the form [nC Vtot MM]
%            nC   = initial number of cells in medium [dimensionless]
%            Vtot = total volume of the simulation [l]
%            MM   = molar mass of drug being simulated [g/mol]
% Binit:     Initial distribution of cells in each bound target category,
%            as a column vector of dimensions [N x 1]. Distributions
%            should be expressed as proportions of the total, such that
%            sum(Binit) = 1
% C0:        Concentration of drug [g/l]

% OPTIONAL ADDITIONAL INPUTS:
%  'mechanism' specifies predefined pair of growth and death rate functions
%   Acceptable inputs are
%    'general': Ciprofloxacin parameter fir to PDE model
%    'cipro': Ciprofloxacin parameter fit to ODE model
%    'nstat': Stabilization/bacteriostatic
%    'ncide': Stabilization/bactericidal
%    'cstat': Competition/bacteriostatic
%    'ccide': Competition/bactericidal
%    'curve': Use kinParams for growth and death rate function parameters
%    If a different input is given, 'curve' is used
% 'solution' specifies the ODE solver method
%   Acceptable inputs are
%    'exact': solve ODEs using MATLAB's matrix exponentiation algorithm
%    'numerical': solve ODEs by taking small time steps from t = 0

% Additional functions needed in directory
% SYSTEMMATRICES9P()
% BFUN()
% RATEFUN()
% GROWTHLIMIT()
% DEATHLIMIT()

function T = findSSDecayTime(kinParams, simParams, Binit, C0, varargin)

% Determine matrix construction method and solution method
defaultsit = 'curve';
defaultsol = 'exact';

if nargin == 4
    situation = defaultsit;
    sol    = defaultsol;

elseif nargin == 5
    if strcmp(varargin{1},'curve')
        situation = 'curve';
        sol    = defaultsol;
    elseif strcmp(varargin{1},'general')
        situation = 'general';
        sol    = defaultsol;
    elseif strcmp(varargin{1},'nstat')
        situation = 'nstat';
        sol    = defaultsol;
    elseif strcmp(varargin{1},'cstat')
        situation = 'cstat';
        sol    = defaultsol;
    elseif strcmp(varargin{1},'ncide')
        situation = 'ncide';
        sol    = defaultsol;
    elseif strcmp(varargin{1},'ccide')
        situation = 'ccide';
        sol    = defaultsol;
    elseif strcmp(varargin{1},'cipro')
        situation = 'cipro';
        sol    = defaultsol;
    elseif strcmp(varargin{1},'exact')
        sol    = 'exact';
        situation = defaultsit;
    elseif strcmp(varargin{1},'numerical')
        sol    = 'numerical';
        situation = defaultsit;
    else
        situation = defaultsit;
        sol    = defaultsol;
    end
    
elseif nargin == 6
    if strcmp(varargin{1},'exact') ||...
       strcmp(varargin{1},'numerical')
        sol = varargin{1};
        if strcmp(varargin{2},'general') ||...
           strcmp(varargin{2},'nstat') ||...
           strcmp(varargin{2},'cstat') ||...
           strcmp(varargin{2},'ncide') ||...
           strcmp(varargin{2},'ccide') ||...
           strcmp(varargin{2},'cipro')
            situation = varargin{2};
        else
            situation = defaultsit;
        end
    elseif strcmp(varargin{1},'general') ||...
           strcmp(varargin{1},'nstat') ||...
           strcmp(varargin{1},'cstat') ||...
           strcmp(varargin{1},'ncide') ||...
           strcmp(varargin{1},'ccide') ||...
           strcmp(varargin{1},'cipro')
        situation = varargin{1};
        if strcmp(varargin{2},'exact') ||...
           strcmp(varargin{2},'numerical')
            sol = varargin{2};
        else
            sol = defaultsol;
        end
    else
        situation = defaultsit;
        sol    = defaultsol;
    end
end

% Check that concentration of drug given is above MIC
MIC = findMIC9P(kinParams, 1e-12, situation);
MICm = MIC * simParams(3); % convert from mol/l to g/l
if C0 < MICm
    error(['[DRUG (',...
           num2str(C0),...
           ' g/l)] must be above MIC (',...
           num2str(MICm),...
           ' g/l)'])
end

% Set tolerance for linearity of log(population)
R2threshold = -0.9999;

% Find X-second time window at which population decay becomes exponential
Trange = 300;
numpoints = 10;
currT = linspace(0,Trange,numpoints) - Trange;
R2 = 0;

while R2 > R2threshold
    currT = currT + Trange;
    Bi = ABkinetic9P(kinParams,simParams,Binit,C0,currT,situation,sol);
    B = sum(Bi);
    logB = log10(B);
    Rmat = corrcoef(currT, logB);
    R2 = Rmat(1,2);
end

% Use midpoint search to find start point for exponential decay
UB = min(currT);
LB = UB - Trange;

while (UB - LB) > 1
    Tstart = (LB + UB) / 2;
    currT = linspace(Tstart, (Tstart+Trange), numpoints);
    Bi = ABkinetic9P(kinParams,simParams,Binit,C0,currT,situation,sol);
    B = sum(Bi);
    logB = log10(B);
    Rmat = corrcoef(currT, logB);
    R2 = Rmat(1,2);
    if R2 > R2threshold
        LB = Tstart;
    else
        UB = Tstart;
    end
end

T = currT(1);

end
```
