## Supplementary data file 1 for "Mechanisms of antibiotic action shape the fitness landscapes of resistance mutations": fitnessCostEffect9P.m

```
% Find MIC as a function of fitness cost for different drug mechanisms

% Colin Hemez
% 2019

function [MICs,ratios,P] = fitnessCostEffect9P(MICfold,...
                                               commoncost,...
                                               costvals)

% Define the drug action mechanisms
mech = {'curve', 'nstat', 'cstat', 'ncide', 'ccide'};

% Set the tolerance for MIC calculations (1e-12 is usually good)
tol = 1e-12;

% Set aside memory
MICs = zeros(length(costvals), length(mech));
MICis = zeros(length(costvals), 1);
ratios = MICs;

% Calculate MICs for each drug action mechanism
for i = 1:length(mech)
    % Retrieve parameters
    mechi = mech{i};
    Xi = getMutParams9P('ecoli', 'cipP1', 'infect9', mechi);
    Kwti = Xi.Kwt;
    
    MICwt = findMIC9P(Kwti, tol, 'curve');
    
    % Determine kF and kR parameters for the resistant strain
    [~,Kresi] = findKdFactor9P(Kwti, MICfold, commoncost);
    
    % Calculate MIC's across the fitness cost range
    for j = 1:length(costvals)
        Kresi(2) = Kwti(2) * (1 - costvals(j));
        MICi = findMIC9P(Kresi, tol, 'curve');
        MICis(j) = MICi;
    end
    
    ratioi = MICis ./ MICwt;
    ratios(:,i) = ratioi;
    
end

% Export the ratios to a csv file
csvwrite('fitnessCostEffect9P_output.csv', ratios)

% Plot the results
figure
hold on
P = plot(costvals, ratios, 'linewidth', 3);
plot([min(costvals) max(costvals)], [MICfold MICfold], '--k')
plot([min(costvals) max(costvals)], [1 1], '--k')
plot([commoncost commoncost], [0 MICfold], '--k')
grid on
xlabel('Fitness cost')
ylabel('MIC_{MUTANT} / MIC_{WT}')
title('Ratios in MIC between mutant and wild-type')
legend(upper(mech{1}),...
       upper(mech{2}),...
       upper(mech{3}),...
       upper(mech{4}),...
       upper(mech{5}),...
       'location', 'northeast')

end
```
