## Supplementary data file 1 for "Mechanisms of antibiotic action shape the fitness landscapes of resistance mutations": fitnessFoldMod9P.m

```
% Fitness fold modification calculator

% Calculates the growth rate and fitness hit for a population exposed to
% antibiotic given a set of parameter values and a target MIC

% Inputs
% params:    wild-type kinetic parameters (from GETMUTPARAMS9P)
% targMIC:   target MIC value
% targParam: parameter to modify
%            n1: target molecule number, survival threshold
%            n2: target molecule number, toxicity threshold
%            n3: target molecule number, survival threshold for growth
%                                        toxicity threshold for death
% paramVals: array of target parameter values
% mode:      mechanism of antibiotic action (cstat, ncide, cipro, etc.)

% Outputs
% GR: array of growth rate values
% FH: array of fitness hit values, where FH(i) = -(GR(i)-GR(wt))/GR(wt)
% B:  matrix of bounds used to estimate each fitness hit
%     column 1 is the array of lower bounds for growth rate
%     column 2 is the array of upper bounds for growth rate

% Notes
% paramVals MUST be ordered such that paramVals(i+1) yields an MIC that is
% greater than paramVals(i) if all other parameters are kept constant

function [FH,GR,B] = fitnessFoldMod9P(kinParams,...
                                      foldMIC,...
                                      targParam,...
                                      paramVals,...
                                      mode)

% Unpack necessary parameters
r0 = kinParams(2);

% Calculate wild-type MIC
MICtol = 1e-12;
WTMIC = findMIC9P(kinParams, MICtol, mode);
targMIC = WTMIC * foldMIC;

% Determine which parameter is the target parameter
if strcmp(targParam, 'r0')
    error('growth rate (r0) cannot be used as target parameter')
elseif strcmp(targParam, 'kF')
    targIndex = 5;
elseif strcmp(targParam, 'kR')
    targIndex = 6;
elseif strcmp(targParam, 'dd')
    targIndex = 4;
elseif strcmp(targParam, 'dn')
    targIndex = 3;
elseif strcmp(targParam, 'iC')
    targIndex = 11;
elseif strcmp(targParam, 'n1') || strcmp(targParam, 'n2')
    targIndex = 1;
    toxG = kinParams(1) - kinParams(11);
    toxD = kinParams(1) - kinParams(12);
elseif strcmp(targParam, 'n3')
    targIndex = 1;
    toxG = kinParams(1) - kinParams(11);
end

tolerance = 1e-8; % tolerance for the growth rate value
fitnessMax = 1.25;
GRmax = r0*fitnessMax; % maximum growth rate allowable during search

% Set aside memory
GR = zeros(length(paramVals), 1);
bounds = zeros(length(paramVals), 2);

% Define the maximum number of iterations while searching for fitness hit
maxIterations = 100;
skipper = 0;

% Scaling factor, to improve lower bound est. in cases of low precision
LBscale = 0.8;

% Set skipper tolerance
skippertol = 1e-6;

tic

for i = 1:length(paramVals)
      
    if skipper
        GR(i:end) = GR(i-1);
        bounds(i:end,1) = 0;
        bounds(i:end,2) = GRmax;
        disp('Skipping remaining parameter values')
        break
    end
    
    % Reset parameters and bounds
    loopParams = kinParams;
    counter = 1;
    
    % Set the lower bound by extrapolating the nearby slope of the curve
    if i > 3 && ~strcmp(targParam, 'n3')
        if GR(i-3) < GRmax
            GR_LB = (2 * GR(i-1) - GR(i-2)) * LBscale;
            if GR_LB < r0 * 0.00001
                GR_LB = r0 * 0.00001;
            end
        else
            GR_LB = r0 * 0.00001;
        end
    else
        GR_LB = loopParams(3);
    end
    
    % Set the upper bound for the current iteration of the search
    if i == 1
        GR_UB = GRmax;
    elseif ~strcmp(targParam, 'n3')
        GR_UB = GR(i-1);
    else
        GR_UB = GRmax;
    end
    
    % Save upper and lower bounds
    if nargout >= 3
        bounds(i,1) = GR_LB;
        bounds(i,2) = GR_UB;
    end
    
    % Determine if fitness hit is above upper bound
    loopParams(2) = GRmax;
    loopParams(targIndex) = paramVals(i);
    if strcmp(targParam, 'n1')
        loopParams(11) = paramVals(i) - toxG;
        loopParams(12) = paramVals(i) - toxD;
    elseif strcmp(targParam, 'n3')
        loopParams(11) = paramVals(i) - toxG;
    end
    maxMICi = findMIC9P(loopParams, MICtol, mode);
    
    if maxMICi <= targMIC
        GR(i) = GRmax;
        
    else
        GRguess = (GR_UB + GR_LB)/2;
        loopParams(2) = GRguess;
        MICi = findMIC9P(loopParams, MICtol, mode);
        
        while abs(MICi - targMIC) >= tolerance
            counter = counter + 1;
            if counter > maxIterations
                warning('Maximum number of iterations reached')
                break
            end
            
            if MICi >= targMIC
                GR_UB = GRguess;
                GRguess = (GR_UB + GR_LB)/2;
            else
                GR_LB = GRguess;
                GRguess = (GR_UB + GR_LB)/2;
            end
            
            loopParams(2) = GRguess;
            MICi = findMIC9P(loopParams, MICtol, mode);
        end
        
        GR(i) = GRguess;
        
        if i > 1
            if (abs(((GR(i)-GR(i-1))/GR(i-1))) < skippertol) && (GR(i)<r0)
                skipper = 1;
            end
        end
        
    end
    
end

% Recalculate fitness hit just above upper threshold to get a smooth curve
disp('Smoothing out the lower part of the curve...')
counter = 1;
maximum = max(GR);
loopParams = kinParams;
maxIterations = 25;

if ~strcmp(targParam, 'n3')
    for i = 1:length(GR)
        if (GR(i) < maximum) && (i > 1)
            GR_LB = GR(i);
            GR_UB = GRmax * 10;
            GRguess = (GR_UB + GR_LB)/2;
            loopParams(2) = GRguess;
            loopParams(targIndex) = paramVals(i-1);
            MICi = findMIC9P(loopParams, MICtol, mode);

            while abs(MICi - targMIC) >= tolerance
                counter = counter + 1;
                if counter > maxIterations
                    warning('Maximum number of iterations reached')
                    break
                end

                if MICi >= targMIC
                    GR_UB = GRguess;
                    GRguess = (GR_UB + GR_LB)/2;
                else
                    GR_LB = GRguess;
                    GRguess = (GR_UB + GR_LB)/2;
                end

                loopParams(2) = GRguess;
                MICi = findMIC9P(loopParams, MICtol, mode);
            end

            GR(i-1) = GRguess;
            break
        end
    end
end

toc

FH = -1 .* (GR-r0) ./ r0;

B = bounds;

end
```
