## Supplementary data file 1 for "Mechanisms of antibiotic action shape the fitness landscapes of resistance mutations": fitnessIsoclimeFamily9P.m

```
% Generate a family of fitness isoclimes

% Colin Hemez
% 2019

function isoFam = fitnessIsoclimeFamily9P(foldMICs,...
                                          targParam,...
                                          foldParams)
                                   
% Define mechanisms
mech = {'cipro','nstat','cstat','ncide','ccide'};

% Set aside memory
isoclimes = zeros(length(foldParams), length(foldMICs));

% Go through each mechanism and foldMIC value to generate isoclimes
for i = 1:length(mech)
    
    % Retreive parameters
    Xi = getMutParams9P('ecoli', 'cipP1', 'infect9', mech{i});
    
    disp(' ')
    disp(['Mechanism: ', upper(mech{i})])
    
    % Generate target parameter values
    if strcmp(targParam, 'kF')
        paramVals = Xi.Kwt(5) ./ foldParams;
    elseif strcmp(targParam, 'kR')
        paramVals = Xi.Kwt(6) .* foldParams;
    elseif strcmp(targParam, 'n3')
        paramVals = foldParams;
    else
        error('Invalid input for TARGPARAM')
    end
    
    for j = 1:length(foldMICs)
        
        % Get the MIC fold value
        foldj = foldMICs(j);
        
        disp(['Evaluating parameter values for MIC fold ', num2str(foldj)])
        
        % Generate the isoclime
        isoclimej = fitnessFoldMod9P(Xi.Kwt,...
                                     foldj,...
                                     targParam,...
                                     paramVals,...
                                     'curve');
                                 
        % Store the resulting fitness hit values
        isoclimej = isoclimej(:);
        isoclimes(:,j) = isoclimej;
        
        disp(' ')
    end
    
    % Write the isoclime curve family into a csv file and a struct
    mechi = mech{i};
    csvwrite(['fitnessIsoclime_',targParam,'_',mechi,'.csv'],...
             isoclimes .* 100);
    isoFam.(mechi) = isoclimes;
end

isoFam.MICfolds = foldMICs;
isoFam.target = targParam;
isoFam.paramFoldVals = foldParams;

end
```
