## Supplementary data file 1 for "Mechanisms of antibiotic action shape the fitness landscapes of resistance mutations": fitnessTargetMod9P.m

```
% Fitness isoclimes for target number modulation
% Growth is on a survival threshold
% Death is on a toxicity threshld

% Colin Hemez
% 2018

function [FH, GR] = fitnessTargetMod9P(foldMICs,...
                                       paramVals,...
                                       varargin)

% Write some input files
csvwrite('fitnessIsoclime_n3_foldMICs.csv', foldMICs)
csvwrite('fitnessIsoclime_n3_paramVals.csv', paramVals)
                                   
% Define drug action mechanisms
mech = {'cipro','cstat','ncide','ccide','nstat'};

% Tolerance for MIC calculations
tol = 1e-12;
FHtol = 1e-3;
maxIterations = 16;

for i = 1:length(mech)

    % Pull out an initial set of parameters
    mechi = mech{i};
    X = getMutParams9P('ecoli','cipP1','infect9',mechi);
    Kwild = X.Kwt;
    currK = Kwild;
    r0 = Kwild(2);
    delG = Kwild(1) - Kwild(11);

    % Set aside memory
    GRi = zeros(length(paramVals), length(foldMICs));

    % Define the maximum growth rate to check
    maxFactor = 1.25;
    GRmax = r0 * maxFactor;
    
    disp(' ')
    disp(['MECHANISM: ', upper(mechi)])
    
    for j = 1:length(foldMICs)
        
        % Find the current target MIC
        targMIC = foldMICs(j) * X.MICwt / X.S(3);
        
        disp(['MIC-FACTOR: ', num2str(foldMICs(j)), ' ------------'])
        
        for k = 1:length(paramVals)
            Nk = paramVals(k);
            GGk = paramVals(k) - delG;
            currK(1) = Nk;
            currK(11) = GGk;
            
            disp(['TARGET NUMBER: ', num2str(Nk)])
            
            % Check to see if current parameters are outside fitness bounds
            currKmax = currK;
            currKmax(2) = GRmax;
            
            currMICmax = findMIC9P(currKmax, tol, 'curve');
            
            if currMICmax < targMIC
                % Outside bounds; don't calculate fitness hit
                currGR = GRmax;
                
            else
                % Inside bounds, calculate fitness hit
                iteration = 1;
                
                GRLB = 1.01 * currK(3);
                GRUB = GRmax;
                currGR = (GRLB + GRUB) / 2;
                
                currK(2) = currGR;
                currMIC = findMIC9P(currK, tol, 'curve');
                
                error = abs(currMIC - targMIC) / targMIC;
                
                while error > FHtol
                    
                    if iteration > maxIterations
                        warning('Maximum number of iterations reached')
                        break
                    end
                    
                    if currMIC > targMIC % currGR is to high
                        GRUB = currGR;
                        currGR = (GRLB + GRUB) / 2;
                    else
                        GRLB = currGR;
                        currGR = (GRLB + GRUB) / 2;
                    end
                    
                    currK(2) = currGR;
                    currMIC = findMIC9P(currK, tol, 'curve');
                    
                    error = abs(currMIC - targMIC) / targMIC;
                    
                    iteration = iteration + 1;
                    
                end
                
            end
            
            GRi(k,j) = currGR;
            
        end
        
    end

GR.(mechi) = GRi;
FH.(mechi) = -1 .* (GRi - r0) ./ r0;
csvwrite(['fitnessIsoclime_n3_',mechi,'.csv'], FH.(mechi).*100)

figure
plot(paramVals, FH.(mechi).*100, 'linewidth', 3)
grid on
ylim([-20 100])
xlim([min(paramVals) max(paramVals)])
title(upper(mechi))
    
end

end
```
