## Supplementary data file 1 for "Mechanisms of antibiotic action shape the fitness landscapes of resistance mutations": generateOptimizationHistogram.m

```
% Generate a color-coded histogram of kD values from optimization data

% Load in data file
rawdata = readtable('parOptimization_CumResults.csv');

% Grab kF, kR, and fval columns
kF = rawdata.o_binding;
kR = rawdata.o_unbinding;
fval = rawdata.o_funval;

% Filter out NaN's and the really bad fits from all datasets
kF = kF(~isnan(kF));
kR = kR(~isnan(kR));
fval = fval(~isnan(fval));
cutoff = 0.2; % Exclude any fits with fvals above the cutoff
kF = kF(fval < cutoff);
kR = kR(fval < cutoff);
fval = fval(fval < cutoff);

% Calculate kD and its log
kD = kR ./ kF;
logkD = log10(kD);

% Generate the histogram
threshold = 0.155;
[H,colorset,binmeans] = parHistogram9P([logkD fval], 0.1, threshold);

% Output the log-kD values as a csv file
csvwrite('logkDVals.csv', logkD);
```
