## Supplementary data file 1 for "Mechanisms of antibiotic action shape the fitness landscapes of resistance mutations": getKDVals9P.m

```
% Generate kF and kR values based on desired fold changes in kD

% Finds kF and kR such that
% A * kD = (sqrt(A) * kR) / (1/sqrt(A) * kF)

% Colin Hemez
% 2019

function [kinetics, kD] = getKDVals9P(kF, kR, foldvals)

% Take the square root of fold change values
sq = sqrt(foldvals);

% Multiply kR and divide kF with sqrt(A)
kRvals = sq .* kR;
kFvals = (1./sq) * kF;

% Package outputs
kinetics = [kFvals(:) kRvals(:)];
kD = kRvals(:) ./ kFvals(:);

end
```
