## Supplementary data file 1 for "Mechanisms of antibiotic action shape the fitness landscapes of resistance mutations": getMutant9P.m

```
% Get parameters for simulated mutants

% Colin Hemez
% 2019

function Kmut = getMutantR9P(K, mutation)

% Replicate the wild-type kinetic parameters vector
Kmut = K;

% Determine whether mutation is custom or preprogrammed
if ischar(mutation)
    if strcmp(mutation, 'ciproR1')
        target = 5;
        fold = 1/16;
        cost = 0.25;
        
        % Implement a mutation in one of the parameters
        Kmut(target) = Kmut(target) * fold;
        
        % Implement a fitness cost
        Kmut(2) = Kmut(2) * (1 - cost);
    else
        error('MUTATION string is not recognized')
    end
        
else
    if length(mutation) == 3
        target = mutation(1);
        fold   = mutation(2);
        cost   = mutation(3);
        
        % Implement a mutation in one of the parameters
        Kmut(target) = Kmut(target) * fold;
        
        % Implement a fitness cost
        Kmut(2) = Kmut(2) * (1 - cost);
    else
        error('MUTATION vector is incorrect length')
    end
end    

end
```
