## Supplementary data file 1 for "Mechanisms of antibiotic action shape the fitness landscapes of resistance mutations": getMutantC9P.m

```
% Get parameters for simulated resistant AND compensated mutants

% Colin Hemez
% 2019

function [Kcom,Kres,mutation] = getMutantC9P(K,comp_mutation,res_mutation)

% Display initial information
disp('getMutantC9P()...')

% Generate the resistant mutant
[Kres, M] = getMutantR9P(K, res_mutation);

if ischar(res_mutation)
    Rtype = res_mutation;
else
    Rtype = 'custom input';
end

% Replicate the resistant mutant's parameter set
Kcom = Kres;

% Determine whether mutation is custom or preprogrammed
if ischar(comp_mutation)
    if strcmp(comp_mutation, 'ciproC1')
        efficiency = 0.8;
        
        % Implement recovery in fitness cost
        Kcom(2) = Kres(2) + K(2) * efficiency * (1 - (Kres(2) / K(2)));
        
        Ctype = comp_mutation;
    else
        error('MUTATION string is not recognized')
    end
        
else
    if length(comp_mutation) == 1
        efficiency = comp_mutation;
        
        % Implement recovery in fitness cost
        Kcom(2) = Kres(2) + K(2) * efficiency * (1 - (Kres(2) / K(2)));
        
        Ctype = 'custom input';
    else
        error('MUTATION vector is incorrect length')
    end
end

% Package mutation vector
mutation = [M efficiency];

% Display more information
disp(' ')
disp(['RESISTANCE MUTATION:    ', Rtype])
disp(['COMPENSATORY MUTATION:  ', Ctype])

disp(' ')
disp(['Target parameter index..................',num2str(mutation(1))])
disp(['Fold change in parameter value..........',num2str(mutation(2))])
disp(['Cost of resistance mutation.............',num2str(mutation(3))])
disp(['Efficiency of compensatory mutation.....',num2str(mutation(4))])

disp(' ')
disp('Mutant parameters loaded')
disp(' ')

end
```
