## Supplementary data file 1 for "Mechanisms of antibiotic action shape the fitness landscapes of resistance mutations": getMutantR9P.m

```
% Get parameters for simulated resistant mutants

% Colin Hemez
% 2019

function [Kres, res_mutation] = getMutantR9P(K, mutation)

% Replicate the wild-type kinetic parameters vector
Kres = K;

% Determine whether mutation is custom or preprogrammed
if ischar(mutation)
    if strcmp(mutation, 'ciproR1')
        target = 5;
        fold = 1/16;
        cost = 0.25;
        
        % Package mutation vector
        res_mutation = [target fold cost];
        
        % Implement a mutation in one of the parameters
        Kres(target) = Kres(target) * fold;
        
        % Implement a fitness cost
        Kres(2) = Kres(2) * (1 - cost);
    elseif strcmp(mutation, '12MIC25C')
        % Preprogrammed 12-fold increase in MIC with 25% fitness cost
        % Mutation occurs by equal fold change in kF and kR
        cost = 0.25;
        [Q,Kres] = findKdFactor9P(K, 12, cost);
        
        % Package mutation vector
        res_mutation = [5 Q cost];
    else
        error('MUTATION string is not recognized')
    end
        
else
    if length(mutation) == 3
        target = mutation(1);
        fold   = mutation(2);
        cost   = mutation(3);
        
        % Package mutation vector
        res_mutation = [target fold cost];
        
        % Implement a mutation in one of the parameters
        Kres(target) = Kres(target) * fold;
        
        % Implement a fitness cost
        Kres(2) = Kres(2) * (1 - cost);
    else
        error('MUTATION vector is incorrect length')
    end
end    

end
```
