## Supplementary data file 1 for "Mechanisms of antibiotic action shape the fitness landscapes of resistance mutations": getMutParams9P.m

```
% Retrieve parameter sets necessary to generate mutation maps

% Colin Hemez
% 2019

function allParams = getMutParams9P(OG,DR,SI,mode)

% Retrieve kinetic and drug simulation parameters
[kinParams, simParams, mutParams] = getParams9P(OG,DR,SI,1,mode);

% Retrieve mutant parameters
disp('.............................................................')
disp(' ')
[kinCom,kinRes,resParams] = getMutantC9P(kinParams,'ciproC1','12MIC25C');

% Display message
disp('.............................................................')
disp(' ')
disp('getMutParams9P()...')
disp('Calculating MICs (in [g/l]) and mutation selection balance...')

% Calculate MICs
wtMIC  = findMIC9P(kinParams,1e-12,'curve');
resMIC = findMIC9P(kinRes,1e-12,'curve');
comMIC = findMIC9P(kinCom,1e-12,'curve');

% Convert MICs to g/l
wtMIC  = wtMIC * simParams(3);
resMIC = resMIC * simParams(3);
comMIC = comMIC * simParams(3);

% Calculate the mutation-selection balance for the resistant mutant
balance = mutSelBal9P(kinParams,simParams,mutParams,resParams);

% Display MSB and MIC values
disp(' ')
disp(['Mutation selection balance.......',num2str(balance),' [cells]'])
disp(['MIC (wild-type)..................',num2str(wtMIC),' [g/l]'])
disp(['MIC (resistant)..................',num2str(resMIC),' [g/l]'])
disp(['MIC (resistant-compensated)......',num2str(comMIC),' [g/l]'])
disp(' ')

% Generate the struct with all parameters
allParams.Kwt        = kinParams;
allParams.Kres       = kinRes;
allParams.Kcom       = kinCom;
allParams.S          = simParams;
allParams.M          = mutParams;
allParams.R          = resParams;
allParams.MICwt      = wtMIC;
allParams.MICres     = resMIC;
allParams.MICcom     = comMIC;
allParams.MSB        = balance;
allParams.resMutRate = mutParams(1);

% Display message once all calculations are finished
disp('All parameters loaded')
disp(' ')
disp(' ')

end
```
