## Supplementary data file 1 for "Mechanisms of antibiotic action shape the fitness landscapes of resistance mutations": getParams9P.m

```
% Get parameters for simulations

% Colin Hemez
% 2019

function [paramsK, paramsS, paramsM] = getParams9P(OG, DR, SI, varargin)

disp(' ')
disp('getParams9P()...')

% Decide whether to show or suppress dialogue and to declare a drug mode
if isempty(varargin)
    dialog = 1;
elseif length(varargin) == 1
    if ischar(varargin{1})
        mode = varargin{1};
        dialog = 1;
    else
        dialog = varargin{1};
    end
elseif length(varargin) == 2
    if ischar(varargin{1})
        mode = varargin{1};
        dialog = varargin{2};
    elseif ischar(varargin{2})
        mode = varargin{2};
        dialog = varargin{1};
    end
end

% Combine first two input strings
OGDR = [OG DR];
    
if strcmp(OGDR, 'ecolicip2')
        
    %%%%%%%%%%%%%%%%%%%%%%%%%%%%%%%%%%%%%%%%%%%%%%%%
    % E. coli, Ciprofloxacin FOR PARAMETER FITTING %
    %%%%%%%%%%%%%%%%%%%%%%%%%%%%%%%%%%%%%%%%%%%%%%%%
    
    if dialog
        disp(' ')
        disp('ORGANISM:   Escherichia coli')
        disp('ANTIBIOTIC: Ciprofloxacin, FOR PARAMETER FITTING')
    end
    
    % Organism-specific parameters
    n      = 183;
    g0     = 1.4617711221e-4;
    d0     = 1.5e-6; % Mid-life death rate, Wang et al. Curr Biol 2010
    dn     = 0.00209182396;
    kF     = 2680;
    kR     = 3e-4;
    alphaG = 1;
    alphaD = 500;
    betaG  = 0;
    betaD  = 1;
    gammaG = 92;
    gammaD = 156;
    
    % Drug-specific parameters for simulation
    MM = 331.346;
    
    % Mutation rate parameters
    mu_r = 2e-7;
    mu_c = 10 * mu_r;
    
    % Simulation conditions
    if strcmp(SI,'tube')
        
        if dialog
            disp('SIMULATION: Test Tube')
        end
        
        B0 = 6.88e9;
        Vtot = 1e-4;
    elseif strcmp(SI,'infect9')
        
        if dialog
            disp('SIMULATION: Infection, 1e9')
        end
        
        B0 = 1e9;
        Vtot = 1e-4;
    else
        
        disp(' ')
        warning(['SI does not match known inputs. Using default ',...
                 'parameters for a test tube simulation instead.'])
        
        B0 = 6.88e9;
        Vtot = 1e-4;
    end
    
elseif strcmp(OGDR, 'ecolicipP1')
        
    %%%%%%%%%%%%%%%%%%%%%%%%%%%%%%%%%%%%%%%%%%%%%%%%
    % E. coli, Ciprofloxacin FOR PARAMETER FITTING %
    %%%%%%%%%%%%%%%%%%%%%%%%%%%%%%%%%%%%%%%%%%%%%%%%
    
    if dialog
        disp(' ')
        disp('ORGANISM:   Escherichia coli')
        disp('ANTIBIOTIC: Ciprofloxacin, PARAMETER FIT 190404')
    end
    
    % Organism-specific parameters
    n      = 183;
    g0     = 1.4617711221e-4;
    d0     = 1.5e-6; % Mid-life death rate, Wang et al. Curr Biol 2010
    dn     = 0.00209182396;
    kF     = 5.238062350476559e+03;
    kR     = 3.180468079577469e-04;
    alphaG = 16.793136035734705;
    alphaD = 7.280324617722957;
    betaG  = 0;
    betaD  = 1;
    gammaG = 24.835562592641857;
    gammaD = 3.637135304720576e+02;
    
    % Drug-specific parameters for simulation
    MM = 331.346;
    
    % Mutation rate parameters
    mu_r = 2e-7;
    mu_c = 10 * mu_r;
    
    % Simulation conditions
    if strcmp(SI,'tube')
        
        if dialog
            disp('SIMULATION: Test Tube')
        end
        
        B0 = 6.88e9;
        Vtot = 1e-4;
    elseif strcmp(SI,'infect9')
        
        if dialog
            disp('SIMULATION: Infection, 1e9')
        end
        
        B0 = 1e9;
        Vtot = 1e-4;
    else
        
        disp(' ')
        warning(['SI does not match known inputs. Using default ',...
                 'parameters for a test tube simulation instead.'])
        
        B0 = 6.88e9;
        Vtot = 1e-4;
    end
    
else
    
    %%%%%%%%%%%%%%%%%%%%%%%
    % Inputs do not match %
    %%%%%%%%%%%%%%%%%%%%%%%
    
    disp(' ')
    warning(['OG and/or DR do not match a known input ',...
             'combination. Using default parameters for ',...
             'E. coli and ciprofloxacin instead.'])
    
    % Organism-specific parameters
    n      = 183;
    g0     = 1.4617711221e-4;
    d0     = 1.5e-6; % Mid-life death rate, Wang et al. Curr Biol 2010
    dn     = 0.00209182396;
    kF     = 2680;
    kR     = 3e-4;
    alphaG = 1;
    alphaD = 500;
    betaG  = 0;
    betaD  = 1;
    gammaG = 92;
    gammaD = 156;
    
    % Drug-specific parameters for simulation
    MM = 331.346;
    
    % Mutation rate parameters
    mu_r = 2e-7;
    mu_c = 10 * mu_r;
    
    % Simulation conditions
    if strcmp(SI,'tube')
        
        if dialog
            disp(' ')
            disp('SIMULATION: Test Tube')
        end
        
        B0 = 6.88e9;
        Vtot = 1e-4;
    elseif strcmp(SI,'infect9')
        
        if dialog
            disp('SIMULATION: Infection, 1e9')
        end
        
        B0 = 1e9;
        Vtot = 1e-4;
    else
        
        disp(' ')
        warning(['SI does not match known inputs. Using default ',...
                 'parameters for a test tube simulation instead.'])
        
        B0 = 6.88e9;
        Vtot = 1e-4;
    end
end

% Drug-specific extremes
if exist('mode', 'var')
    if strcmp(mode, 'nstat')
        alphaG = 500;
        alphaD = 1;
        betaG  = 0;
        betaD  = 0;
        gammaG = 35;
        gammaD = 150;
    elseif strcmp(mode, 'ncide')
        alphaG = 1;
        alphaD = 500;
        betaG  = 1;
        betaD  = 1;
        gammaG = 35;
        gammaD = 150;
    elseif strcmp(mode, 'cstat')
        alphaG = 1;
        alphaD = 1;
        betaG  = 0;
        betaD  = 0;
        gammaG = 35;
        gammaD = 150;
    elseif strcmp(mode, 'ccide')
        alphaG = 1;
        alphaD = 1;
        betaG  = 1;
        betaD  = 1;
        gammaG = 35;
        gammaD = 150;
    elseif strcmp(mode, 'general')
        alphaG = 1;
        alphaD = 500;
        betaG  = 0;
        betaD  = 1;
        gammaG = 35;
        gammaD = 150;
    elseif strcmp(mode, 'curve')
    elseif strcmp(mode, 'cipro')
    else
        error('MODE not recognized')
    end
end

% Display parameters
if dialog
    disp(' ')
    disp('KINETIC PARAMETERS:')
    disp(['Number of targets.............',num2str(n),'  [#/cell]'])
    disp(['Growth rate...................',num2str(g0),'  [1/sec]'])
    disp(['Death rate (w/o drug).........',num2str(d0),'  [1/sec]'])
    disp(['Death rate (w/ drug)..........',num2str(dn),'  [1/sec]'])
    disp(['Forward binding rate..........',num2str(kF),'  [1/M/sec]'])
    disp(['Reverse unbinding rate........',num2str(kR),'  [1/sec]'])
    disp(['Growth curve sharpness........',num2str(alphaG),'  [unitless]'])
    disp(['Death curve sharpness.........',num2str(alphaD),'  [unitless]'])
    disp(['Bacteriostatic potency........',num2str(betaG),'  [unitless]'])
    disp(['Bactericidal potency..........',num2str(betaD),'  [unitless]'])
    disp(['Growth inflection point.......',num2str(gammaG),'  [#]'])
    disp(['Death inflection point........',num2str(gammaD),'  [#]'])
    disp(' ')
    disp('SIMULATION PARAMETERS:')
    disp(['Initial population size.......',num2str(B0),' [#]'])
    disp(['Simulation volume.............',num2str(Vtot),' [l]'])
    disp(['Molar mass of drug............',num2str(MM),' [g/mol]'])
    disp(' ')
    disp('MUTATION RATE PARAMETERS:')
    disp(['Mutation rate (resistance)....',num2str(mu_r),' [1/division]'])
    disp(['Mutation rate (compensation)..',num2str(mu_c),' [1/division]'])
    disp(' ')
end

paramsK = [n g0 d0 dn kF kR alphaG alphaD betaG betaD gammaG gammaD];
paramsS = [B0 Vtot MM];
paramsM = [mu_r mu_c];

disp('Parameters loaded')
disp(' ')

end
```
