## Supplementary data file 1 for "Mechanisms of antibiotic action shape the fitness landscapes of resistance mutations": GiDiCurvePlotter.m

```
% Plotting the effect of alpha and gamma on growth and death curves
% Data generation for supplementary figure

% Colin Hemez
% 2019

[K,S] = getParams9P('ecoli','cipP1','tube');

Nvals = (0:1:K(1))';

Ki = K;

aG = [1 5 12 20 50 500];
cG = [-100 0 40 100 160 180];
aD = [1 5 12 20 50 500];
cD = [5 20 60 120 170 300];

aGcurves = zeros((K(1)+1), length(aG));
cGcurves = zeros((K(1)+1), length(cG));
aDcurves = zeros((K(1)+1), length(aD));
cDcurves = zeros((K(1)+1), length(cD));

%%%%%%%%%%%
% ALPHA-G %
%%%%%%%%%%%
figure
subplot(2,2,1)
Ki(11) = 60;
for i = 1:length(aG)
    Ki(7) = aG(i);
    [A,B,G,D] = systemMatrices9P(Ki,0,0);
    plot(Nvals,G,'k','linewidth',2)
    aGcurves(:,i) = G;
    hold on
end
grid on
xlim([0 K(1)])
title(['Varying aG (cG = ',num2str(Ki(11)),')'])
Ki = K;

%%%%%%%%%%%
% GAMMA-G %
%%%%%%%%%%%
subplot(2,2,2)
Ki(7) = 8;
for i = 1:length(cG)
    Ki(11) = cG(i);
    [A,B,G,D] = systemMatrices9P(Ki,0,0);
    plot(Nvals,G,'k','linewidth',2)
    cGcurves(:,i) = G;
    hold on
end
grid on
xlim([0 K(1)])
title(['Varying cG (aG = ',num2str(Ki(7)),')'])
Ki = K;

%%%%%%%%%%%
% ALPHA-D %
%%%%%%%%%%%
subplot(2,2,3)
Ki(12) = 150;
for i = 1:length(aD)
    Ki(8) = aD(i);
    [A,B,G,D] = systemMatrices9P(Ki,0,0);
    plot(Nvals,D,'k','linewidth',2)
    aDcurves(:,i) = D;
    hold on
end
grid on
xlim([0 K(1)])
title(['Varying aD (cD = ',num2str(Ki(12)),')'])
Ki = K;

%%%%%%%%%%%
% GAMMA-D %
%%%%%%%%%%%
subplot(2,2,4)
Ki(8) = 40;
for i = 1:length(cD)
    Ki(12) = cD(i);
    [A,B,G,D] = systemMatrices9P(Ki,0,0);
    plot(Nvals,D,'k','linewidth',2)
    cDcurves(:,i) = D;
    hold on
end
grid on
xlim([0 K(1)])
title(['Varying cD (aD = ',num2str(Ki(8)),')'])
Ki = K;

writematrix(aGcurves.*3600,'aGcurves.csv')
writematrix(cGcurves.*3600,'cGcurves.csv')
writematrix(aDcurves.*3600,'aDcurves.csv')
writematrix(cDcurves.*3600,'cDcurves.csv')

% Define color maps
R = [224 25 76];
B = [56 113 237];
G = [247 193 14];
Y = [61 174 12];

P = length(aG);
R1 = linspace(B(1), R(1), P);
G1 = linspace(B(2), R(2), P);
B1 = linspace(B(3), R(3), P);
RGB1 = [R1' G1' B1'];

R2 = linspace(G(1), Y(1), P);
G2 = linspace(G(2), Y(2), P);
B2 = linspace(G(3), Y(3), P);
RGB2 = [R2' G2' B2'];
```
