## Supplementary data file 1 for "Mechanisms of antibiotic action shape the fitness landscapes of resistance mutations": isoclimePlotter.m

```
% Plot fitness isoclimes

mech = {'cipro','nstat','cstat','ncide','ccide'};
param = {'kF','kR'};

foldvals = linspace(1,100,100);

figure
subp = 1;

for i = 1:length(param)
    for j = 1:length(mech)
        
        isoi = load(['fitnessIsoclime_',param{i},'_',mech{j},'.csv']);
        
        subplot(length(param),length(mech),subp)
        plot(foldvals,isoi,'-k','linewidth',2)
        grid on
        ylim([-20 100])
        
        subp = subp + 1;
    end
end
```
