## Supplementary data file 1 for "Mechanisms of antibiotic action shape the fitness landscapes of resistance mutations": isoclimesKFKR.m

```
% Generate fitness isoclimes for kF anf kR parameter fold changes
% 190527

clear;
close all;

resolution = 100;
foldMICs = [1 10 20 30 40 50 60 70 80 90 100];
foldParamVals = linspace(1, 100, resolution)';

csvwrite('fitnessIsoclimeMICfolds.csv', foldMICs)
csvwrite('fitnessIsoclimeFoldParams.csv', foldParamVals);

isoFamKF = fitnessIsoclimeFamily9P(foldMICs, 'kF', foldParamVals);
isoFamKR = fitnessIsoclimeFamily9P(foldMICs, 'kR', foldParamVals);
```
