## Supplementary data file 1 for "Mechanisms of antibiotic action shape the fitness landscapes of resistance mutations": killCurvesMech1.m

```
% Simulated kill curves for four extreme modes of antibiotic action

clear; close all;

mech = {'cipro','nstat','cstat','ncide','ccide'};

res = 200;
factor = 4;
Binit = 1;
trange = 6;
T = linspace(0, trange*3600, res);

B = zeros(res, length(mech));

for i = 1:length(mech)
    
    X = getMutParams9P('ecoli','cipP1','infect9',mech{i});
    
    X.S(1) = 1e8;
    
    C = factor * X.MICwt;
    
    Bi = ABkinetic9P(X.Kwt,X.S,Binit,C,T,'curve','exact');
    
    Bs = sum(Bi);
    Bs = Bs(:);
    
    B(:,i) = Bs;

end

figure
plot(T./3600, B, 'linewidth', 3);
grid on

figure
semilogy(T./3600, B, 'linewidth', 3);
grid on

csvwrite('mechKillCurves_timeVals.csv',T./3600)
csvwrite('mechKillCurves_popVals.csv',B./(X.S(1)))
```
