## Supplementary data file 1 for "Mechanisms of antibiotic action shape the fitness landscapes of resistance mutations": kineticCurvesConc1.m

```
% Plot ABkinetic curves at different concentrations relative to MIC

clear;

X = getMutParams9P('ecoli','cipP1','infect9','cipro');

% Set the initial cell population
B0 = 1e8;
X.S(1) = B0;

% Set the time range and resolution
Thours = 8;
res = 100;
T = linspace(0, Thours, res)' .* 3600;
csvwrite('kineticCurves1_timeVals.csv', T ./ 3600)

% Define the drug concentrations
MICfolds = [0 0.25 0.5 1 2 4 8 16];
csvwrite('kineticCurves1_MICfolds.csv', MICfolds)
Cvals = MICfolds .* X.MICwt;

% Set aside some memory
popVals = zeros(res, length(MICfolds));

% Run the ABkinetic simulations
Binit = 1;
for i = 1:length(Cvals)
    C = Cvals(i);
    Bi = ABkinetic9P(X.Kwt, X.S, Binit, C, T, 'curve', 'exact');
    Bisum = sum(Bi);
    Bisum = Bisum(:);
    popVals(:,i) = Bisum;
end

csvwrite('kineticCurves1_PopVals.csv', popVals)

% Plot
figure
semilogy(T ./ 3600, popVals, 'linewidth', 3)
grid on
```
