## Supplementary data file 1 for "Mechanisms of antibiotic action shape the fitness landscapes of resistance mutations": MICfoldKFKR1.m

```
% Fold-increase in MIC given fold-changes in kF and kR

clear; close all;

tol = 1e-12;
foldmax = 100;
res = 40;
foldvals = linspace(1, foldmax, res);

mech = {'cipro','nstat','cstat','ncide','ccide'};

% Allocate memory
mkf = zeros(res, length(mech));
mkr = mkf;

% Run the simulations
for i = 1:length(mech)
    
    X = getMutParams9P('ecoli','cipP1','infect9',mech{i});
    
    MWT = X.MICwt / X.S(3);
    currKf = X.Kwt;
    currKr = currKf;
    
    for j = 1:res
        currKf(5) = X.Kwt(5) / foldvals(j);
        currKr(6) = X.Kwt(6) * foldvals(j);
        
        MICKF = findMIC9P(currKf, tol, 'curve');
        MICKR = findMIC9P(currKr, tol, 'curve');
        
        mkf(j,i) = MICKF / MWT;
        mkr(j,i) = MICKR / MWT;
        
    end
    
end

figure
subplot(1,2,1)
plot(foldvals, mkf)
grid on

subplot(1,2,2)
plot(foldvals, mkr)
grid on

csvwrite('MICfold_foldVals.csv',foldvals');
csvwrite('MICfold_KFvals.csv', mkf);
csvwrite('MICfold_KRvals.csv', mkr);
```
