## Supplementary data file 1 for "Mechanisms of antibiotic action shape the fitness landscapes of resistance mutations": MICrate1.m

```
% Calculate MIC as a function of growth rate

mech = {'cipro','nstat','cstat','ncide','ccide'};

Nvals = [1 3] .* 183;

res = 40;
MICvals = zeros(res,length(mech),length(Nvals));

for i = 1:length(mech)
    X = getMutParams9P('ecoli','cipP1','infect9',mech{i});
    K = X.Kwt;
    currK = K;
    
    delG = K(1) - K(11);
    gamVals = Nvals - delG;

    rates = linspace((K(3) * 1.01), K(2), res);

    MICs = zeros(res, length(Nvals));
    
    for j = 1:length(Nvals)
        currK(1) = Nvals(j);
        currK(11) = gamVals(j);
        currK(2) = rates(end);
        MICw = findMIC9P(currK,1e-12,'curve');
        
        for k = 1:res
            currK(2) = rates(k);
            
            MICijk = findMIC9P(currK,1e-12,'curve');
            
            MICs(k,j) = MICijk;
            
        end
        
        MICvals(:,i,j) = MICs(:,j) ./ MICw;
        
    end
    
end

figure
for i = 1:length(Nvals)
    subplot(1,length(Nvals),i)
    plot(rates, MICvals(:,:,i),'linewidth',2)
    grid on
    legend(mech,'location','southeast')
end
```
