## Supplementary data file 1 for "Mechanisms of antibiotic action shape the fitness landscapes of resistance mutations": MICSurvTox1.m

```
% MIC finder for survival threshold on growth and toxicity on death

% clear;
% close all;

X = getMutParams9P('ecoli','cipP1','infect9','cipro');

K = X.Kwt;

res = 100;
delG = K(1) - K(11);

Nvals = round(linspace(61,900,res));
gammaGvals = Nvals - delG;

Kmat = zeros(res, length(K));
for i = 1:res
    Kmat(i,:) = K;
    Kmat(i,1) = Nvals(i);
    Kmat(i,11) = gammaGvals(i);
end

MICs = zeros(res,1);

for i = 1:res
    disp(['Nval: ', num2str(Nvals(i))])
    MICi = findMIC9P(Kmat(i,:), 1e-12, 'curve');
    MICs(i) = MICi;
end

figure
plot(Nvals, MICs.*X.S(3), 'linewidth', 4)
hold on
plot(K(1), X.MICwt, 'ok', 'markerfacecolor', [0 0 0], 'markersize', 10)
grid on
xlabel('Target Number')
ylabel('MIC')
title('Varying target number; growth on a survival threshold and death on a toxicity threshold')
```
