## Supplementary data file 1 for "Mechanisms of antibiotic action shape the fitness landscapes of resistance mutations": MICtargNum1.m

```
% MIC as a function of target number and fitness cost

clear
close all

X = getMutParams9P('ecoli','cipP1','infect9','cipro');
Ki = X.Kwt;
delG = Ki(1) - Ki(11);

res1 = 50;
res2 = 50;
FH = linspace(0, 90, res1);
TN = round(linspace(61, 732, res2));

MICs = zeros(res1, res2);

for i = 1:res1
    FHi = FH(i);
    FHi = FHi / 100;
    Ki(2) = Ki(2) * (1 - FHi);
    
    disp(['FH: ',num2str(FHi * 100)])
    
    for j = 1:res2
        TNj = TN(j);
        Ki(1) = TNj;
        Ki(11) = TNj - delG;
        
        MICij = findMIC9P(Ki, 1e-12, 'curve');
        
        MICs(i,j) = MICij;
        
    end
end

normMIC = MICs ./ X.MICwt;
```
