## Supplementary data file 1 for "Mechanisms of antibiotic action shape the fitness landscapes of resistance mutations": MSWFamily9P.m

```
% Generate a family of MSW curves for all drug mechanisms

% Colin Hemez
% 2019

function [concFactors, netRateDiffs] = MSWFamily9P(resolution, expTime)

% Define the drug mechanisms
mech = {'cipro', 'nstat', 'cstat', 'ncide', 'ccide'};

% Set aside memory
concFactors = zeros(resolution, length(mech));
netRateDiffs= concFactors;

% Determine the MSWs for each mechanism
for i = 1:length(mech)
    
    % Retrieve parameters
    mechi = mech{i};
    Xi = getMutParams9P('ecoli','cipP1','infect9',mechi);
    
    % Calculate the MSW and store
    [Fi,Di] = pdCurveMSW9P(Xi.Kwt, Xi.Kres, Xi.S, resolution, expTime);
    concFactors(:,i) = Fi;
    netRateDiffs(:,i) = Di;
end

% Plot the results
figure
for i = 1:length(mech)
    semilogx(concFactors(:,i), netRateDiffs(:,i)./Xi.Kres(2),...
             'linewidth', 3)
    hold on
end
legend(upper(mech),'location','northwest')
grid on

end
```
