## Supplementary data file 1 for "Mechanisms of antibiotic action shape the fitness landscapes of resistance mutations": mutationMap9P.m

```
% Generate a heat map of probabilities for mutation emergence

% Colin Hemez
% 2019

function [P,result] = mutationMap9P(kinParams,...
                                    simParams,...
                                    N0,...
                                    mu,...
                                    C0,...
                                    Xtarget,...
                                    Xvals,...
                                    Ytarget,...
                                    Yvals,...
                                    varargin)

% Package initial population size into simulation parameters vector
simParams(1) = N0;

% Package all parameters into a single vector
mapParams = [kinParams simParams mu C0];

% Determine which parameter to vary along Y axis of mutation map
if ischar(Ytarget)
    if strcmp(Ytarget, 'kF')
        Yt = 5;
    elseif strcmp(Ytarget, 'kR')
        Yt = 6;
    elseif strcmp(Ytarget, 'kD')
        Yt = [5 6];
    elseif strcmp(Ytarget, 'N0') % Mutation selection balance
        Yt = 13;
    elseif strcmp(Ytarget, 'mu')
        Yt = 16;
    elseif strcmp(Ytarget, 'C0')
        Yt = 17;
    end
else
    Yt = Ytarget;
end

% Convert inputs to column vectors
Xvals = Xvals(:);
% Yvals = Yvals(:);

% Calculate probabilities for all combinations of X and Y parameters
Pmat = zeros(length(Yvals), length(Xvals));

for i = 1:length(Yvals)
    
    for j = 1:length(Yt)
        mapParams(Yt(j)) = Yvals(i,j);
    end
    
    % Display some information
    disp(' ')
    disp(' ')
    disp('%%%%%%')
    disp('%%%%%%%%%')
    disp(['CALCULATING PROBABILITIES FOR Y-AXIS VALUE '...
          num2str(i),...
          ' OF ',...
          num2str(length(Yvals))])
    disp('%%%%%%%%%')
    disp('%%%%%%')
    
    % Calculate expected number of mutants and associated probability
    Pi = probabilityCurve9P(mapParams(1:12),...
                            mapParams(13:15),...
                            mapParams(13),...
                            mapParams(16),...
                            mapParams(17),...
                            Xtarget,...
                            Xvals,...
                            varargin);
    
    Pmat(i,:) = Pi;
   
end

% Package outputs into struct
P = Pmat;
result.P = Pmat;
result.Xtarget = Xtarget;
result.Xvalues = Xvals;
result.Ytarget = Ytarget;
result.Yvalues = Yvals;

% Generate color scheme and probability cutoffs
C = [35  35  35
     13  101 168
     224 13  104
     247 193 14]; % Color map
 
cutoff = [0.01 0.1];

% Make color map based on probability values
dim = size(Pmat);
Pmap = zeros(dim(1), dim(2), 3);

for i = 1:dim(1)
    for j = 1:dim(2)
        Pij = Pmat(i,j);
        if Pij < cutoff(1)
            ind = (0 - Pij) / (0 - cutoff(1));
            Crange = C(2,:) - C(1,:);
            A = C(1,:) + Crange .* ind;
            Pmap(i,j,:) = A;
        elseif Pij < cutoff(2)
            ind = (cutoff(1) - Pij) / (cutoff(1) - cutoff(2));
            Crange = C(3,:) - C(2,:);
            A = C(2,:) + Crange .* ind;
            Pmap(i,j,:) = A;
        else
            ind = (cutoff(2) - Pij) / (cutoff(2) - 1);
            Crange = C(4,:) - C(3,:);
            A = C(3,:) + Crange .* ind;
            Pmap(i,j,:) = A;
        end
    end
end

disp(' ')
disp(' ')

% Show the image
figure
Pmap = Pmap ./ 255;
image(flip(Pmap));
axis off

end
```
