## Supplementary data file 1 for "Mechanisms of antibiotic action shape the fitness landscapes of resistance mutations": mutSelBal9P.m

```
% Determine the mutation selection balance for a resistant mutant

% Colin Hemez
% 2019

function E = mutSelBal9P(kinParams,simParams,mutParams,resParams)

% Unpack parameters
D  = kinParams(3); % Death rate in the absence of drug [1/sec]
N0 = simParams(1); % Initial size of infecting population [cells]
mu = mutParams(1); % Mutation rate for resistance emergence [1/division]
cr = resParams(3); % Cost of the resistance mutation [unitless]

% Convert mutation rate to 1/sec
mu = mu * kinParams(2);

% Compute the mutation selection balance
E = (N0 * mu) / (cr * D);

end
```
