## Supplementary data file 1 for "Mechanisms of antibiotic action shape the fitness landscapes of resistance mutations": mutSelWindowVals.m

```
% Generate concentration values for mutant selection windows

% Colin Hemez
% 2019

clear;
close all;

mech = {'cipro','nstat','cstat','ncide','ccide'};

Cvals = zeros(3, length(mech));

for i = 1:length(mech)
    mechi = mech{i};
    
    Xi = getMutParams9P('ecoli','cipP1','infect9',mechi);
    
    B = Xi.MICres ./ Xi.MICwt;
    
    % Find the concentration at which resistant outcompetes wild-type
    A = pdCurveIntersect9P(Xi.Kwt,Xi.Kres,Xi.S,1e-12,96);
    A = A ./ Xi.MICwt;
    
    % Find the concentration at which compensation probability < 0.01
    C = compensationProb9P(Xi.Kres,Xi.S,Xi.M,Xi.MSB,0.01,1e-4,'curve','exact');
    C = C ./ Xi.MICwt;
    
    Cvals(:,i) = [A ; B ; C];
    
end

csvwrite('MSW.csv', Cvals)
csvwrite('MSWlog.csv', log10(Cvals))
```
