## Supplementary data file 1 for "Mechanisms of antibiotic action shape the fitness landscapes of resistance mutations": netRateCalc9P.m

```
% Net growth rate calculator
% Colin Hemez
% 2018

function change = netRateCalc9P(kinParams,...
                                simParams,...
                                Binit,...
                                C0,...
                                timeVals,...
                                varargin)

% Run kinetic simulation
step = max(timeVals) - min(timeVals);
T = [min(timeVals) max(timeVals)];

netrate = zeros(length(C0), 1);

for i = 1:length(C0)
    C = C0(i);
    Bi = ABkinetic9P(kinParams, simParams, Binit, C, T, varargin);
    Bsum = sum(Bi);
    
    % Calculate net growth rate
    netrate(i) = log(Bsum(2) / Bsum(1)) / step;
end

change = netrate;

end
```
