## Supplementary data file 1 for "Mechanisms of antibiotic action shape the fitness landscapes of resistance mutations": netRateCalcData.m

```
% Calculate net growth rates from experimental time kill curves

% Colin Hemez
% 2019

function [netrates,conc] = netRateCalcData(timekill)

% Pull out concentration value vector, time vector, and population values
conc = timekill(2:end,1);
time = timekill(1,2:end);
pops = timekill(2:end,2:end);

% Pull out the maximum timepoint value
endpoint = max(time);

% Calculate a net growth rate for each concentration value
netrates = zeros(length(conc), 1);

for i = 1:length(conc)
    startpop = pops(i,1);
    endpop = pops(i,end);
    
    netrates(i) = log(endpop / startpop) / endpoint;
end

end
```
