## Supplementary data file 1 for "Mechanisms of antibiotic action shape the fitness landscapes of resistance mutations": noParOptimize9P.m

```
% Replicate model minimizations NOT in parallel

% Colin Hemez
% 2019

function solutions = noParOptimize9P(problem, iterations)

solutions = zeros(iterations,16);
soli = zeros(1,16);

for i = 1:iterations
    [S1i] = ABkineticMinimize9P(problem);
    close all
    
    % Package data to store
    soli(1:12) = S1i.paramset;
    soli = [soli S1i.fvalue S1i.pearson S1i.exit S1i.output.totaltime];
    solutions(i,:) = soli;
end

end
```
