## Supplementary data file 1 for "Mechanisms of antibiotic action shape the fitness landscapes of resistance mutations": parHistogram9P.m

```
% Generate a histogram of parameter values

% Colin Hemez
% 2019

function [H,colors,binmeans] = parHistogram9P(data, bin, cthreshold)

Hdata = data(:,1); % Values to plot on the histogram
Cdata = data(:,2); % Values for color-coding the histogram bars

H = figure;

% Define the bin limits
mag = floor(log10(bin));
minbin = round((min(Hdata) - bin), abs(mag));
maxbin = round((max(Hdata) + bin), abs(mag));
binlim = minbin:bin:maxbin;
binnum = length(binlim) - 1;
binavg = zeros(1, binnum);

% Calculate average Cdata values for each bin
for i = 1:(binnum)
    binmini = binlim(i);
    binmaxi = binlim(i+1);
    Bi = Cdata((Hdata >= binmini) & (Hdata < binmaxi));
    Mi = mean(Bi);
    binavg(i) = Mi;
end

% Generate the color map for the histogram
RGB = [56  113 237
       224 25  76
       247 193 14];
   
minavg = min(binavg);
maxavg = max(binavg);

binrgb = zeros(binnum, 3);

for i = 1:binnum
    Ci = binavg(i);
    
    if Ci < cthreshold
        P = (Ci - minavg) / (cthreshold - minavg);
        R = RGB(3,1) - (P * (RGB(3,1) - RGB(2,1)));
        G = RGB(3,2) - (P * (RGB(3,2) - RGB(2,2)));
        B = RGB(3,3) - (P * (RGB(3,3) - RGB(2,3)));
    else
        P = (Ci - cthreshold) / (maxavg - cthreshold);
        R = RGB(2,1) - (P * (RGB(2,1) - RGB(1,1)));
        G = RGB(2,2) - (P * (RGB(2,2) - RGB(1,2)));
        B = RGB(2,3) - (P * (RGB(2,3) - RGB(1,3)));
    end
    
    binrgb(i,:) = [R G B] ./ 255;
    
end

% Make the plot
for i = 1:binnum
    binmini = binlim(i);
    binmaxi = binlim(i+1);
    Hi = Hdata((Hdata >= binmini) & (Hdata < binmaxi));
    rgbi = binrgb(i,:);
    if isnan(sum(rgbi))
        rgbi = [0 0 0];
    end
    histogram(Hi, 'binwidth', bin,...
                  'facecolor', rgbi,...
                  'facealpha', 1,...
                  'linewidth', 1.5)
    hold on
end

xlim([minbin maxbin])
grid on
xlabel('Parameter value')
ylabel('Frequency')

colors = binrgb .* 255;

binmeans = binavg';

end
```
