## Supplementary data file 1 for "Mechanisms of antibiotic action shape the fitness landscapes of resistance mutations": parOptimize9P.m

```
% Replicate model minimizations in parallel

% Colin Hemez
% 2019

function [solutions, plotstuff] = parOptimize9P(problem, iterations)

% Unpack the needed parameters
strategy = problem.strategy;
LB       = problem.paramLB;
UB       = problem.paramUB;

% Determine the optimization strategy
if strcmp(strategy, '8param1')
    % 8-parameter minimization: kF, kR, aG, aD, bG, bD, cG, cD
    optIndex = 5:12;
elseif strcmp(strategy, '2param1')
    % 2-parameter minimization: cG, cD
    optIndex = 11:12;
elseif strcmp(strategy, '7param1')
    % 7-parameter minimization: kF, kR, aG, aD, bG, cG, cD
    optIndex = [5 6 7 8 9 11 12];
elseif strcmp(strategy, '6param1')
    % 6-parameter minimization: kF, kR, aG, aD, cG, cD
    optIndex = [5 6 7 8 11 12];
elseif strcmp(strategy, '4param1')
    % 4-parameter minimization: aG, aD, cG, cD
    optIndex = [7 8 11 12];
elseif strcmp(strategy, '2param2')
    % 4-parameter minimization: kF, kR
    optIndex = [5 6];
end

% Make a figure to display kF/kR/kD values as they arrive from workers
plotLB = [LB(5) LB(6) (LB(6)/UB(5))];
plotUB = [UB(5) UB(6) (UB(6)/LB(5))];

close all
figure

K1 = nan(1,iterations);
K2 = K1;

% kF plot
subplot(3,1,1)
loglog([plotLB(1) plotUB(1)],[plotLB(1) plotUB(1)],'--r','linewidth',2)
hold on
P1 = loglog(K1,K2,'ok','markersize',5,'markerfacecolor',[0 0 0]);
xlim([plotLB(1) plotUB(1)])
ylim([plotLB(1) plotUB(1)])
xlabel('Parameter start value')
ylabel('Parameter value after optimization')
grid on

% kR plot
subplot(3,1,2)
loglog([plotLB(2) plotUB(2)],[plotLB(2) plotUB(2)],'--r','linewidth',2)
hold on
P2 = loglog(K1,K2,'ok','markersize',5,'markerfacecolor',[0 0 0]);
xlim([plotLB(2) plotUB(2)])
ylim([plotLB(2) plotUB(2)])
xlabel('Parameter start value')
ylabel('Parameter value after optimization')
grid on

% kD plot
subplot(3,1,3)
loglog([plotLB(3) plotUB(3)],[plotLB(3) plotUB(3)],'--r','linewidth',2)
hold on
P3 = loglog(K1,K2,'ok','markersize',5,'markerfacecolor',[0 0 0]);
xlim([plotLB(3) plotUB(3)])
ylim([plotLB(3) plotUB(3)])
xlabel('Parameter start value')
ylabel('Parameter value after optimization')
grid on

P = [P1 P2 P3];

% Set aside memory for the dataset
dataset = nan(iterations, 29);
filename = 'parData9P.csv';
writematrix(dataset, filename)

% Start a parallel pool
delete(gcp('nocreate'))
parpool;

% Set up a data queue
Q = parallel.pool.DataQueue;
Q.afterEach(@(solution)parOptimizeData9P(solution,...
                                         iterations,...
                                         P,...
                                         filename));

parfor ii = 1:iterations
    % Generate a random set of starting conditions
    Xii = problem;
    randK = randKinParams9P(problem, optIndex);
    K = problem.kinParams;
    K(optIndex) = randK;
    Xii.kinParams = K;
    
    % Run the optimization
    [S1i] = ABkineticMinimizePar9P(Xii);
    
    % Package data to store
    solK = S1i.paramset;
    soli = [K solK S1i.fvalue S1i.pearson S1i.exit S1i.output.totaltime];
    solutions(ii,:) = soli;
    send(Q, [ii soli]);
end

delete(gcp('nocreate'))

solutions = [(1:iterations)' solutions];

plotstuff = parResultProcessor9P(solutions, problem);

end
```
