## Supplementary data file 1 for "Mechanisms of antibiotic action shape the fitness landscapes of resistance mutations": parOptimizeData9P.m

```
% Manage data coming from the data queue of parOptimize9P

% Colin Hemez
% 2019

function parOptimizeData9P(solution,...
                           iterations,...
                           plot,...
                           filename)

% Name the parameters to plot
paramNames = {'k_F','k_R','k_D'};
paramIndex = [5 6];

ii = solution(1);

% Update kF plot data and define number of completed iterates
subplot(3,1,1)
plot(1).XData(ii) = solution(paramIndex(1) + 1);
X = plot(1).XData;
incompleteIterates = sum(isnan(X));
completeIterates = iterations - incompleteIterates;
plot(1).YData(ii) = solution(paramIndex(1) + 13);
title([paramNames{1},...
       ', Number of completed optimizations: ',...
       num2str(completeIterates)])
drawnow('limitrate')

% Update kR plot data
subplot(3,1,2)
plot(2).XData(ii) = solution(paramIndex(2) + 1);
plot(2).YData(ii) = solution(paramIndex(2) + 13);
title([paramNames{2},...
       ', Number of completed optimizations: ',...
       num2str(completeIterates)])
drawnow('limitrate')

% Update kD plot data
subplot(3,1,3)
plot(3).XData(ii) = solution(paramIndex(2)+1)/solution(paramIndex(1)+1);
plot(3).YData(ii) = solution(paramIndex(2)+13)/solution(paramIndex(1)+13);
title([paramNames{3},...
       ', Number of completed optimizations: ',...
       num2str(completeIterates)])
drawnow('limitrate')

% Update data file
data = load(filename);
data(ii,:) = solution;
writematrix(data,filename)
fvals = data(:,26);

% Give client an update on progress
disp(' ')
disp('===============================')
disp('==================================')
disp(['ITERATIONS COMPLETED: ',...
      num2str(completeIterates),...
      ' of ',...
      num2str(iterations)])
disp(['Best fvalue so far:    ', num2str(min(fvals))])
disp(['Average fvalue so far: ', num2str(mean(fvals(~isnan(fvals))))])
disp('==================================')
disp('===============================')
disp(' ')

end
```
