## Supplementary data file 1 for "Mechanisms of antibiotic action shape the fitness landscapes of resistance mutations": parResultProcessor9P.m

```
% Plot the data generated from parallel optimizations

function plotdata = parResultProcessor9P(parsolutions, problem)

% Extract bounds from the problem struct
LB = problem.paramLB;
UB = problem.paramUB;

% Extract results from the solutions matrix
kF    = parsolutions(:,18);
kR    = parsolutions(:,19);
aG    = parsolutions(:,20);
aD    = parsolutions(:,21);
cG    = parsolutions(:,24);
cD    = parsolutions(:,25);
fvals = parsolutions(:,26);
kFs   = parsolutions(:,6);
kRs   = parsolutions(:,7);
aGs   = parsolutions(:,8);
aDs   = parsolutions(:,9);
cGs   = parsolutions(:,12);
cDs   = parsolutions(:,13);

% Remove NaNs from result vectors
kF    = kF(~isnan(kF));
kR    = kR(~isnan(kR));
aG    = aG(~isnan(aG));
aD    = aD(~isnan(aD));
cG    = cG(~isnan(cG));
cD    = cD(~isnan(cD));
fvals = fvals(~isnan(fvals));
kFs   = kFs(~isnan(kFs));
kRs   = kRs(~isnan(kRs));
aGs   = aGs(~isnan(aGs));
aDs   = aDs(~isnan(aDs));
cGs   = cGs(~isnan(cGs));
cDs   = cDs(~isnan(cDs));

% Filter out bad fits
filter = 0.20;
kF    = kF(fvals < filter);
kR    = kR(fvals < filter);
aG    = aG(fvals < filter);
aD    = aD(fvals < filter);
cG    = cG(fvals < filter);
cD    = cD(fvals < filter);
kFs   = kFs(fvals < filter);
kRs   = kRs(fvals < filter);
aGs   = aGs(fvals < filter);
aDs   = aDs(fvals < filter);
cGs   = cGs(fvals < filter);
cDs   = cDs(fvals < filter);
fvals = fvals(fvals < filter);

% Calculate affinity
kD  = kR ./ kF;
kDs = kRs ./ kFs;

% Define plot bounds
plotLB = [LB(5) LB(6) (LB(6)/UB(5))];
plotUB = [UB(5) UB(6) (UB(6)/LB(5))];

% Threshold objective function value for color map
T = 0.155;

% Histogram of kD and objective function values
parHistogram9P([log10(kD) fvals], 0.1, T);
title('Distribution of log_{10}(k_D) values')

figure
histogram(fvals)
title('Distribution of objective function values')

RGB = [56  113 237
       224 25  76
       247 193 14];

%%%%%%%%%%%%%%%%%%%%%%%%%%%%%%%%%%%%%%%%%%%%%%%%%%%%%%%%%%%%%%%%%%%%%
% Plot kF, kR, and kD as a function of the objective function value %
%%%%%%%%%%%%%%%%%%%%%%%%%%%%%%%%%%%%%%%%%%%%%%%%%%%%%%%%%%%%%%%%%%%%%

% kF
figure
subplot(2,3,1)
for i = 1:length(kF)
    kFsi = kFs(i);
    kFi  = kF(i);
    Fi   = fvals(i);
    P = (log10(kFsi)-log10(plotLB(1)))/(log10(plotUB(1))-log10(plotLB(1)));
    R = RGB(1,1) + (P * (RGB(2,1) - RGB(1,1)));
    G = RGB(1,2) + (P * (RGB(2,2) - RGB(1,2)));
    B = RGB(1,3) + (P * (RGB(2,3) - RGB(1,3)));
    semilogy(Fi, kFi, 'o',...
             'markerfacecolor', ([R G B]./255),...
             'markersize', 8,...
             'markeredgecolor', ([0 0 0]./255))
    hold on
end
grid on
ylim([plotLB(1) plotUB(1)])
xlabel('Objective Function Value')
ylabel('k_F value')
title(['k_F values from parallel optimization (n = ',...
      num2str(length(kF)),...
      ')'])

% kR
subplot(2,3,2)
for i = 1:length(kR)
    kRsi = kRs(i);
    kRi  = kR(i);
    Fi   = fvals(i);
    P = (log10(kRsi)-log10(plotLB(2)))/(log10(plotUB(2))-log10(plotLB(2)));
    R = RGB(1,1) + (P * (RGB(2,1) - RGB(1,1)));
    G = RGB(1,2) + (P * (RGB(2,2) - RGB(1,2)));
    B = RGB(1,3) + (P * (RGB(2,3) - RGB(1,3)));
    semilogy(Fi, kRi, 'o',...
             'markerfacecolor', ([R G B]./255),...
             'markersize', 8,...
             'markeredgecolor', ([0 0 0]./255))
    hold on
end
grid on
ylim([plotLB(2) plotUB(2)])
xlabel('Objective Function Value')
ylabel('k_R value')
title(['k_R values from parallel optimization (n = ',...
      num2str(length(kF)),...
      ')'])

% kD
subplot(2,3,3)
for i = 1:length(kD)
    kDsi = kDs(i);
    kDi  = kD(i);
    Fi   = fvals(i);
    P = (log10(kDsi)-log10(plotLB(3)))/(log10(plotUB(3))-log10(plotLB(3)));
    R = RGB(1,1) + (P * (RGB(2,1) - RGB(1,1)));
    G = RGB(1,2) + (P * (RGB(2,2) - RGB(1,2)));
    B = RGB(1,3) + (P * (RGB(2,3) - RGB(1,3)));
    semilogy(Fi, kDi, 'o',...
             'markerfacecolor', ([R G B]./255),...
             'markersize', 8,...
             'markeredgecolor', ([0 0 0]./255))
    hold on
end
grid on
ylim([plotLB(2)/plotUB(1) plotUB(2)/plotLB(1)])
xlabel('Objective Function Value')
ylabel('k_D value')
title(['k_D values from parallel optimization (n = ',...
      num2str(length(kF)),...
      ')'])

%%%%%%%%%%%%%%%%%%%%%%%%%%%%%%%%%%%%%%%%%%%%%%%%%%
% Plot starting and ending kF, kR, and kD values %
%%%%%%%%%%%%%%%%%%%%%%%%%%%%%%%%%%%%%%%%%%%%%%%%%%

% Determine the color map for objective function values
fRGB = zeros(length(fvals), 3);

for i = 1:length(fvals)
    Fi   = fvals(i);
    if Fi < T
        P = (Fi - min(fvals)) / (T - min(fvals));
        R = RGB(3,1) - (P * (RGB(3,1) - RGB(2,1)));
        G = RGB(3,2) - (P * (RGB(3,2) - RGB(2,2)));
        B = RGB(3,3) - (P * (RGB(3,3) - RGB(2,3)));
    else
        P = (Fi - T) / (max(fvals) - T);
        R = RGB(2,1) - (P * (RGB(2,1) - RGB(1,1)));
        G = RGB(2,2) - (P * (RGB(2,2) - RGB(1,2)));
        B = RGB(2,3) - (P * (RGB(2,3) - RGB(1,3)));
    end
    fRGB(i,:) = [R G B];
end

% kF
subplot(2,3,4)
for i = 1:length(kF)
    kFsi = kFs(i);
    kFi  = kF(i);
    loglog(kFsi, kFi, 'o',...
           'markerfacecolor', (fRGB(i,:)./255),...
           'markersize', 8,...
           'markeredgecolor', ([0 0 0]./255))
    hold on
end
loglog([plotLB(1) plotUB(1)], [plotLB(1) plotUB(1)],...
       'k--', 'linewidth', 2)
grid on
xlim([plotLB(1) plotUB(1)])
ylim([plotLB(1) plotUB(1)])
xlabel('Starting k_F value')
ylabel('Ending k_F value')
title(['k_F convergene (color threshold = ',num2str(T),')'])

% kR
subplot(2,3,5)
for i = 1:length(kR)
    kRsi = kRs(i);
    kRi  = kR(i);
    loglog(kRsi, kRi, 'o',...
           'markerfacecolor', (fRGB(i,:)./255),...
           'markersize', 8,...
           'markeredgecolor', ([0 0 0]./255))
    hold on
end
loglog([plotLB(2) plotUB(2)], [plotLB(2) plotUB(2)],...
       'k--', 'linewidth', 2)
grid on
xlim([plotLB(2) plotUB(2)])
ylim([plotLB(2) plotUB(2)])
xlabel('Starting k_R value')
ylabel('Ending k_R value')
title(['k_R convergene (color threshold = ',num2str(T),')'])

% kD
subplot(2,3,6)
for i = 1:length(kD)
    kDsi = kDs(i);
    kDi  = kD(i);
    loglog(kDsi, kDi, 'o',...
           'markerfacecolor', (fRGB(i,:)./255),...
           'markersize', 8,...
           'markeredgecolor', ([0 0 0]./255))
    hold on
end
grid on
loglog([plotLB(3) plotUB(3)], [plotLB(3) plotUB(3)],...
       'k--', 'linewidth', 2)
xlim([plotLB(3) plotUB(3)])
ylim([plotLB(3) plotUB(3)])
xlabel('Starting k_D value')
ylabel('Ending k_D value')
title(['k_D convergene (color threshold = ',num2str(T),')'])

% Make a unique figure of kD vs fval
figure
for i = 1:length(kD)
    kDsi = kDs(i);
    kDi  = kD(i);
    Fi   = fvals(i);
    P = (log10(kDsi)-log10(plotLB(3)))/(log10(plotUB(3))-log10(plotLB(3)));
    R = RGB(1,1) + (P * (RGB(2,1) - RGB(1,1)));
    G = RGB(1,2) + (P * (RGB(2,2) - RGB(1,2)));
    B = RGB(1,3) + (P * (RGB(2,3) - RGB(1,3)));
    semilogy(Fi, kDi, 'o',...
             'markerfacecolor', ([R G B]./255),...
             'markersize', 10,...
             'markeredgecolor', ([0 0 0]./255))
    hold on
end
grid on
ylim([plotLB(2)/plotUB(1) plotUB(2)/plotLB(1)])
xlabel('Objective Function Value')
ylabel('k_D value')
title(['k_D values from parallel optimization (n = ',...
      num2str(length(kF)),...
      ')'])

% Make a unique figure for kD start vs kD end
figure
for i = 1:length(kD)
    kDsi = kDs(i);
    kDi  = kD(i);
    loglog(kDsi, kDi, 'o',...
           'markerfacecolor', (fRGB(i,:)./255),...
           'markersize', 10,...
           'markeredgecolor', ([0 0 0]./255))
    hold on
end
grid on
loglog([plotLB(3) plotUB(3)], [plotLB(3) plotUB(3)],...
       'k--', 'linewidth', 2)
xlim([plotLB(3) plotUB(3)])
ylim([plotLB(3) plotUB(3)])
xlabel('Starting k_D value')
ylabel('Ending k_D value')
title(['k_D convergene (color threshold = ',num2str(T),')'])

plotdata = [kFs kF kRs kR kDs kD fvals fRGB];
writematrix(plotdata, 'parProcessedResults9P.csv')

%%%%%%%%%%%%%%%%%%%%%%%%%%%%%%%%%%%%%%%%%%%%%%%%%%%%%%%%%%%%%%%%%%%%%
% Make a figure with kF, kR, kD, aG, aD, cG, cD, and fval histogram %
%%%%%%%%%%%%%%%%%%%%%%%%%%%%%%%%%%%%%%%%%%%%%%%%%%%%%%%%%%%%%%%%%%%%%

figure

% Histogram
subplot(2,4,1)
histogram(fvals, 'linewidth', 2,...
                 'facecolor', [61 174 12] ./ 255,...
                 'facealpha', 1)
xlabel('Objective function value')
ylabel('Frequency')
title('Distribution of objective function values')
axis('square')

% kF
subplot(2,4,2)
for i = 1:length(kF)
    kFsi = kFs(i);
    kFi  = kF(i);
    loglog(kFsi, kFi, 'o',...
           'markerfacecolor', (fRGB(i,:)./255),...
           'markersize', 12,...
           'markeredgecolor', ([0 0 0]./255))
    hold on
end
grid on
xlim([plotLB(1) plotUB(1)])
ylim([plotLB(1) plotUB(1)])
xlabel('Starting k_F value')
ylabel('Ending k_F value')
title(['k_F convergene (color threshold = ',num2str(T),')'])
axis('square')

% kR
subplot(2,4,3)
for i = 1:length(kR)
    kRsi = kRs(i);
    kRi  = kR(i);
    loglog(kRsi, kRi, 'o',...
           'markerfacecolor', (fRGB(i,:)./255),...
           'markersize', 12,...
           'markeredgecolor', ([0 0 0]./255))
    hold on
end
grid on
xlim([plotLB(2) plotUB(2)])
ylim([plotLB(2) plotUB(2)])
xlabel('Starting k_R value')
ylabel('Ending k_R value')
title(['k_R convergene (color threshold = ',num2str(T),')'])
axis('square')

% kD
subplot(2,4,4)
for i = 1:length(kD)
    kDsi = kDs(i);
    kDi  = kD(i);
    loglog(kDsi, kDi, 'o',...
           'markerfacecolor', (fRGB(i,:)./255),...
           'markersize', 12,...
           'markeredgecolor', ([0 0 0]./255))
    hold on
end
grid on
xlim([plotLB(3) plotUB(3)])
ylim([plotLB(3) plotUB(3)])
xlabel('Starting k_D value')
ylabel('Ending k_D value')
title(['k_D convergene (color threshold = ',num2str(T),')'])
axis('square')

ACLB = [1 1 0 0];
ACUB = [500 500 183 183];

% aG
subplot(2,4,5)
for i = 1:length(aG)
    aGsi = aGs(i);
    aGi  = aG(i);
    plot(aGsi, aGi, 'o',...
         'markerfacecolor', (fRGB(i,:)./255),...
         'markersize', 12,...
         'markeredgecolor', ([0 0 0]./255))
    hold on
end
grid on
xlim([ACLB(1) ACUB(1)])
ylim([ACLB(1) ACUB(1)])
xlabel('Starting a_G value')
ylabel('Ending a_G value')
title('a_G convergene')
axis('square')

% aD
subplot(2,4,6)
for i = 1:length(aD)
    aDsi = aDs(i);
    aDi  = aD(i);
    plot(aDsi, aDi, 'o',...
         'markerfacecolor', (fRGB(i,:)./255),...
         'markersize', 12,...
         'markeredgecolor', ([0 0 0]./255))
    hold on
end
grid on
xlim([ACLB(2) ACUB(2)])
ylim([ACLB(2) ACUB(2)])
xlabel('Starting a_D value')
ylabel('Ending a_D value')
title('a_D convergene')
axis('square')

% cG
subplot(2,4,7)
for i = 1:length(cG)
    cGsi = cGs(i);
    cGi  = cG(i);
    plot(cGsi, cGi, 'o',...
         'markerfacecolor', (fRGB(i,:)./255),...
         'markersize', 12,...
         'markeredgecolor', ([0 0 0]./255))
    hold on
end
grid on
xlim([ACLB(3) ACUB(3)])
ylim([ACLB(3) ACUB(3)])
xlabel('Starting c_G value')
ylabel('Ending c_G value')
title('c_G convergene')
axis('square')

% cD
subplot(2,4,8)
for i = 1:length(cD)
    cDsi = cDs(i);
    cDi  = cD(i);
    plot(cDsi, cDi, 'o',...
         'markerfacecolor', (fRGB(i,:)./255),...
         'markersize', 12,...
         'markeredgecolor', ([0 0 0]./255))
    hold on
end
grid on
xlim([ACLB(4) ACUB(4)])
ylim([ACLB(4) ACUB(4)])
xlabel('Starting c_D value')
ylabel('Ending c_D value')
title('c_D convergene')
axis('square')

end
```
