## Supplementary data file 1 for "Mechanisms of antibiotic action shape the fitness landscapes of resistance mutations": pdCurveIntersect9P.m

```
% Find the concentration at which two pharmacodynamic curves intersect
% Colin Hemez
% 2018

% Inputs
% WTKparams = Kinetic parameters for wild-type strain
% MTKparams = Kinetic parameters for mutant strain (higher MIC)
% Sparams   = Simulation parameters (should be the same for WT and M)
% tolerance = Tolerance for concentration estimate (1e-12 is usually good)
% expTime   = Length of time (hrs) of the simulated growth rate experiment

function [C, Factor] = pdCurveIntersect9P(Kwt,...
                                          Kres,...
                                          simParams,...
                                          tolerance,...
                                          expTime)

% Unpack parameters
MM = simParams(3);
                                     
% Find the wild-type and mutant MIC
MICtol = 1e-11;
WTMIC = findMIC9P(Kwt, MICtol, 'curve');
MTMIC = findMIC9P(Kres, MICtol, 'curve');
WTMIC = WTMIC * MM;
MTMIC = MTMIC * MM;

% Display some initial information
disp(' ')
disp('pdCurveIntersect9P()...')
disp(['MICwt: ',...
      num2str(WTMIC), ' g/l; MICres: ',...
      num2str(MTMIC/WTMIC), ' x MICwt'])

% Find a lower bound for the concentration
currC = WTMIC;
T = [0 expTime] .* 3600;
folditeration = 10;

WTnet = netRateCalc9P(Kwt,simParams,1,currC,T,'curve','exact');
MTnet = netRateCalc9P(Kres,simParams,1,currC,T,'curve','exact');

while (MTnet > WTnet)
    currC = currC / folditeration;
    WTnet = netRateCalc9P(Kwt,simParams,1,currC,T,'curve','exact');
    MTnet = netRateCalc9P(Kres,simParams,1,currC,T,'curve','exact');
end

% Define upper and lower bounds
LBC = currC;
UBC = currC * folditeration;

% Use midpoint search to find the concentration
currC = (LBC + UBC) / 2;
WTnet = netRateCalc9P(Kwt,simParams,1,currC,T,'curve','exact');
MTnet = netRateCalc9P(Kres,simParams,1,currC,T,'curve','exact');

iterationcount = 0;
iterationlim = 50;

while abs(WTnet - MTnet) > tolerance
    
    iterationcount = iterationcount + 1;
    if iterationcount > iterationlim
        disp('MAXIMUM NUMBER OF ITERATIONS REACHED')
        break
    end
    
    disp(['Current concentration: ', num2str(currC), ' g/l'])
    
    if MTnet > WTnet
        UBC = currC;
    else
        LBC = currC;
    end
    
    currC = (LBC + UBC) / 2;
    
    WTnet = netRateCalc9P(Kwt, simParams, 1, currC, T, 'curve', 'exact');
    MTnet = netRateCalc9P(Kres, simParams, 1, currC, T, 'curve', 'exact');
end

disp(' ')

C = currC;
Factor = currC / WTMIC;
    
end
```
