## Supplementary data file 1 for "Mechanisms of antibiotic action shape the fitness landscapes of resistance mutations": pdCurveMSW9P.m

```
% Find the selective advantage of a mutant strain within MSW

% Colin Hemez
% 2019

function [concFactors,netRateDiff] = pdCurveMSW9P(Kwt,...
                                                  Kres,...
                                                  simParams,...
                                                  resolution,...
                                                  expTime)

% Find the MIC of the wild-type and resistant strains
MICwt = findMIC9P(Kwt, 1e-12, 'curve');
MICres = findMIC9P(Kres, 1e-12, 'curve');
MICwt = MICwt * simParams(3);
MICres = MICres * simParams(3);

% Find the concentration at which the PD curves intersect
concTol = 1e-12;
Cint = pdCurveIntersect9P(Kwt, Kres, simParams, concTol, expTime);

% Generate the concentration range vector
concVals = logspace(log10(Cint), log10(MICres), resolution);
concFactors = concVals ./ MICwt;

% Calculate the PD curves
Binit = 1;
T = [0 expTime] .* 3600;
PDwt = netRateCalc9P(Kwt,simParams,Binit,concVals,T,'curve','exact');
PDres = netRateCalc9P(Kres,simParams,Binit,concVals,T,'curve','exact');

% Replace all negative values in wild-type PD curve with zeros
PDwtpos = PDwt(PDwt > 0);
PDwtnumneg = length(PDwt(PDwt <= 0));
PDwtzero = zeros(PDwtnumneg, 1);
PDwt = [PDwtpos ; PDwtzero];

% Take the difference between wt and res PD curves
netRateDiff = PDres - PDwt;

end
```
