## Supplementary data file 1 for "Mechanisms of antibiotic action shape the fitness landscapes of resistance mutations": pdCurves1.m

```
% Pharmacodynamic curves for different drug action schemes
% Colin Hemez
% 2018

clear; close all;

[K,S] = getParams8('ecoli','cip','tube');

% Mass of drug
MM = S(3);

% Define parameters and drug action modes to model
mechanism = {'KF', 'KR'};
drug = {'nstat', 'ncide', 'cstat', 'ccide', 'cipro'};

% Fold change in MIC for mutants, fitness hit of mutants
Mfold = 10;
hit = 0.75; % percentage of wild-type r0

% Wild-type and mutant parameter sets
WTK = K;
MTK = K;
MTK(1) = MTK(1) * hit;

% Timecourse for simulations
T = [0 18] .* 3600;

% Define concentration range to test (as a multiple of WTMIC)
minC = 0.01;
maxC = 100;
res = 5;
Crange = logspace(log10(minC), log10(maxC), res);

% Find wild-type MICs
WTMIC = zeros(length(drug),1);
tol = 1e-10;
for i = 1:length(drug)
    mode = drug{i};
    MIC = findMIC8(K, tol, mode);
    WTMIC(i) = MIC;
end
WTMIC = WTMIC .* MM;

% Calculate net growth rates for wild-type strains
disp(' ')
disp('Calculating net growth rates for wild-type strains...')

GWT = zeros(length(drug), res);

for j = 1:length(drug)
    mode = drug{j};
    C = Crange .* WTMIC(j);
    Grates = netRateCalc(K, S, 1, C, T, mode);
    GWT(j,:) = Grates;
end

% Determine fold changes in parameters needed
Pfold = zeros(length(drug), length(mechanism));
Pexps = Pfold;

Pfold = [10.0006   10.3005
         10.0004   12.2125
         10.0001   10.1750
          9.9629   16.4023
          9.9834   11.4658];
Pexps = [-1     1
         -1     1
         -1     1
         -1     1
         -1     1];
     
recalculate = 1;
if recalculate
    
    for i = 1:length(drug)
        mode = drug{i};

        for j = 1:length(mechanism)
            param = mechanism{j};

            disp(' ')
            disp('Searching for fold change...')
            disp(['Mode: ',upper(mode),'; Parameter: ',upper(param)])
            
            [F,E] = foldMod8(WTK, MTK, mode, Mfold, param);

            Pfold(i,j) = F;
            Pexps(i,j) = E;
            
        end
    end
end

% Verify mutant MICs
MTMIC = WTMIC .* Mfold;

% Calculate net growth rates for fold changes in kF and kR
disp(' ')
disp('Calculating net growth rates for mutant strains...')

GM1 = zeros(length(drug), res);
GM2 = zeros(length(drug), res);

set1ind = 2;
set2ind = 3;

fold1 = Pfold(:,1);
fold2 = Pfold(:,2);
exps1 = Pexps(:,1);
exps2 = Pexps(:,2);

for i = 1:length(drug)
    KM1 = K;
    KM2 = K;
    
    KM1(1) = KM1(1) * hit;
    KM2(1) = KM2(1) * hit;
    
    KM1(set1ind) = K(set1ind) * (fold1(i)^exps1(i));
    KM2(set2ind) = K(set2ind) * (fold2(i)^exps2(i));
    
    mode = drug{i};
    
    C = Crange .* WTMIC(i);
    
    Grates1 = netRateCalc(KM1, S, 1, C, T, mode);
    Grates2 = netRateCalc(KM2, S, 1, C, T, mode);
    
    GM1(i,:) = Grates1;
    GM2(i,:) = Grates2;
end

% Define color scheme (one color per drug action mode)
map = [26  108 239
       41  183 9  
       249 217 34 
       229 42  18 
       139 9   183];
map = map ./ 255;
map = map(1:length(drug), :);

% Plot first parameter PD curves
disp(' ')
disp('Plotting pharmacodynamic curves...')

figure

ymin = -K(5)*1.1;
ymax = K(1)*1.5;

semilogx([minC maxC], [0 0], 'k--', 'linewidth', 2)
hold on
semilogx([1 1], [ymin ymax], 'k--', 'linewidth', 2)
semilogx([Mfold Mfold], [ymin ymax], 'k--', 'linewidth', 2)
P = zeros(length(drug),1);
for i = 1:length(drug)
    P(i) = semilogx(Crange, GM1(i,:), 'linewidth', 3, 'color', map(i,:));
    semilogx(Crange, GWT(i,:), '--', 'linewidth', 3, 'color', map(i,:));
end

semilogx([minC maxC], [0 0], 'k--', 'linewidth', 2)
legend(P,upper(drug),'location','southwest')
ylim([ymin ymax])
title(['PD Curves (Fitness hit: ',...
       num2str((1-hit)*100),...
       '%; Parameter: ',...
       mechanism{1},...
       ')'])
xlabel('[C]_{DRUG} / MIC_{WT}')
ylabel('Net growth rate (1/sec)')
grid on


% Plot second parameter PD curves
figure

semilogx([minC maxC], [0 0], 'k--', 'linewidth', 2)
hold on
semilogx([1 1], [ymin ymax], 'k--', 'linewidth', 2)
semilogx([Mfold Mfold], [ymin ymax], 'k--', 'linewidth', 2)
P = zeros(length(drug),1);
for i = 1:length(drug)
    P(i) = semilogx(Crange, GM2(i,:), 'linewidth', 3, 'color', map(i,:));
    semilogx(Crange, GWT(i,:), '--', 'linewidth', 3, 'color', map(i,:));
end

semilogx([minC maxC], [0 0], 'k--', 'linewidth', 2)
legend(P,upper(drug),'location','southwest')
ylim([ymin ymax])
title(['PD Curves (Fitness hit: ',...
       num2str((1-hit)*100),...
       '%; Parameter: ',...
       mechanism{2},...
       ')'])
xlabel('[C]_{DRUG} / MIC_{WT}')
ylabel('Net growth rate (1/sec)')
grid on

% Create parameter sets for the mutants
set1K = zeros(length(drug), length(K));
set2K = set1K;

for i = 1:length(drug)
    modparams1 = MTK;
    modparams2 = MTK;
    
    modparams1(set1ind) = modparams1(set1ind) * (fold1(i) ^ exps1(i));
    modparams2(set2ind) = modparams2(set2ind) * (fold2(i) ^ exps2(i));
    
    set1K(i,:) = modparams1;
    set2K(i,:) = modparams2;
end

% Find  concentrations at which mutant and wild-type PD curves intersect
intersect1 = zeros(length(drug), 1);
intersect2 = intersect1;

for i = 1:length(drug)
    mode = drug{i};
    
    MTK1 = set1K(i,:);
    MTK2 = set2K(i,:);
    
    C1 = pdCurveIntersect8(WTK, MTK1, S, 1e-12, mode);
    C2 = pdCurveIntersect8(WTK, MTK2, S, 1e-12, mode);
    
    intersect1(i) = C1;
    intersect2(i) = C2;
    
end

% Calculate growth rate differences between WT and mutant strains
disp(' ')
disp('Calculating growth rate differences for WT and mutant strains...')

points = 101;

conc1 = zeros(length(drug), points);
conc2 = conc1;
diff1 = conc1;
diff2 = conc1;
conc1fold = conc1;
conc2fold = conc2;

for i = 1:length(drug)
    mode = drug{i};
    MIC = WTMIC(i);
    
    Crange1 = logspace(log10(intersect1(i)), log10(MTMIC(i)), points);
    Crange2 = logspace(log10(intersect2(i)), log10(MTMIC(i)), points);
    
    WTrates1 = netRateCalc(K, S, 1, Crange1, T, mode);
    MTrates1 = netRateCalc(set1K(i,:), S, 1, Crange1, T, mode);
    
    WTrates2 = netRateCalc(K, S, 1, Crange2, T, mode);
    MTrates2 = netRateCalc(set2K(i,:), S, 1, Crange2, T, mode);
    
    % Calculate differences in GROWTH rates
    for j = 1:points
        if WTrates1(j) > 0
            diff1(i,j) = MTrates1(j) - WTrates1(j);
        else
            diff1(i,j) = MTrates1(j);
        end
        
        if WTrates2(j) > 0
            diff2(i,j) = MTrates2(j) - WTrates2(j);
        else
            diff2(i,j) = MTrates2(j);
        end
    end
    
    conc1(i,:) = Crange1;
    conc2(i,:) = Crange2;
    
    conc1fold(i,:) = Crange1 ./ MIC;
    conc2fold(i,:) = Crange2 ./ MIC; 
end

% Plot the growth rate differences
figure
semilogx([1 1], [0 K(1)], 'k--', 'linewidth', 2)
hold on
diffPlot = zeros(length(drug), 1);

for i = 1:length(drug)
    diffPlot(i) = semilogx(conc1fold(i,:), diff1(i,:),...
                           'linewidth', 3,...
                           'color', map(i,:));
    semilogx(conc2fold(i,:), diff2(i,:),'--',...
             'linewidth', 3,...
             'color', map(i,:));
end
plot([0.01 10], [0 0], 'k--', 'linewidth', 2)
semilogx([1 1], [0 K(1)], 'k--', 'linewidth', 2)
legend(diffPlot, upper(drug), 'location', 'northwest')
grid on
xlabel('[C]_{DRUG} / MIC_{WT}')
ylabel('Growth rate advantage of mutant (1/sec)')
title(['Growth Rate Differences between WT and Mutant (Fitness hit: ',...
       num2str((1-hit)*100),...
       ')'])
ylim([-2e-5 14e-5])
```
