## Supplementary data file 1 for "Mechanisms of antibiotic action shape the fitness landscapes of resistance mutations": pdEmax.m

```
% Emax model for pharmacodynamic action

% Colin Hemez
% 2019

% Inputs:
% H: Hill coefficient
% MIC: Minimum inhibitory concentration
% Gmax: Maximum (drug-free) growth rate
% Dmax: Minimum (drug-saturated) death rate
% C: Range of drug concentrations

function pd = pdEmax(H,MIC,Gmax,Dmax,C)

Dmax = -1 * Dmax;

Cterm = (C ./ MIC) .^ H;

pd = Gmax - (((Gmax - Dmax) .* Cterm) ./ (Cterm - (Dmax / Gmax)));

end
```
