## Supplementary data file 1 for "Mechanisms of antibiotic action shape the fitness landscapes of resistance mutations": pdFamily9P.m

```
% Make a family of PD curves for each drug mechanism

% Colin Hemez
% 2019

% Inputs:
% concvals: Drug concentration values, expressed as multiples of MICwt
% expTime:  Length of time (in hrs) of the simulated growth rate experiment
% genotype: Genotype of the strain to simulate ('wt', 'res', or 'comp')

function [netRate,P] = pdFamily9P(concvals,expTime,genotype)

% Dertermine the genotype
if strcmp(genotype, 'wt')
    genstring = 'wild-type';
elseif strcmp(genotype, 'res')
    genstring = 'resistant';
elseif strcmp(genotype, 'comp')
    genstring = 'resistant-compensated';
else
    error('GENOTYPE not recognized')
end

% Display some initial information
disp(' ')
disp('pdFamily9P()...')
disp(['Genotype: ', upper(genstring)])
disp(' ')

% Define drug mechanisms
mech = {'cipro','nstat','cstat','ncide','ccide'};

% Convert experimental time to vector of start/end values in seconds
T = [0 expTime] * 3600;

% Set aside memory and prepare for ABkinetic simulations
netRate = zeros(length(concvals), length(mech));
Binit = 1;

% Generate PD curves for each mechanism
for i = 1:length(mech)
    
    % Retreive parameters
    mechi = mech{i};
    X = getMutParams9P('ecoli', 'cipP1', 'infect9', mechi);
    Si = X.S;
    MICwt = X.MICwt;
    
    % Find the correct kinetic parameter vector
    if strcmp(genotype, 'wt')
        Ki = X.Kwt;
    elseif strcmp(genotype, 'res')
        Ki = X.Kres;
    elseif strcmp(genotype, 'comp')
        Ki = X.Kcom;
    end
    
    % Generate vector of concentration values
    C = MICwt .* concvals;
    
    % Calculate net growth rates
    netRatei = netRateCalc9P(Ki, Si, Binit, C, T, 'curve', 'exact');
    
    netRate(:,i) = netRatei;
    
end

figure
P = semilogx(concvals, netRate, 'linewidth', 3);
legend(mech)
hold on
plot([min(concvals) max(concvals)], [Ki(2) Ki(2)], '--k')
plot([min(concvals) max(concvals)], [-Ki(4) -Ki(4)], '--k')
plot([min(concvals) max(concvals)], [0 0], '--k')
grid on
xlim([min(concvals) max(concvals)])
xlabel('[DRUG] / MIC_{WT}')
ylabel('Net growth rate')

end
```
