## Supplementary data file 1 for "Mechanisms of antibiotic action shape the fitness landscapes of resistance mutations": pdFitFamily9P.m

```
% Fit Emax models to family of PD curves simulated by model

% Colin Hemez
% 2019

% Inputs
% concvals: Drug concentration values, expressed as multiples of MICwt
% expTime:  Length of time (in hrs) of the simulated growth rate experiment

% Fits are to PD curves of RESISTANT STRAINS

function [PD, H, R2, MSW] = pdFitFamily9P(concvals, expTime)

% Obtain PD curves
PD = pdFamily9P(concvals, expTime, 'res');

% Define mechanisms
mech = {'cipro','nstat','cstat','ncide','ccide'};


% Find Hill coefficients
H   = zeros(length(mech),1);
R2  = zeros(length(mech),1);
MSW = zeros(length(mech),2);

for i = 1:length(mech)
    
    % Get parameters
    mechi = mech{i};
    X = getMutParams9P('ecoli','cipP1','infect9',mechi);
    
    % Find the fit
    [Hi,Ri] = findHill9P(PD(:,i), X, concvals);
    
    % Find the size of the mutation selection window
    [~,Cmin] = pdCurveIntersect9P(X.Kwt, X.Kres, X.S, 1e-12, expTime);
    
    % Store results
    H(i)     = Hi;
    R2(i)    = Ri;
    MSW(i,1) = Cmin;
    MSW(i,2) = X.MICres / X.MICwt;
end

csvwrite('logMSWBounds.csv', log10(MSW));
csvwrite('hillCoefficients.csv', H);
csvwrite('hillFitsRsq.csv', R2);

end
```
