## Supplementary data file 1 for "Mechanisms of antibiotic action shape the fitness landscapes of resistance mutations": plotKinetic9P.m

```
% Plot experimental and simulated bacterial kill curves

% Model by
% Pia Abel zur Wiesch
% Leonid Chindelevitch
% Colin Hemez

% Implemented by
% Colin Hemez
% 2019

function [H, R2] = plotKinetic9P(experiment,concvals,timevals,K,S)

% Set up time vector for simulations
timepoints = 100;
T = linspace(min(timevals), max(timevals), timepoints);

% Simulate population curves
simB = zeros(length(concvals), timepoints);
B0 = 1;

for i = 1:length(concvals)
    C0 = concvals(i);
    Btot = ABkinetic9P(K, S, B0, C0, T, 'curve', 'exact');
    B = sum(Btot);
    simB(i,:) = B;
end

% Simulate population curves for correlation coefficient calculation
simBcorr = zeros(length(concvals), length(timevals));

for i = 1:length(concvals)
    C0 = concvals(i);
    Btot = ABkinetic9P(K, S, B0, C0, timevals, 'curve', 'exact');
    B = sum(Btot);
    simBcorr(i,:) = B;
end

% Calculate the correlation coefficient
Rmat = corrcoef(experiment,simBcorr);
R2 = Rmat(2,1);

% Put together a color scheme
color1 = [56 113 237];
color2 = [224 25 76];
Rspace = linspace(color1(1), color2(1), length(concvals));
Gspace = linspace(color1(2), color2(2), length(concvals));
Bspace = linspace(color1(3), color2(3), length(concvals));
RGBmap = [Rspace' Gspace' Bspace'];
RGBmap = RGBmap./255;

% Plot simulated curves against experimental data
H = figure;
for i = 1:length(concvals)
    semilogy(timevals./60,experiment(i,:),'--','linewidth',1,...
             'color',RGBmap(i,:))
    hold on
    semilogy(timevals./60,experiment(i,:),'o',...
             'markerfacecolor',RGBmap(i,:),...
             'markeredgecolor',RGBmap(i,:))
    semilogy(T./60,simB(i,:),'-','linewidth',3,'color',RGBmap(i,:))
end

xlabel('Time (min)')
ylabel('CFU/ml')

grid on

end
```
