## Supplementary data file 1 for "Mechanisms of antibiotic action shape the fitness landscapes of resistance mutations": popSizeChange.m

```
function change = popSizeChange(kinParams,...
                                simParams,...
                                Binit,...
                                C0,...
                                trange)
% Run kinetic simulation
step = max(trange) - min(trange);
timepoints = [min(trange) max(trange)];

netrate = zeros(length(C0), 1);

for i = 1:length(C0)
    C = C0(i);
    Bi = ABkinetic4(kinParams,simParams,Binit,C,timepoints,step);
    Bsum = sum(Bi);
    
    % Calculate net growth rate
    netrate(i) = log(Bsum(2) / Bsum(1)) / step;
end

change = netrate;

end
```
