## Supplementary data file 1 for "Mechanisms of antibiotic action shape the fitness landscapes of resistance mutations": probabilityCurve9P.m

```
% Generate probabilities of mutant emergence

% Colin Hemez
% 2019

function Pc = probabilityCurve9P(kinParams,...
                                 simParams,...
                                 N0,...
                                 mu,...
                                 C0,...
                                 Ptarget,...
                                 Pvals,...
                                 varargin)

% Package initial population size into simulation parameters vector
simParams(1) = N0;

% Package all parameters into a single vector
mapParams = [kinParams simParams mu C0];

% Determine which parameters to vary along X and Y axes of mutation map
if ischar(Ptarget)
    if strcmp(Ptarget, 'kF')
        Pt = 5;
    elseif strcmp(Ptarget, 'kR')
        Pt = 6;
    elseif strcmp(Ptarget, 'kD')
        Pt = [5 6];
    elseif strcmp(Ptarget, 'N0')
        Pt = 13;
    elseif strcmp(Ptarget, 'mu')
        Pt = 16;
    elseif strcmp(Ptarget, 'C0')
        Pt = 17;
    end
else
    Pt = Xtarget;
end

% Rewrite Pvals as a column vector
Pvals = Pvals(:);

% Define the resolution for ABkinetic simulations (100 is a good default)
res = 100;

% Calculate probabilities
probVals = zeros(length(Pvals), 1);

for i = 1:length(Pvals)
    
    % Display information on progress of calculation
    disp(['Calculating probability for parameter value ',...
          num2str(i),...
          ' of ',...
          num2str(length(Pvals))])
    
    for j = 1:length(Pt)
        mapParams(Pt(j)) = Pvals(i,j);
    end
    
    K  = mapParams(1:12);
    S  = mapParams(13:15);
    N0 = mapParams(13);
    M  = mapParams(16);
    C  = mapParams(17);
    
    [~,~,Pi] = expectedMutants9P(K,S,N0,M,C,res,varargin);
    
    probVals(i) = Pi;
    
end

Pc = probVals;

end
```
