## Supplementary data file 1 for "Mechanisms of antibiotic action shape the fitness landscapes of resistance mutations": probCompFamily9P.m

```
% Generate a family of compensation probability curves

% Colin Hemez
% 2019

function [prob,SSinfo] = probCompFamily9P(concFactors, varargin)

% See if an MICfactor was inputted
factorCalc = 0;
if length(varargin) == 1
    factorCalc = 1;
    MICfactor = varargin{1};
end

% Define the drug mechanisms
mech = {'cipro', 'nstat', 'cstat', 'ncide', 'ccide'};

% Load an initial parameter set
K = getParams9P('ecoli','cipP1','infect9','cipro');

% Set aside memory
Gnum = 1000; % Number of growth rate values to measure at steady-state
Dnum = 1000; % Number of death rate values to measure at steady-state
prob = zeros(length(concFactors), length(mech));
if factorCalc
    SSdist = zeros((K(1) + 1), length(mech));
    SSgrowth = zeros(Gnum, length(mech));
    SSdeath = zeros(Dnum, length(mech));
    SSprob = zeros(length(mech), 1);
    SStime = SSprob;
    SSmode = SSprob;
    SSrange = zeros(2, length(mech));
    Gbox = zeros(3, length(mech));
    Dbox = Gbox;
end

% Generate probabilities
Ptarget = 'C0';

for i = 1:length(mech)
    
    % Retreive parameters
    Xi = getMutParams9P('ecoli','cipP1','infect9',mech{i});
    
    % Define parameter values
    concVals = concFactors .* Xi.MICres;
    
    % Calculate probabilties and store
    Pi = probabilityCurve9P(Xi.Kres,...
                            Xi.S,...
                            Xi.MSB,...
                            Xi.M(2),...
                            Xi.MICres,...
                            Ptarget,...
                            concVals,...
                            mech{i}, 'exact');
    
    Pi = Pi(:);
    prob(:,i) = Pi;
    
    % Calculate values at steady-state and MICfactor if needed
    if factorCalc
        [~,~,Pf] = expectedMutants9P(Xi.Kres,...
                                     Xi.S,...
                                     Xi.MSB,...
                                     Xi.M(2),...
                                     (Xi.MICres * MICfactor),...
                                     100,...
                                     'curve', 'exact');
        SSprob(i) = Pf;
        
        per = 0.99; % Percentile for range of bound target numbers
        [SSi,SStimei,Ri,SSmodei] = SSDistribution(Xi.Kres,...
                                                  Xi.S,...
                                                  1,...
                                                  (Xi.MICres*MICfactor),...
                                                  per,...
                                                  'curve', 'exact');
        SSdist(:,i) = SSi;
        SStime(i) = SStimei;
        SSmode(i) = SSmodei;
        SSrange(1,i) = min(Ri);
        SSrange(2,i) = max(Ri);
        
        % Get the growth and death curves for the mechanism
        [~,~,G,D] = systemMatrices9P(Xi.Kres, 0, 0);
        
        % Determine steady-state growth and death rate distributions
        fillIndG = 1;
        fillIndD = 1;
        for j = 1:(Xi.Kres(1) + 1)
            Gj = G(j);
            Dj = D(j);
            GNj = round(SSi(j) * Gnum);
            DNj = round(SSi(j) * Dnum);
            Gvalsi(fillIndG:(fillIndG+GNj)) = Gj;
            Dvalsi(fillIndD:(fillIndD+DNj)) = Dj;
            fillIndG = fillIndG + GNj;
            fillIndD = fillIndD + DNj;
        end
        
        Gvalsi = Gvalsi(:);
        if length(Gvalsi) > Gnum
            Gvalsi = Gvalsi(1:Gnum);
        end
        
        Dvalsi = Dvalsi(:);
        if length(Dvalsi) > Dnum
            Dvalsi = Dvalsi(1:Dnum);
        end
        
        SSgrowth(1:length(Gvalsi),i) = Gvalsi;
        SSdeath(1:length(Dvalsi),i) = Dvalsi;
        
        % Find growth and death rates a mode and upper/lower percentiles
        Gbox(1,i) = G(SSrange(1,i));
        Gbox(2,i) = G(SSmode(i));
        Gbox(3,i) = G(SSrange(2,i));
        Dbox(1,i) = D(SSrange(1,i));
        Dbox(2,i) = D(SSmode(i));
        Dbox(3,i) = D(SSrange(2,i));
        
    end
    
end

if factorCalc
    SSinfo.prob = SSprob;
    SSinfo.dist = SSdist;
    SSinfo.growth = SSgrowth;
    SSinfo.growthNorm = SSgrowth ./ (Xi.Kres(2));
    SSinfo.death = SSdeath;
    SSinfo.deathNorm = SSdeath ./ (Xi.Kres(4));
    SSinfo.time = SStime;
    SSinfo.mode = SSmode;
    SSinfo.percentileRange = SSrange - 1;
    SSinfo.Gpercentiles = Gbox;
    SSinfo.Dpercentiles = Dbox;
    SSinfo.percentile = per;
else
    SSinfo = [];
end

% Plot the probability curves
figure
plot(concFactors, prob, 'linewidth', 3)
grid on
legend(upper(mech),'location', 'northeast')
xlabel('[DRUG] / MIC{RES}')
ylabel('Probability of compensation')

end
```
