## Supplementary data file 1 for "Mechanisms of antibiotic action shape the fitness landscapes of resistance mutations": randKinParams9P.m

```
% Generate random starting conditions for SA optimization

% Colin Hemez
% 2019

function randK = randKinParams9P(problem,optIndex)

rng('shuffle')

% Make sure the index variable is sorted properly
optIndex = sort(optIndex);

% Unpack needed variables from problem struct
LB = problem.paramLB;
UB = problem.paramUB;
N  = problem.kinParams(1);

% Go through the parameters to optimize and generate random start values
randK = zeros(1, length(optIndex));

for i = 1:length(optIndex)
    P = optIndex(i);
    
    if P == 5 % kF
        % Generate a starting value from a logarithmic random distribution
        A = log10(LB(P));
        B = log10(UB(P));
        randK(i) = 10.^(A + (B-A).*rand(1,1));
        
    elseif P == 6 % kR
        % Generate a starting value from a logarithmic random distribution
        A = log10(LB(P));
        B = log10(UB(P));
        randK(i) = 10.^(A + (B-A).*rand(1,1));
        
    elseif P == 7 % aG
        A = LB(P);
        B = UB(P);
        randK(i) = A + (B-A)*rand(1,1);
        if randK(i) > 50
            LB(11) = 0;
        elseif randK(i) > 30
            LB(11) = 0.2 * LB(11);
        end
        
    elseif P == 8 % aD
        A = LB(P);
        B = UB(P);
        randK(i) = A + (B-A)*rand(1,1);
        if randK(i) > 50
            UB(12) = N;
        elseif randK(i) > 30
            UB(12) = 1.2 * UB(12);
        end
        
    elseif P == 9 % bG
        randK(i) = rand(1,1);
        
    elseif P == 10 % bD
        randK(i) = rand(1,1);
        
    elseif P == 11 % cG
        A = LB(P);
        B = UB(P);
        randK(i) = A + (B-A)*rand(1,1);
        
    elseif P == 12 % cD
        A = LB(P);
        B = UB(P);
        randK(i) = A + (B-A)*rand(1,1);
        
    else % none of the above
        error('Cannot identify the parameter specified in optIndex')
        
    end
end

end
```
