## Supplementary data file 1 for "Mechanisms of antibiotic action shape the fitness landscapes of resistance mutations": randOpt1.m

```
% See if randomly chosen start conditions affect optimization

clear; close all;

Xt = load('ciproOptProblem9P.mat');
Xt = Xt.X;

% Unpack the needed parameters
strategy = Xt.strategy;

% Determine the optimization strategy
if strcmp(strategy, '8param1')
    % 8-parameter minimization: kF, kR, aG, aD, bG, bD, cG, cD
    optIndex = 5:12;
    figuremat = [4 2];
elseif strcmp(strategy, '2param1')
    % 2-parameter minimization: cG, cD
    optIndex = 11:12;
    figuremat = [2 1];
elseif strcmp(strategy, '7param1')
    % 7-parameter minimization: kF, kR, aG, aD, bG, cG, cD
    optIndex = [5 6 7 8 9 11 12];
    figuremat = [7 1];
elseif strcmp(strategy, '6param1')
    % 6-parameter minimization: kF, kR, aG, aD, cG, cD
    optIndex = [5 6 7 8 11 12];
    figuremat = [3 2];
elseif strcmp(strategy, '4param1')
    % 4-parameter minimization: aG, aD, cG, cD
    optIndex = [7 8 11 12];
    figuremat = [2 2];
elseif strcmp(strategy, '2param2')
    % 4-parameter minimization: kF, kR
    optIndex = [5 6];
    figuremat = [2 1];
end

Xt.annealingParams = [200 100 100 30];
Xi = Xt;

tests = 1000;

data = zeros(tests,29);

for i = 1:tests
    % Generate random initial starting conditions
    randK = randKinParams9P(Xt, optIndex);
    K = Xt.kinParams;
    K(optIndex) = randK;
    Xi.kinParams = K;
    
    % Optimize
    Si = ABkineticMinimize9P(Xi);
    close(Si.plothandle)
    
    % Package data
    solK = Si.paramset;
    soli = [i K solK Si.fvalue Si.pearson Si.exit Si.output.totaltime];
    data(i,:) = soli;
end
```
