## Supplementary data file 1 for "Mechanisms of antibiotic action shape the fitness landscapes of resistance mutations": rateFun.m

```
% Constrained logistic function
% Colin Hemez
% 2018

% Inputs:
% params = A 5-element vector with the following components:
%          alpha: Steepness parameter (higher values -> more switch-like)
%          Lval:  Value of logistic function at n = 0
%          Uval:  Value of logistic function at n = N
%          iC:    Threshold for killing initiation and growth inhibition
%          N:     Total number of target molecules
% X =      A vector with values of n (usually ranging from 0 to N)

% Output:
% S =      Growth or death rate as a function of number of bound targets

% When modeling growth rates, Lval > Uval
% When modeling death rates, Lval < Uval

function S = rateFun(params, X)
alpha = params(1); % Steepness parameter (1 ~ linear; 500 ~ switch-like)
Lval =  params(2); % Value when n = 0%
Uval =  params(3); % Value when n = 100%
iC =    params(4); % Threshold for switch (inflection point)
N =     params(5); % Total number of target molecules

% Convert iC and N values to percentages to avoid numerical instability
iCp = iC / N;
Np = 1;
Xp = X ./ N;

beta =  (1 + exp(alpha * iCp));
gamma = (1 + exp(alpha * (iCp - Np)));

Xn = (1 + exp(-alpha .* (Xp - iCp)));

S = (Uval - Lval) .* ((gamma)/(gamma-beta)) .* (1 - (beta./Xn)) + Lval;

end
```
