## Supplementary data file 1 for "Mechanisms of antibiotic action shape the fitness landscapes of resistance mutations": scaleFreeAnnealFcn9P.m

```
% Scale-free annealing function for simulated annealing

function newx = scaleFreeAnnealFcn9P(optimValues,problem)

% Unpack variables
currx = optimValues.x;
currx = currx';
currt = optimValues.temperature;
nvars = problem.nvar;
LB    = problem.lb;
UB    = problem.ub;

% Define range of exploration space for each parameter
range = UB - LB;
scale = 0.3;

% Generate random direction of next step for each parameter
U = rand(nvars, 1);
D = sign(U - 0.5);

% Determine size of step for each parameter to take
stepprop = scale .* sqrt(currt) ./10;
step = range .* stepprop;
step = step .* D;

% Add the step to the current parameter value, staying within bounds
X = zeros(nvars,1);
for i = 1:nvars
    X(i) = currx(i) + step(i);
    
    if X(i) > UB(i)
        diff = UB(i) - X(i);
        newstep = rand(1) * diff;
        X(i) = currx(i) + newstep;
    elseif X(i) < LB(i)
        diff = X(i) - LB(i);
        newstep = rand(1) * diff;
        X(i) = currx(i) - newstep;
    end
end

newx = X;

end
```
