## Supplementary data file 1 for "Mechanisms of antibiotic action shape the fitness landscapes of resistance mutations": SSDistribution.m

```
% Determine the steady-state distribution of bound targets

% Colin Hemez
% 2019

function [SS,T,X,M,Z] = SSDistribution(kinParams,...
                                       simParams,...
                                       Binit,...
                                       C0,...
                                       proportion,...
                                       varargin)

% Rename parameters
K = kinParams;
S = simParams;

% Find the time at which the population reaches steady-state
if isempty(varargin)
    T = findSSDecayTime(K,S,Binit,C0);
elseif length(varargin) == 1
    T = findSSDecayTime(K,S,Binit,C0,varargin{1});
else
    T = findSSDecayTime(K,S,Binit,C0,varargin{1},varargin{2});
end

% Obtain the steady-state distribution
plotter = 0;

if isempty(varargin)
    P = cellSubPop(K,S,Binit,C0,T,plotter);
elseif length(varargin) == 1
    P = cellSubPop(K,S,Binit,C0,T,plotter,varargin{1});
else
    P = cellSubPop(K,S,Binit,C0,T,plotter,varargin{1},varargin{2});
end

SS = P;

% Find the range of bound targets, centered around the mode of the
% steady-state distribution, that contains a defined percentage of the
% total population
targets = 0:1:K(1);
mode = targets(SS == max(SS));

Xrange = mode;
Prange = SS(Xrange);
Xprop = sum(Prange);

while Xprop < proportion
    Xrange = (min(Xrange)-1):1:(max(Xrange)+1);
    Prange = SS(Xrange);
    Xprop = sum(Prange);
end

X = Xrange';
M = mode;
Z = Xprop;

end
```
