## Supplementary data file 1 for "Mechanisms of antibiotic action shape the fitness landscapes of resistance mutations": subPop2.m

```
% Subpopulation ABkinetic simulation

close all;

[K,S] = getParams9P('ecoli','cipP1','tube');

MIC = findMIC9P(K,1e-12,'curve',0);

S(1) = 1e8;

MIC = MIC*S(3);
MICfactor = 3;
C0 = MIC * MICfactor;
B0 = 1;
Thrs = 3;
T = linspace(0,Thrs*3600,2000);

[Bi, R, P] = ABkinetic9P(K,S,B0,C0,T,'curve','exact');

plot(T./3600, sum(Bi), '-k', 'linewidth', 3)
grid on
xlabel('Time (hrs)')
ylabel('CFU/ml')

Btot = sum(Bi);

Bprop = zeros((K(1)+1),length(T));
for i = 1:length(T)
    Bcurr = Btot(i);
    Bpop = Bi(:,i);
    Bpropi = Bpop./Bcurr;
    Bprop(:,i) = Bpropi;
end

% Generate the pixel values for the image
blog = log10(Bprop);

% Define threshold colors
C = [35  35  35
     13  101 168
     224 13  104
     247 193 14]; % Color map
 
cutoffs = [-6 -3 -1 -0.9];
cutoffs = sort(cutoffs);

% Go pixel-by-pixel to assign a color
dim = size(blog);
bmap = zeros(dim(1), dim(2), 3);

for i = 1:dim(1)
    for j = 1:dim(2)
        P = blog(i,j);
        if P <= cutoffs(1)
            bmap(i,j,1) = C(1,1);
            bmap(i,j,2) = C(1,2);
            bmap(i,j,3) = C(1,3);
        elseif P <= cutoffs(2)
            ind = (cutoffs(1) - P) / (cutoffs(1) - cutoffs(2));
            Crange = C(2,:) - C(1,:);
            bmap(i,j,1) = C(1,1) + Crange(1)*ind;
            bmap(i,j,2) = C(1,2) + Crange(2)*ind;
            bmap(i,j,3) = C(1,3) + Crange(3)*ind;
        elseif P <= cutoffs(3)
            ind = (cutoffs(2) - P) / (cutoffs(2) - cutoffs(3));
            Crange = C(3,:) - C(2,:);
            bmap(i,j,1) = C(2,1) + Crange(1)*ind;
            bmap(i,j,2) = C(2,2) + Crange(2)*ind;
            bmap(i,j,3) = C(2,3) + Crange(3)*ind;
        elseif P <= cutoffs(4)
            ind = (cutoffs(3) - P) / (cutoffs(3) - cutoffs(4));
            Crange = C(4,:) - C(3,:);
            bmap(i,j,1) = C(3,1) + Crange(1)*ind;
            bmap(i,j,2) = C(3,2) + Crange(2)*ind;
            bmap(i,j,3) = C(3,3) + Crange(3)*ind;
        else
            bmap(i,j,1) = C(4,1);
            bmap(i,j,2) = C(4,2);
            bmap(i,j,3) = C(4,3);
        end
    end
end

% Make the image
bmap = bmap ./ 255;
image(flip(bmap))
axis off

% Make a graph showing the whole population and subpopulations
Thrs = 8;
T = linspace(0,Thrs*3600,50);

[Bil, Rl, Pl] = ABkinetic9P(K,S,B0,C0,T,'curve','exact');

figure
semilogy(T./3600,sum(Bil),'-k','linewidth',3)
hold on
for i = 1:(K(1)+1)
    if (i == 1) || (i == K(1)+1) % index i is 1+(# bound targets)
        semilogy(T./3600,Bil(i,:),'-r')
    elseif (i > 1) && (mod(i,10) == 1) % plot every 10th sub-curve
        semilogy(T./3600,Bil(i,:),'-r')
    end
end
ylim([1e0 1e9])

% Make a graph showing the whole population and cumulative subpopulations
dim = size(Bil);
cumB = zeros(dim(1), dim(2));
cumB(1,:) = Bil(1,:);
for i = 2:dim(1)
    cumB(i,:) = cumB(i-1,:) + Bil(i,:);
end

figure
hold on
plot(T./3600,sum(Bil),'-k','linewidth',3)
plotcount = 0;
plotted = zeros(19,dim(2));
for i = 1:(K(1)+1)
    if (i == 1) % index i is 1+(# bound targets)
        plot(T./3600,cumB(i,:),'-r')
        plotcount = plotcount + 1;
        plotted(plotcount,:) = cumB(i,:);
    elseif (i > 1) && (mod(i,10) == 1) % plot every 10th sub-curve
        plot(T./3600,cumB(i,:),'-r')
        plotcount = plotcount + 1;
        plotted(plotcount,:) = cumB(i,:);
    end
end

csvwrite('cumPop.csv', (([plotted ; sum(Bil)])./1e8)');
csvwrite('cumPopTime.csv', (T./3600)');

% csvwrite('Proportions.csv', [Btot ; Bprop])
% csvwrite('ProportionsLog.csv', [log10(Btot) ; log10(Bprop)])
```
