## Supplementary data file 1 for "Mechanisms of antibiotic action shape the fitness landscapes of resistance mutations": systemMatrices9P.m

```
% Linear system matrix construction for antibiotic model
% VERSION 9P, FOR PARAMETERIZATION

% Based on the model by
% Pia Abel zur Wiesch
% Leonid Chindelevitch
% Colin Hemez

% Implemented by
% Colin Hemez
% 2019

% INPUTS
% Kparams: Kinetic parameters
%          Vector of form [N G0 D0 Dn kF kR aG aD bG bD cG cD]
%          N  = number of target complexes per cell [1/cell]
%          G0 = growth rate in the absence of antibiotic [1/sec]
%          D0 = death rate in the absence of antibiotic [1/sec]
%          Dn = death rate in saturating antibiotic concentration [1/sec]
%          kF = drug-target binding rate constant [1/M/sec]
%          kR = drug-target unbinding rate constant [1/sec]
%          aG = Sharpness of growth rate function G[i]
%          aD = Sharpness of death rate function D[i]
%          bG = Bacteriostatic potency of drug at high concentrations
%          bD = Bactericidal potency of drug at high concentrations
%          cG = Inflection point for growth rate function G[i]
%          cD = Inflection point for death rate function D[i]
% plotmode (1 or 0): 1 plots the growth and death curves
%                    0 suppresses plotting
% dialog (1 or 0): 1 turns on function dialogue
%                  0 suppresses dialogue

% Makes use of the exponential gamma function (BFUN)
% to calculate binomial coefficients

% OUTPUTS
% A: kF*C multiple of system matrix
% B: Constant portion of system matrix
% G: Growth rate as a function of bound target number
% D: Death rate as a function of bound target number

% Additional functions required in directory:
% BFUN()
% RATEFUN()
% GROWTHLIMIT()
% DEATHLIMIT()

function [A,B,G,D] = systemMatrices9P(Kparams,...
                                      plotmode,...
                                      dialog)

if dialog
    disp(' ')
    disp('SystemMatricies9P()...')
end

% Unpack parameters
N  = Kparams(1);
G0 = Kparams(2);
D0 = Kparams(3);
Dn = Kparams(4);
kR = Kparams(6);
aG = Kparams(7);
aD = Kparams(8);
bG = Kparams(9);
bD = Kparams(10);
cG = Kparams(11);
cD = Kparams(12);
states = N + 1;

if dialog
    disp('Six-parameter population dynamics')
    disp(['ALPHA (growth): ', num2str(aG)])
    disp(['ALPHA (death):  ', num2str(aD)])
    disp(['BETA (growth):  ', num2str(bG)])
    disp(['BETA (death):   ', num2str(bD)])
    disp(['GAMMA (growth): ', num2str(cG)])
    disp(['GAMMA (death):  ', num2str(cD)])
end

% Allocate memory
A0 = zeros(states);
A1 = zeros(states);

% Build kF*C multiple of system matrix (A0)
for icount = 1:states
    i = icount - 1; % Actual state number
    for jcount = 1:states
        j = jcount - 1; % Actual state index
        if j == i - 1
            M = (N - (i - 1));
            A0(icount,jcount) = M;
        elseif j == i
            M = -1 * (N - i);
            A0(icount,jcount) = M;
        else
            A0(icount,jcount) = 0;
        end
    end
end
      
%%%%%%%%%%%%%%%%%%%%%%%%%%%%%%%%%%%%%%%%%%%%%%%%%%%%%%%%%%%%%
%%% BUILD GROWTH AND DEATH RATE FUNCTIONS (G[i] and D[i]) %%%
%%%%%%%%%%%%%%%%%%%%%%%%%%%%%%%%%%%%%%%%%%%%%%%%%%%%%%%%%%%%%

% Define upper and lower bounds for growth and death
RL = G0;
RU = growthLimit(bG, G0);
DL = D0;
DU = deathLimit(bD, D0, Dn);

% Package parameters
LparamsG = [aG RL RU cG N];
LparamsD = [aD DL DU cD N];

X = 0:1:N;

% Calculate values of discrete functions G[i] and D[i]
G = rateFun(LparamsG, X);
D = rateFun(LparamsD, X);

% Set any negative values of G[i] and D[i] to zero
Gneg = (G<0);
G(Gneg == 1) = 0;
Dneg = (D<0);
D(Dneg == 1) = 0;

% Show a plot of the chosen growth and death dynamics
if plotmode
    stateset = 0:N;
    figure
    hold on
    plot(stateset, G, 'b', 'linewidth', 4)
    plot(stateset, D, 'r', 'linewidth', 4)
    grid on
    xlabel('Number of bound targets')
    ylabel('Growth/Death rate')
    title('Growth and death profiles')
    legend('Growth rate', 'Death rate')
end

%%%%%%%%%%%%%%%%%%%%%%%%%
%%% CREATE THE MATRIX %%%
%%%%%%%%%%%%%%%%%%%%%%%%%

% Suppress warnings about non-exactness for binom coeff of large N
warning('off','all')
if dialog
    warning
end

% Create the matrix
for icount = 1:states
    i = icount - 1;
    for jcount = 1:states
        j = jcount - 1;

        % Situation: j = i
        if j == i
            death = D(icount);

            if i > 0
                rev = kR * i;
            else
                rev = 0;
            end

            growth = G(icount);

            % Calculate binomial coefficient
            beef = bfun(j,i);
            %%%%%%%%%%%%%%%%%%%%%%%%%%%%%%%%%%%%
            jgrowth = G(jcount);
            split = beef * jgrowth;

            M = split - death - rev - growth;
            A1(icount,jcount) = M;

        % Situation: j = i + 1
        elseif j == i + 1
            rev = kR * (i + 1);

            % Calculate binomial coefficient
            beef = bfun(j,i);
            %%%%%%%%%%%%%%%%%%%%%%%%%%%%%%%%%%%%
            jgrowth = G(jcount);
            split = beef * jgrowth;

            M = rev + split;
            A1(icount,jcount) = M;

        % Situation: i + 2 =< j =< N
        elseif (j >= (i + 2)) && (j <= N)
            % Calculate binomial coefficient
            beef = bfun(j,i);
            %%%%%%%%%%%%%%%%%%%%%%%%%%%%%%%%%%%%
            jgrowth = G(jcount);
            split = beef * jgrowth;

            M = split;
            A1(icount,jcount) = M;

        % All other cases
        else
            A1(icount,jcount) = 0;
        end
    end

end

% Reset warnings
warning('on','all')
if dialog
    warning('query','all')
end

% Return matrices
A = A0;
B = A1;

end
```
