## Supplementary data file 1 for "Mechanisms of antibiotic action shape the fitness landscapes of resistance mutations": wtResSim1.m

```
% Simulate kill-curves for wild-type and resistant populations

% Colin Hemez
% 2019

clear; close all;

X = getMutParams9P('ecoli','cipP1','infect9','cipro');

MICfactor = 3;

Tmax = 48;
res = 80;
T = linspace(0, Tmax, res)' .* 3600;

Swt = X.S;
Swt(1) = Swt(1) - X.MSB;
Sres = X.S;
Sres(1) = X.MSB;

Bwt = ABkinetic9P(X.Kwt,Swt,1,X.MICres*MICfactor,T,'curve','exact');
Bres = ABkinetic9P(X.Kres,Sres,1,X.MICres*MICfactor,T,'curve','exact');

Bwt = sum(Bwt)';
Bres = sum(Bres)';

figure
semilogy(T./3600, Bwt, 'linewidth', 3);
hold on
semilogy(T./3600, Bres, 'linewidth', 3);
grid on
ylim([1 inf])
xlim([0 Tmax])

csvwrite('selWtSim.csv', Bwt ./ X.S(1))
csvwrite('selResSim.csv', Bres ./ X.S(1))
csvwrite('selTime.csv', T ./ 3600)
```
